# An Atlas of Phosphorylation and Proteolytic Processing Events During Excitotoxic Neuronal Death Reveals New Therapeutic Opportunities

**DOI:** 10.1101/2020.06.15.151456

**Authors:** S. Sadia Ameen, Antoine Dufour, M. Iqbal Hossain, Ashfaqul Hoque, Sharelle Sturgeon, Harshal Nandurkar, Dominik Draxler, Robert Medcalf, Mohd Aizuddin Kamaruddin, Isabelle S. Lucet, Michael G. Leeming, Dazhi Liu, Amardeep Dhillon, Jet Phey Lim, Hong-Jian Zhu, Laita Bokhari, Carli Roulston, Oded Kleifeld, D. Ciccotosto Giuseppe, Nicholas A. Williamson, Ching-Seng Ang, Heung-Chin Cheng

**Author notes:** Send all correspondence to Giuseppe Ciccotosto, Nicholas Williamson, Ching-Seng Ang and Heung-Chin Cheng., (Lead Contact).

## Abstract

Excitotoxicity, a neuronal death process in neurological disorders, is initiated by over-stimulation of neuronal ionotropic glutamate receptors. The over-stimulated receptors dysregulate proteases, protein kinases and phosphatases, which in turn modify target neuronal proteins to induce cell death. To decipher this cell death mechanism, we used quantitative proteomics, phosphoproteomics and N-terminomics to identify modified proteins in excitotoxic neurons. Data, available in ProteomeXchange (identifiers: PXD019527 and PXD019211), enabled us to identify over one thousand such proteins with calpains, cathepsins and over twenty protein kinases as their major modifiers. These protein modification events can potentially perturb signalling pathways governing cell survival, synaptogenesis, axonal guidance and mRNA processing. Importantly, blocking the modification of Src protein kinase, a signalling hub in excitotoxic neurons, protected against neuronal loss *in vivo* in a rat model of neurotoxicity. Besides offering new insights into excitotoxic neuronal death mechanism, our findings suggest potential neuroprotective therapeutic targets for treating neurological disorders.

**Graphical abstract:** 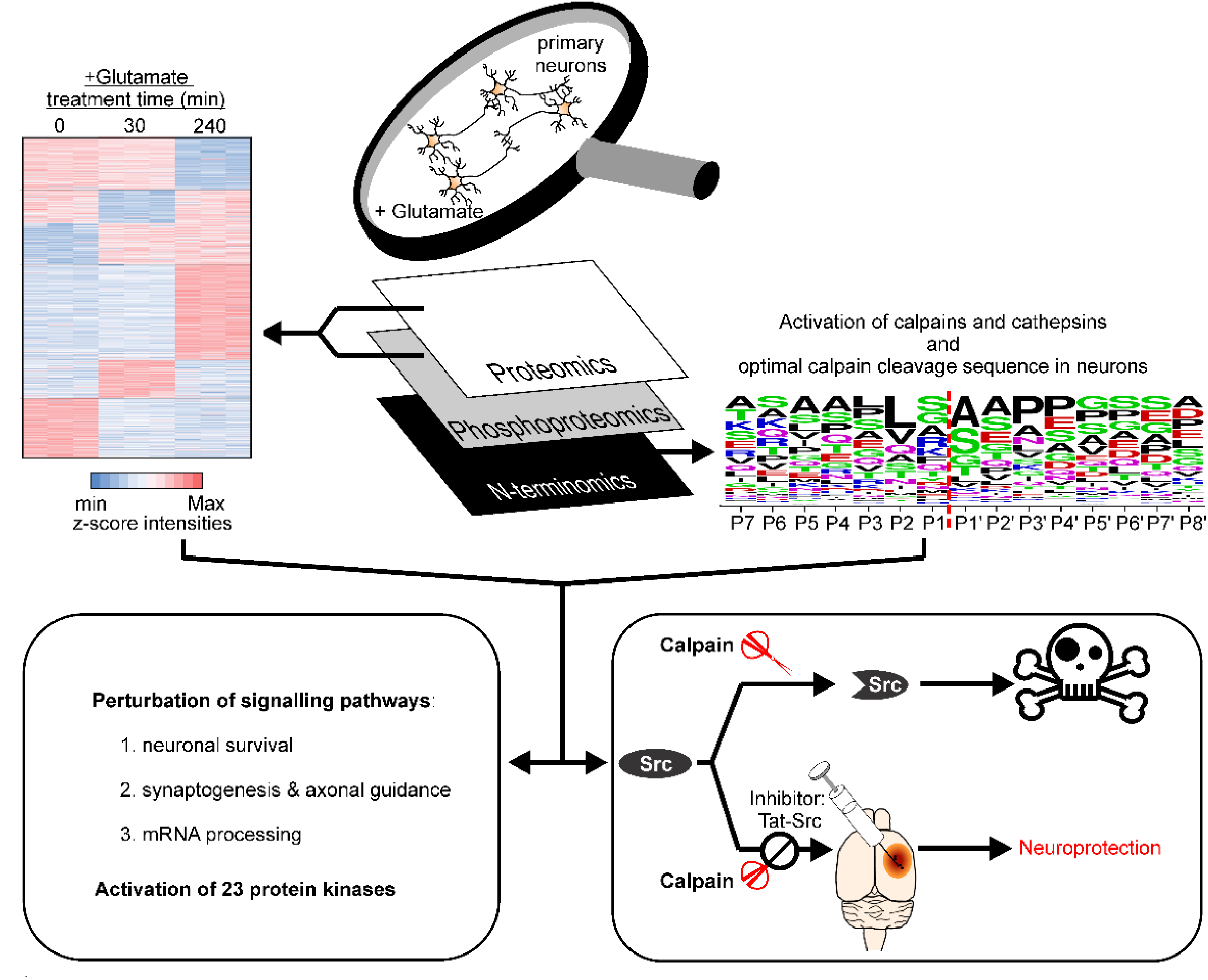

**Highlights:**

- Multi-dimensional proteomic analysis identified proteins modified by proteolysis and altered phosphorylation in neurons undergoing excitotoxic cell death.
- Calpains, cathepsins and over twenty protein kinases are major modifiers of these proteins.
- These protein modification events are predicted to impact cell survival, axonal guidance, synaptogenesis and mRNA processing.
- Blocking modification of an identified protein Src, which acts as a major signalling hub in neurons, was protective against excitotoxic injury *in vivo*.

**In Brief:** Using multidimensional proteomic approaches, Ameen, *et al*. mapped the changes of proteome, phosphoproteome and N-terminome of cultured primary neurons during excitotoxicity, a crucial neuronal death process in neurological disorders. These proteomic changes document new excitotoxicity-associated molecular events, and offer insights into how these events are organized to induce neuronal death. Potential therapeutic relevance of these molecular events is illustrated by the demonstration that *in vivo* blockade of one of these events could protect against excitotoxic neuronal loss.

## INTRODUCTION

Excitotoxicity is a pathological cell death process underpinning neuronal loss in multiple acute and chronic neurological disorders such as ischemic stroke, Parkinson’s and Alzheimer’s diseases (reviewed in (Fricker et al., 2018)). Unfortunately, there are no current FDA-approved pharmacological agents targeted to protect against excitotoxic neuronal loss in neurological disorders (O’Collins *et al*., 2006; Savitz and Fisher, 2007). This pessimistic scenario has been challenged by the promising results of two clinical trials of a cell membrane permeable peptide Nerinetide (also referred to as Tat-NR2B9c), which inhibits a key pathological cellular event directing excitotoxic neuronal death (Aarts et al., 2002; Hill et al., 2020; Hill et al., 2012).

In the first phase 2 clinical trial, Nerinetide effectively reduced the number and volume of new ischaemic strokes defined by MRI following procedurally induced ischaemic strokes in patients undergoing treatment to repair ruptured cerebral aneurysm (Hill *et al*., 2012). In a larger phase 3 clinical trial, favourable clinical outcomes of Nerinetide treatment were found in a group of patients who underwent endovascular thrombectomy for acute ischemic stroke with large vessel occlusion but not treated with the thrombolytic drug alteplase (Hill et al., 2020). Besides proving that neuroprotection to reduce brain damage in human acute ischaemic stroke is achievable, the positive clinical outcomes of Nerinetide treatment illustrate that other pathological cellular events occurring in neurons during excitotoxicity are potential therapeutic targets for the development of neuroprotective strategies to treat ischaemic stroke patients. In our endeavour to discover these potential therapeutic targets, we used a multi-dimensional quantitative proteomic approach to identify the pathological cellular events in cultured primary neurons undergoing excitotoxic cell death.

Excitotoxicity is initiated by over-stimulation of ionotropic glutamate receptors (iGluRs), especially the N-methyl-D-aspartate (NMDA) receptors (Choi, 1988; Olney, 1969; Simon et al., 1984), which permit excessive influx of extracellular calcium (Ca^2+^) into the cytosol to over-activate proteases (Ginet et al., 2014; Lankiewicz et al., 2000; Wang et al., 1996; Yamashima et al., 1998), neuronal nitric oxide synthase (nNOS) (Sattler et al., 1999) and NADPH oxidase 2 (NOX2) (Brennan et al., 2009). The excitotoxicity-activated proteases cleave specific neuronal proteins to modulate their activities, biological functions and stability (Tominaga et al., 1998; Wang et al., 1996). The activated nNOS and NOX2 catalyze over-production of reactive nitrogen species (RNS) and reactive oxygen species (ROS) to cause oxidative stress (Chan, 2001; Nakamura et al., 2013). By inhibiting interaction between the iGluRs with nNOS and NOX2, Nerinetide effectively prevents the over-stimulated iGluRs from activating nNOS and NOX2 (Brennan-Minnella et al., 2013; Sattler et al., 1999). Protein kinases and phosphatases are known mediators of excitotoxic cell death operating downstream of the excitotoxicity-activated proteases and excess ROS and RNS (Arthur et al., 2007; Dudek et al., 1997; Haass and Mandelkow, 2010; Hetman and Gozdz, 2004; Hossain et al., 2013; Xu et al., 2009). Identification of these protein kinases and phosphatases and their target substrates in neurons during excitotoxicity will help charting the signalling pathways governing excitotoxic neuronal death.

To chart the neurotoxic signalling pathways in mouse primary cortical neurons, we aim to (i) identify the substrates of the excitotoxicity-activated proteases and understand how proteolysis alters their biological functions, (ii) profile the excitotoxicity-related changes in phosphorylation of neuronal proteins and define their impacts on biological functions, and (iii) identify the upstream kinases and phosphatases controlling the excitotoxicity-related phosphorylation changes and elucidate how their activities are regulated in excitotoxicity. To achieve these aims, we used quantitative label-free global and phospho-proteomic methods (Ludwig et al., 2018) and a quantitative N-terminomic procedures called Terminal Amine Isotopic Labelling of Substrates (TAILS) (Kleifeld et al., 2010; Kleifeld et al., 2011). to document changes in stability and post-translational modification of thousands of proteins in neurons during excitotoxicity induced by glutamate over-stimulation.

We found that while glutamate treatment of neurons up to 240 min had minimal impact on neuronal protein abundance, it caused significantly altered phosphorylation states and enhanced proteolysis of close to 900 neuronal proteins. Bioinformatic analysis of these data revealed that some of the modified neuronal proteins were potential substrates of 23 protein kinases and calpain and cathepsin proteases, suggesting activation of these upstream modifying enzymes during excitotoxicity. A significant proportion of these modified neuronal proteins are key components of the signalling pathways governing mRNA processing, cell survival, synaptogenesis and axonal guidance, suggesting their dysregulation during excitotoxicity. Several modified proteins including Src, Mapk1 and Cdk1 are protein kinases operating as signalling hubs of these pathways, suggesting that their modifications during excitotoxicity contribute to excitotoxic neuronal death. This notion was supported by our demonstration that an inhibitor blocking the modification of Src could protect against excitotoxic neuronal loss *in vivo* in a rat model of neurotoxicity.

In addition to providing new information on the neurotoxic signalling pathways directing excitotoxic neuronal death, our findings also unveil new therapeutic targets such as calpain-mediated cleavage of Src kinase for the development of neuroprotective drugs that reduce excitotoxic neuronal loss in neurological disorders.

## RESULTS

### Generation of a proteomic resource for discovery of molecular events involved in glutamate-induced excitotoxic neuronal death

The signalling pathways initiated by over-stimulation of iGluRs to direct neuronal death remains poorly characterized. While these signalling pathways are likely activated at the early stage following glutamate treatment of neuronal cultures, excitotoxic cell death does not occur immediately; rather it is the prolonged activation of these pathways that causes the ultimate demise of neurons (Hoque et al., 2019; Hossain et al., 2013). Previous studies by us and others have shown that these events can be modelled in cultured rodent cortical and hippocampal neurons (Choi et al., 1987; Hossain et al., 2013; MacDermott et al., 1986). Consistent with these reports, our experiments involving timed treatment of primary neuronal cultures also demonstrated delayed cell damage and death, observed only after 240 min of glutamate treatment (Figure S1). Therefore, proteomic analysis was performed at the 30-min (early) and 240-min (late) time points to identify neuronal proteins demonstrating altered abundance or modification by proteolysis and/or phosphorylation at the early and late stages of excitotoxicity. We reasoned that identification of significantly modified neuronal proteins at these two treatment time points could unveil initiating and effector molecular events occurring during excitotoxicity. Some of these events, initiated at the early stage of excitotoxicity when neurons are still alive and sustained until late stage of excitotoxicity when neuronal death is noticeable, may represent potential drivers of excitotoxic cell death.

For global proteomic analysis, we used Data Independent Acquisition (DIA) methodology and identified 1,696 quantifiable proteins derived from neurons of all three experimental conditions (i.e. untreated (Control) neurons and neurons treated with glutamate for 30 and 240 min) (Figure 1 and Table S1). For phosphoproteomic analysis, a total of 4,700 phosphopeptides from 2,454 proteins were identified with a false discovery rate (FDR) of 1% (Figures 1A, 1B and 1C, Tables S2A and S2B). Among them, 1,676 phosphopeptides were confidently quantified using the same DIA methodology with spectral libraries constructed using two independent search engines to increase the confidence of identification (Figure S2). These phosphopeptides contained 1,954 phosphorylation sites mapped to distinct locations (termed phosphosites) in 867 neuronal proteins. These phosphosites were chosen for further analysis to identify neuronal proteins whose phosphorylation is significantly altered at 30 and 240 min after glutamate treatment. For N-terminomic analysis, a total of 5,063 and 5,552 quantifiable N-terminal peptides were identified by the TAILS method from neurons treated with glutamate for 30 min and 240 min, respectively (Kleifeld et al., 2011) (Figure 1C). They were divided into acetyl and dimethyl labelled N-terminal peptides, which for the first time, document sites of acetylation and proteolytic processing associated with the biosynthesis and cleavage of neuronal proteins proteolysed by the excitotoxicity-activated proteases.

**Figure 1.**
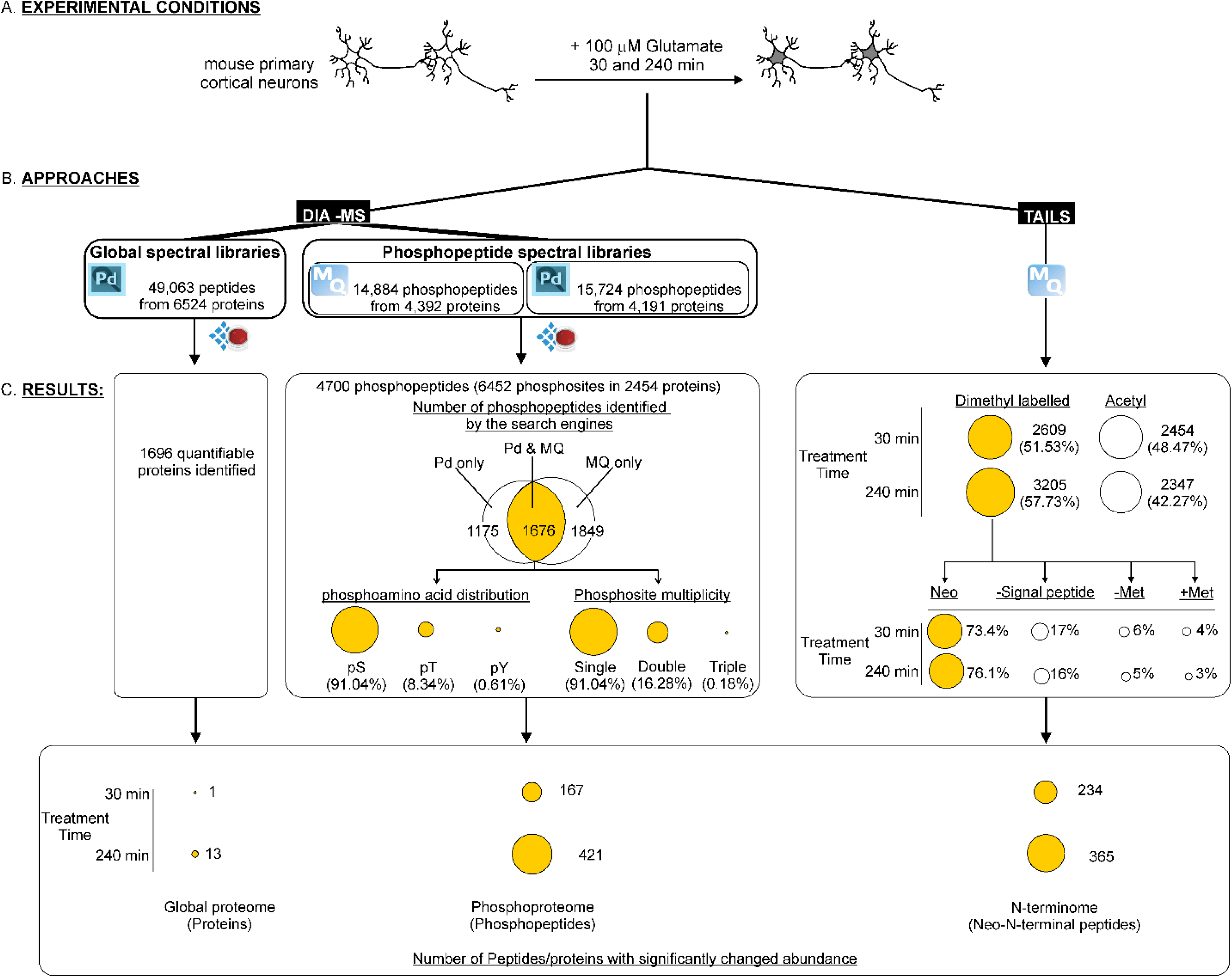
Experimental workflow and overview of results from global, phospho- and N- terminomic proteomic analysis of excitotoxic neurons. **(A)** Primary cultures of mouse cortical neurons were left untreated (Control) or treated with glutamate for 30 and 240 min to mimic early and late stages of excitotoxicity. **(B)** Analysis of the changes in global and phospho-proteomes involved two steps: (i) construction of spectral libraries with tryptic digests of cell lysates of control and glutamate-treated neurons and from tissue lysates of sham, ischemic stroke and TBI mouse brains; (ii) identification and quantification of the glutamate-induced changes in abundance of peptides and phosphopeptides from neuronal proteins with the data-independent acquisition-mass spectrometry (DIA-MS) method. Analysis of changes in the N-terminome induced by glutamate treatment was performed using the TAILS method. The search engines SEQUEST of Proteome Discover 2.1 (Pd) and Andromeda of MaxQuant (MQ) were used to identify and quantify the peptides, phosphopeptides and N-terminal peptides. Spectronaut software was used to build the spectral libraries and to analyze the DIA-MS results. **(C)** The numbers of peptides, phosphopeptides and N-terminal peptides identified and quantified in all analyses are shown. Only the phosphopeptides identified and quantified by both Pd and MQ were selected for further analysis. The identified phosphopeptides were subdivided based upon their phospho-amino acid contents. N-terminal peptides identified by the TAILS methods were classified into those derived from proteins with acetylated N-termini and those from proteins with free N-termini dimethyl labelled in the TAILS procedures (Dimethyl labelled). These two groups of N-terminal peptides were subdivided based upon other attributes including (i) the presence and absence of the methionine (-Met & +Met) encoded by the start codon for both the acetyl- and dimethyl labelled peptides, (ii) truncation that removes the N-terminal signal peptide sequence (-Signal peptide) and (iii) the presence of neo-N-termini derived from truncated protein fragments generated by internal processing of intact proteins in response to glutamate treatment (Neo). Statistical analysis was applied to identify the peptides, phosphopeptides and neo-N-terminal peptides with significantly changed abundance.

Collectively, the global proteomic, phosphoproteomic and N-terminomic changes in excitotoxic neurons defined by our multi-dimensional proteomic analysis form a comprehensive resource documenting the stability, and the types and sites of post-translational modification of thousands of neuronal proteins during excitotoxicity.

### Extensive changes in the phospho- but not global proteome of neurons during excitotoxicity

Figure S3 shows volcano plots generated by statistical analysis of changes in abundance of neuronal proteins identified in our global proteomic analysis. We chose a two-fold change in abundance ratio and a FDR of 5 % as the criteria to determine if the change in abundance of a neuronal protein was significant. Using these criteria, only 1 and 13 neuronal proteins showed significant changes in abundance at 30 and 240 min of glutamate treatment, respectively (Figure 1C and Table S1). Thus, glutamate treatment for up to 240 min had little impact on the abundance of neuronal proteins.

For phosphoproteomics, we observed a strong correlation in the abundance of phosphopeptides between biological replicates of the same treatment group (Figure S4A). However, correlation of phosphopeptide abundance between biological replicates from different treatment groups was significantly lower, indicating that glutamate treatment significantly changed the phosphorylation states of neuronal proteins. Table S2 and the volcano plots shown in Figure S4B document the distribution and significance of changes in the 1,676 phosphopeptides quantified by DIA methodology with the use of the spectral libraries. We then applied two criteria: (i) an abundance ratio > 2 or < 0.5 (i.e. log_2_ fold-change > 1 or < 1) and (ii) FDR < 1 % to define the phosphopeptides undergoing significant changes in abundance induced by glutamate treatment. With these criteria, 167 and 421 phosphopeptides at 30 and 240 min of glutamate treatment respectively, were selected as exhibiting significant changes (Figure 1C). Together, there were 483 significantly changed phosphopeptides derived from 305 phosphorylated neuronal proteins in both treatment time points (Table S3). Hence, in contrast to the little impact on the abundance of neuronal proteins, glutamate treatment for up to 240 min induced significant changes in the phosphorylation state of many more neuronal proteins.

Based on temporal changes in their abundance, the 1,676 quantified phosphopeptides were grouped into six clusters (Figure 2A). For the 483 significantly changed phosphopeptides, the temporal changes are shown (Figure 2B). Their identities and temporal changes in abundance form the basis for defining the protein kinases and phosphatases dysregulated in excitotoxic neurons.

**Figure 2.**
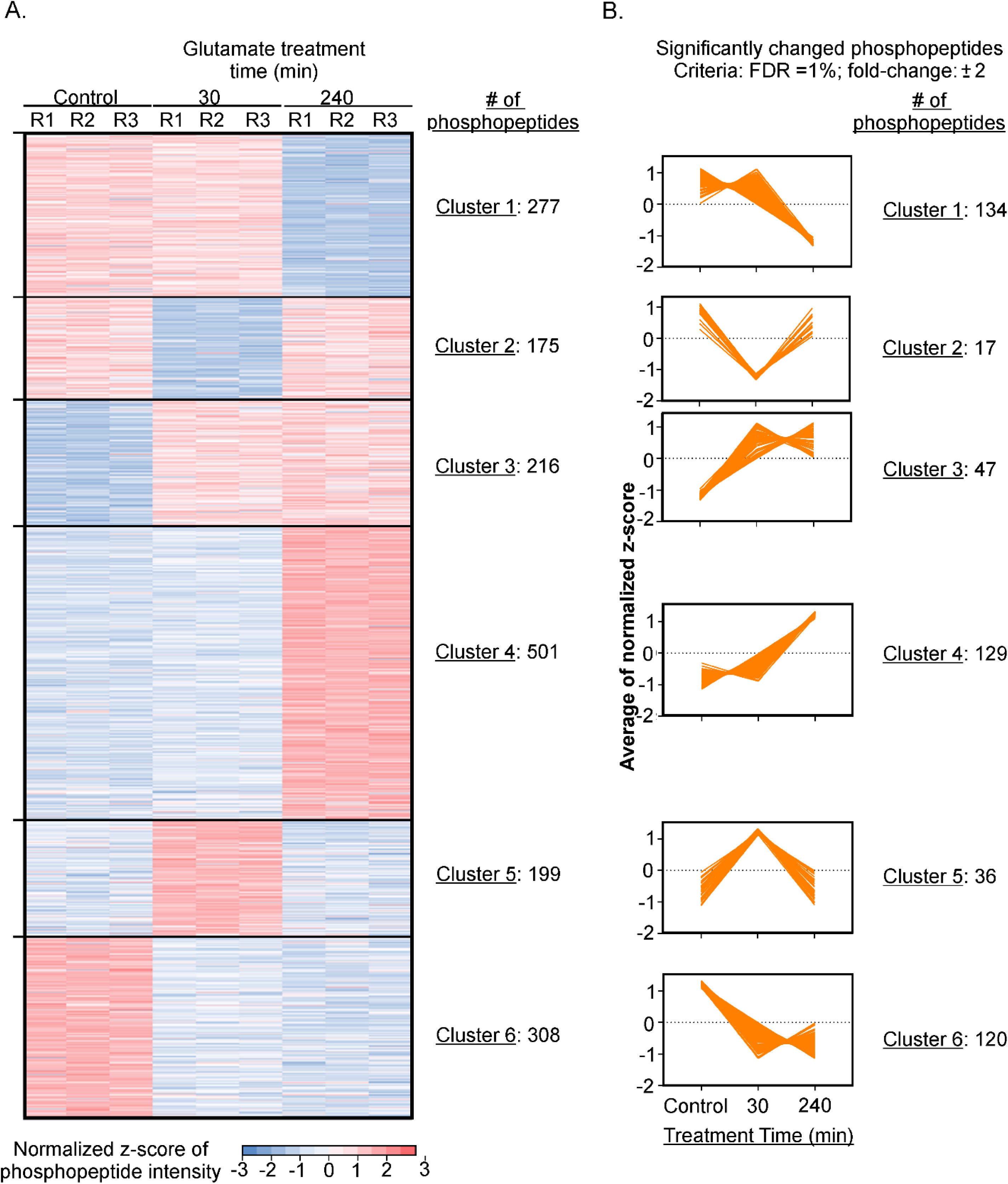
Phosphoproteomics changes of excitotoxic neurons. Heat map showing temporal changes in abundance of identified phosphopeptides. Phosphorylation states are color-coded based on the log_2_-normalized z-scores of their intensities determined from the corresponding LC-MS/MS spectra. The phosphopeptides are grouped into six clusters exhibiting different patterns of temporal changes in phosphorylation states. R1, R2 and R3: replicates 1, 2 and 3, respectively. Phosphopeptides exhibiting significant changes in abundance were identified based on a false discovery rate < 0.01 and abundance ratio > 2 or < 0.5. Plots of the normalized z-scores of the their intensities in all six clusters versus treatment time are presented. The number of phosphopeptides in each cluster is shown on the right of the plot.

### Extensive changes of the neuronal N-terminome occur during excitotoxicity

Proteolysis in cells occurs for two purposes: (i) to degrade cellular proteins into intermediate peptide fragments destined for clearance into dipeptide and amino acids (referred to as degradation in this manuscript), (ii) to process cellular proteins by removing regulatory domains or motifs to form truncated protein fragments with altered biological activities (referred to as proteolytic processing in this manuscript) (Figure S5). Degradation of cellular proteins is mediated by the proteasomal and lysosomal systems (Hofmann and Falquet, 2001; Liebl and Hoppe, 2016). In contrast, proteolytic processing is catalyzed by modulator proteases that specifically target one or several cleavage sites in accessible motifs of an intact protein to generate one or more truncated protein fragments (Sorimachi et al., 2011). As most of these protein fragments contain intact functional domain(s), they are relatively stable and may even perform specific biological functions. For example, the neuronal metabotropic glutamate receptor (mGluR1α) is cleaved by calpains during excitotoxicity to generate a stable truncated fragment that retains glutamate binding capability but lacks the ability to activate the neuroprotective PI3-Kinase/Akt signalling pathway (Xu et al., 2007).

Enhanced proteolysis of neuronal proteins is a key cellular event directing neuronal death in excitotoxicity (Brorson et al., 1995; Hossain et al., 2013; Yamashima et al., 1998). Using the TAILS method (Kleifeld et al., 2011), we identified and quantified over 5000 N-terminal peptides derived from neuronal proteins in all experimental conditions. Based upon the criteria such as the presence of an acetylated N-terminus and the methionine encoded by the start codon, these identified N-terminal peptides were classified into different groups (Figure 1C, Tables S6A and S7A, and Star Methods). Among them, we focused our further analysis on the group called neo-N-terminal peptides (Figure 1C), which contain the neo-N-terminal amino acid residue generated by proteolysis of intact neuronal proteins during excitotoxicity (Figure S5). From the identities and abundance of these neo-N-terminal peptides, we could define the identities and cleavage sites of neuronal proteins undergoing proteolysis during excitotoxicity. Based on the rationale depicted in Figure S5, neo-N-terminal peptides exhibiting increased abundance were assigned as those derived from the stable truncated protein fragments generated from enhanced proteolytic processing of neuronal proteins during excitotoxicity. Those exhibiting decreased abundance were assigned as being derived from neuronal proteins undergoing enhanced degradation for clearance during excitotoxicity.

Over 2,000 neo-N-terminal peptides were found to be generated by enhanced proteolysis during excitotoxicity (Figures 1C and 3). We then applied statistical analysis to define the thresholds for their assignment to be the N-terminal peptides generated by significantly enhanced degradation and those generated by significantly enhanced proteolytic processing of neuronal proteins during excitotoxicity (Figure 3A and Star Methods). From these thresholds, 234 and 365 neo-N-terminal peptides were found to undergo significant changes in abundance due to proteolysis of their parent neuronal proteins by the excitotoxicity-activated proteases after 30 and 240 min of glutamate treatment, respectively (Figures 1C and 3B). These peptides were further classified into those derived from proteins undergoing enhanced proteolytic processing and those derived from proteins undergoing enhanced degradation (Figure 3B and Tables S6B and S7B). Collectively, these findings reveal for the first time the identities, exact cleavage sites, stability and consequences of proteolysis of cellular proteins targeted by excitotoxicity-activated protease in neurons.

**Figure 3.**
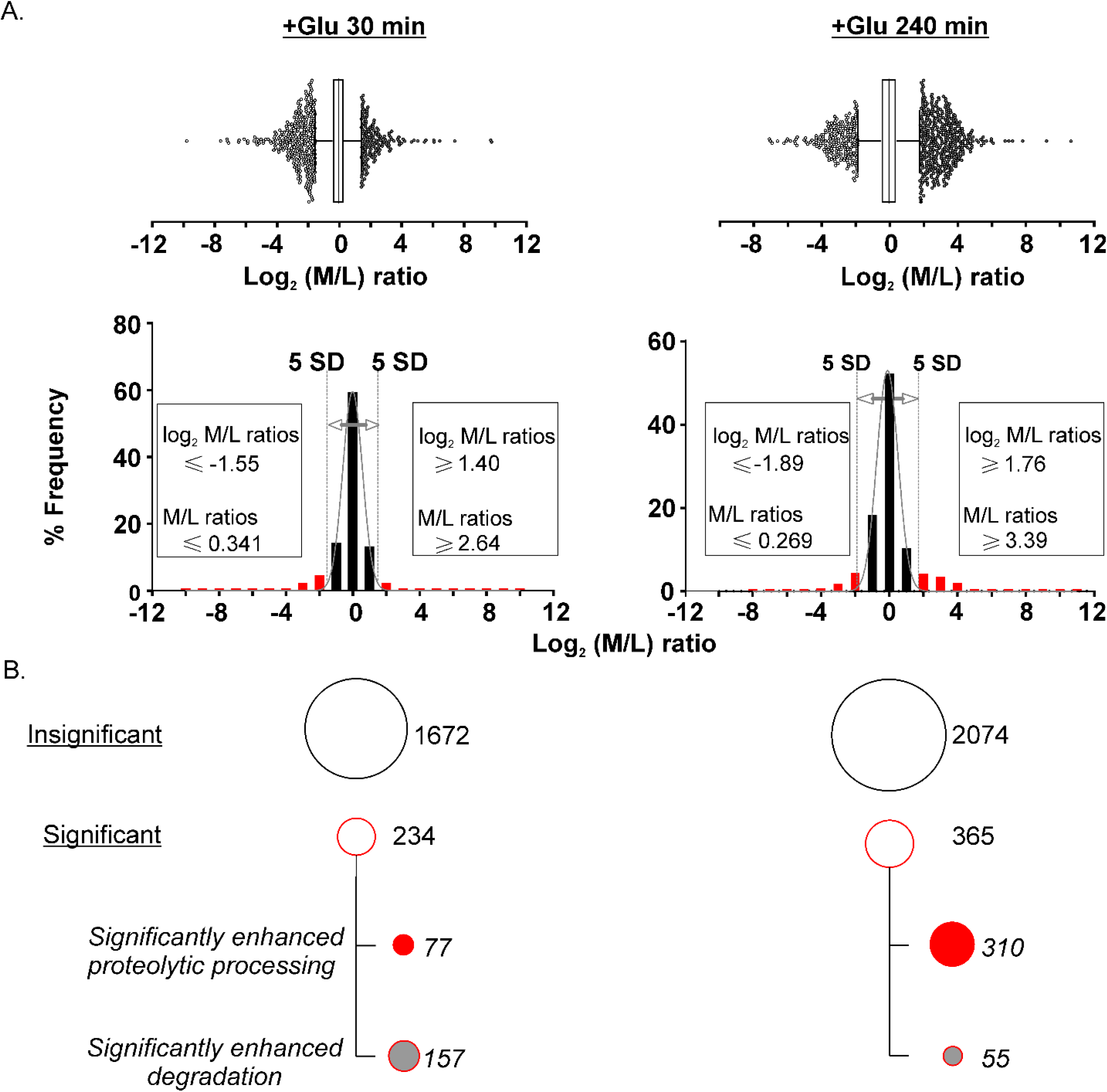
Determination of the abundance ratio (M/L) cut-off values for classification of neo-N-terminal peptides with significantly altered abundance during excitotoxicity. **(A)** Box-and-whisker plots (upper panels) of M/L ratio cut-off values determining if the neo-N-terminal peptides were generated by significantly enhanced degradation or by significantly enhanced proteolytic processing. The histograms (lower panels) show normal distribution of the data after 30 and 240 min of glutamate treatment. Insets: M/L ratio cut-off values to define if the neo-N-terminal peptides were derived from proteins undergoing significantly enhanced degradation or proteolytic processing during excitotoxicity. **(B)** Bubble plots depicting the number of neo-N-terminal peptides exhibiting significant changes in abundance (Significant) and those without significant changes in abundance (Insignificant) in excitotoxicity determined by the M/L cut-off values. The “Significant” group of peptides were subdivided into neo-N-terminal peptides derived from proteins undergoing significantly enhanced proteolytic processing and those derived from proteins undergoing significantly enhanced degradation.

### Phosphoproteomics and N-terminomics profiling revealed perturbation of synaptogenesis, cell-cell communications, cell survival and mRNA processing in excitotoxic neurons

Previous biochemical and *in vivo* studies of neurons and non-neuronal cells have defined the functions and signal transmission mechanism of a number of canonical signalling pathways [reviewed in (Kramer et al., 2014)]. Based upon the phosphopeptides and neo-N-terminal peptides exhibiting significantly changed abundance induced by glutamate treatment listed in Tables S3, S4B and S5B, we identified the proteins undergoing phosphorylation, dephosphorylation and/or enhanced proteolytic processing in excitotoxic neurons. We then used the Ingenuity Pathway Analysis (IPA) software to interrogate these neuronal proteins to identify canonical signalling pathways perturbed in neurons during excitotoxicity. This analysis revealed that signalling pathways governing synaptogenesis and axonal guidance, cell junction and cell-cell interactions and cell survival were the most perturbed neuronal pathways during excitotoxicity (Figures S6A and S7). Significant perturbation of these pathways was observed after 30 min of glutamate treatment and sustained over 240 min of treatment. These findings suggest that excitotoxic neuronal death involves perturbation of these pathways. The IPA software also predicted how these neuronal proteins form interaction networks. Figure S6B shows that the five top-ranked predicted networks perform the cellular functions of mRNA processing and regulation of organization of cytoplasm and cytoskeleton.

The site at which phosphorylation occurs is a major determinant of its effect on the function of a protein. For example, autophosphorylation of the Src tyrosine kinase at Y419 leads to activation, while phosphorylation of Y530 causes its inhibition (Boczek et al., 2019; Okada et al., 1991). We reasoned that the analysis of phosphosites in neuronal proteins that significantly changed during excitotoxicity could provide information on the impact of specific phosphorylation events on intracellular signalling of excitotoxic neurons. As the predictive algorithm of the IPA core analysis software does not consider the phosphosites on proteins, we used the PhosphoPATH app in the cytoscape software package to interrogate the 305 phosphoproteins harbouring the 483 significantly changed phosphosites (Figure 2B) against the databases documenting previously discovered *in vitro* protein-protein interactions and phosphosite-specific functional interactions among cellular proteins in neurons and other cell types (Raaijmakers et al., 2015). Figure 4A shows the predicted interaction networks and heat maps of the changes in phosphorylation levels of the significantly changed phosphosites. The PhosphoPATH analysis predicted that phosphosite-enriched interactions among the neuronal proteins shown in the networks are mostly involved in regulating the signalling pathways governing mRNA processing and focal adhesion/PI3K/Akt/mTOR signalling critical to neuritogenesis, cytoskeletal organization and cell survival (Figure S6C). Thus, in agreement with the predictions by the IPA software, the PhosphoPATH app predicted significant perturbation of signalling pathways governing mRNA processing, synaptogenesis and neuronal survival in excitotoxic neurons. In these networks (Figure 4A), Src, Mapk1, Cdk1, Ywhaq, Hsp90aa1, Flnc, Mcm2 and Sh3kp1 are phosphoproteins forming 6 to 15 interactions with other neuronal proteins in excitotoxic neurons, suggesting that they act as hubs of signalling by engaging in phosphosite-specific interactions with multiple neuronal phosphoproteins during excitotoxicity. Among them, Src, Mapk1 and Cdk1 are protein kinases, which can potentially phosphorylate their interacting partner neuronal proteins to modulate their functions.

**Figure 4.**
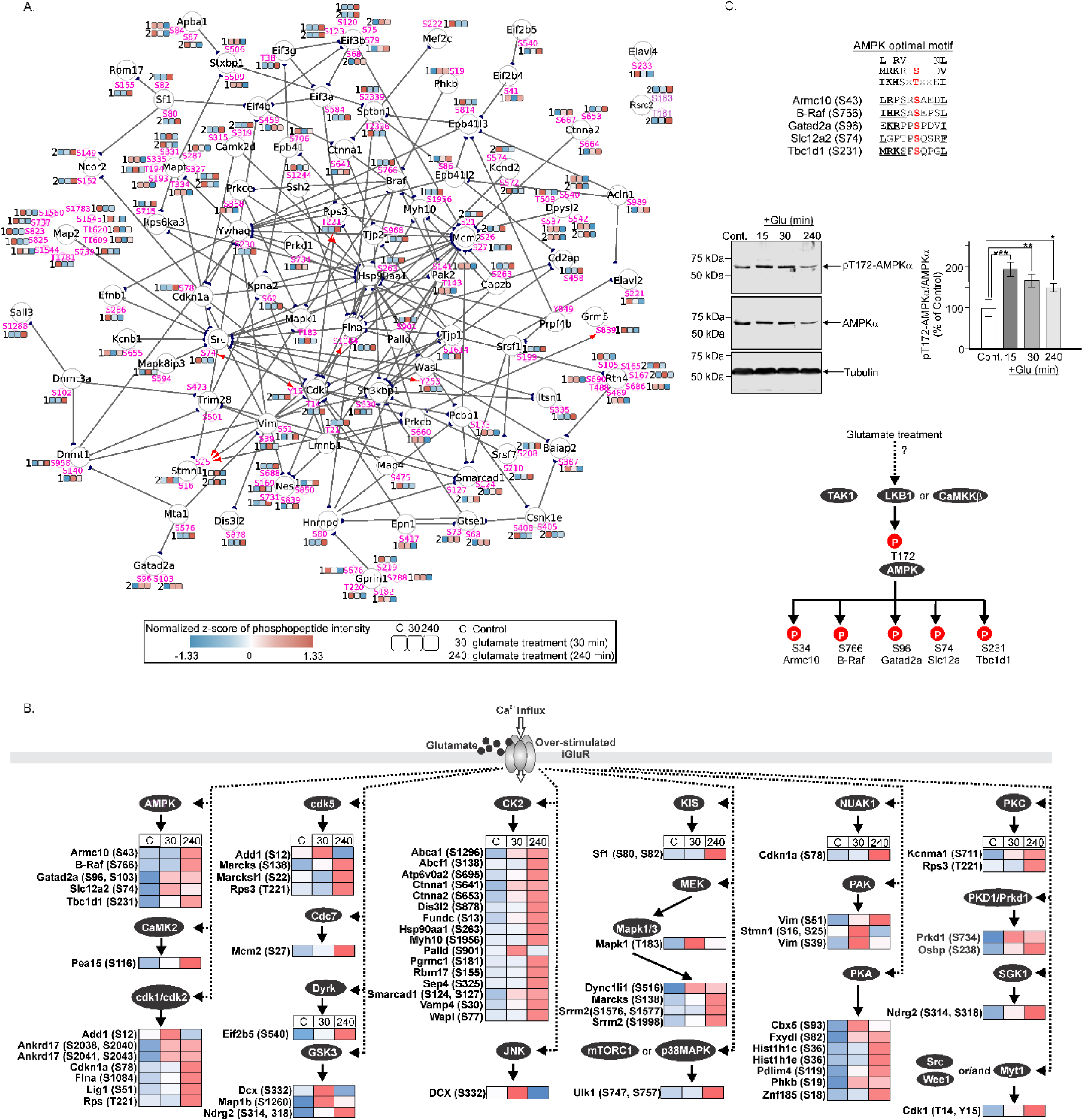
Phosphoproteome-derived interaction networks and protein kinase activity in excitotoxic neurons. **(A)** Phosphosite-specific interactions of neuronal proteins during excitotoxicity predicted by the PhosphoPATH App in Cytoscape. Quantitative information (shown as heat maps defined by the normalized z-score) of each selected phosphosites in the three experimental conditions (Control, + Glutamate 30 min and + Glutamate 240 min) are added to the network. Straight lines: protein-protein interactions defined by Biogrid; red arrowhead: kinase-substrate interactions defined by PhosphoSitePlus; filled half circle in black: the interactor protein initiating the specific interaction. Numbers in front of each heat map color bar: number of phosphosites per phosphopeptide in each quantified peptide. **(B)** Prediction of the protein kinases activated in neurons during excitotoxicity. The heat map besides each selected phosphosite shows the temporal changes in normalized z-score of the phosphopeptide intensity during excitotoxicity. Gene names and the phosphosites in the selected neuronal proteins are shown. These proteins have at least one phosphosite exhibiting increased abundance during excitotoxicity. Arrows with solid lines: the upstream protein kinases predicted to directly phosphorylate the selected phosphosites. Publications supporting the prediction are listed in Table S9. Arrows with dotted lines: Over-stimulation of the neuronal iGluRs is predicted to either activate these upstream kinases or facilitate their phosphorylation of the corresponding phosphosites. S12 of Add1: cdk1/cdk2 and cdk5 are the predicted upstream kinases; T221 of RPS3: PKC and cdk5 are the predicted upstream kinases. S732 of PKD1/Prkd1: its autophosphorylation leads to autoactivation. Insets: experimental conditions and the scales of phosphopeptide intensity shown in the heatmap. **(C)** Validation of predicted activation of AMPK during excitotoxicity. *Upper panel*: Red bold fonts represent the phosphosites. Major determinants for optimal phosphorylation are presented in bold fonts. Determinants of optimal phosphorylation are presented in normal fonts. Residues in the identified neuronal proteins showing conformity with residues in the AMPK optimal motif are underlined. *Middle panel*: Western blots showing the immunoreactive signals of phospho-T172 relative to total AMPKα level in untreated (Control) neurons and neurons at 15, 30 and 240 min of glutamate treatment and densitometric analysis (results are presented as mean ±SD, n = 3; * indicates p<0.05, ** indicates p<0.01 and *** indicates p<0.001; one-way ANOVA with Dunnett’s multiple comparison test). *Lower panel*: A model of the mechanism of AMPK activation during excitotoxicity. Glutamate treatment activates TAK1, LKB1 or CaMKKβ, which in turn activates AMPK by phosphorylation of T172 its α-subunit.

In summary, the interaction networks predicted by IPA and PhosphoPATH app provide valuable clues of how proteins undergoing significant proteolysis and/or changes in phosphorylation interplay to perturb mRNA processing, synaptogenesis and cell survival in excitotoxic neurons (Figures 4A, S6 and S7). Despite these predictions, the networks cannot fully reveal the neurotoxic signalling pathways directing excitotoxic neuronal death as they were only based on known information derived from previous studies of other systems and we did not consider other new findings such as the exact sites of cleavage in the proteolysed neuronal proteins generated in the present study.

### Phosphoproteome-derived kinase activities in excitotoxic neurons

One way to chart the signalling pathways associated with excitotoxic neuronal death is to predict upstream protein kinases and phosphatases directly phosphorylating and/or dephosphorylating the significantly changed phosphosites that we identified. While much is known about the molecular basis of substrate recognition of protein kinases, relatively little is known about the molecular basis of phosphosite-specific dephosphorylation of cellular proteins by phosphatases (Li et al., 2013; Miller and Turk, 2018). As such, we focused our attention on predicting which upstream protein kinases phosphorylate the significantly changed phosphosites (Figure 2B). Of the six clusters of the significantly changed phosphosites shown in Figure 2B, the increased abundance of phosphosites in clusters 3, 4 and 5 (Figure 2) was likely caused by activation of their upstream protein kinases and/or inactivation of the upstream phosphatases in neurons during excitotoxicity. To predict the upstream kinases of these phosphosites, we compared sequences of the phosphosites in clusters 3, 4 and 5 (Figure 2B; Table S3) against the optimal phosphorylation sequences of known protein kinases. We then performed a literature search of public repositories such as Signor (Perfetto et al., 2016), PubMed and Phosphositeplus (Hornbeck et al., 2015) to ascertain if the selected phosphosites were previously found to be phosphorylated by the predicted kinases *in vitro* and/or in cells. As many protein kinases co-localize with their protein substrates to form protein complexes, our analysis also involves interrogating whether the phosphoproteins were previously found to form protein complexes with the predicted upstream protein kinases in cells and/or *in vivo*. For example, glutamate treatment induced a significant increase in abundance of the phospho-T183 phosphopeptide derived from the Mapk1 (Figure 4B). Since MEK forms protein complexes with Mapk1 and directly phosphorylates T183 of Mapk1 (Xue et al., 2014; Yu et al., 1998), the increased abundance of this phosphopeptide suggests activation of neuronal MEK during excitotoxicity. Results of this analysis are shown in Figure 4B, Table S3 and Table S10.

Figure 4B shows heat maps of changes in the phosphorylation levels of selected phosphosites and their predicted upstream kinases. In our previous phosphoproteomic analysis of excitotoxic neurons, we employed similar criteria to analyse our data and predicted activation of seven neuronal protein kinases including CK2, Cdk5, GSK3, JNK, MEK, Rock1 and SGK1 during excitotoxicity (Hoque et al., 2019). Our analysis of the significantly changed phosphosites (Figure 2B) predicted activation of these seven kinases as well as sixteen additional protein kinases in neurons during excitotoxicity (Figure 4B).

### Validation of predicted activation of neuronal AMP-dependent protein kinase (AMPK) during excitotoxicity

The increased phosphorylation of neuronal Armc10, B-Raf, Gatad2c, Slc12a2 and Tbc1d1 during excitotoxicity suggests that the upstream kinase(s) targeting them were activated in excitotoxic neurons (Figure 4B). The sequences around phosphosites identified in these proteins show strong conformity to the optimal phosphorylation motif of AMPK (Figure 4C) (Schaffer et al., 2015). Furthermore, AMPK phosphorylation of these proteins at the identified phosphosites in cultured cells and/or *in vivo* has been documented (Chen et al., 2008; Chen et al., 2019; Ding et al., 2017; Fraser et al., 2014; Shen et al., 2013). Consistent with our prediction that AMPK was activated and directly phosphorylated these proteins in neurons during excitotoxicity (Figure 4B), Western blot analysis revealed increased phosphorylation of T172 in the kinase activation loop of the AMPK α-subunit (Figure 4C). Liver kinase B1 homolog (LKB1), CaM kinase kinase β (CaMKKβ) and transforming growth factor-β-activating kinase 1 (TAK1) are the known upstream kinases directly phosphorylating T172 of AMPKα to activate AMPK (Herrero-Martin et al., 2009; Woods et al., 2005; Woods et al., 2003). Our findings therefore predict activation of these kinases during excitotoxicity (Figure 4C).

### A new mechanism of neuronal CRMP2 dysregulation during excitotoxicity

Dysregulation of CRMP2, a microtubule-associated protein that regulates neuronal polarity in developing neurons (Morita and Sobue, 2009) was previously found to contribute to neuronal loss in neurodegenerative diseases (Kondo et al., 2019; Wakatsuki et al., 2011). However, the exact mechanism of CRMP2 dysregulation in neurons and the neurotoxic mechanism of the dysregulated CRMP2 remain unclear. Our TAILS data indicate that CRMP2 underwent significantly enhanced proteolytic processing of its C-terminal tail at sites (A^516^↓S^517^ and S^517^↓S^518^) in excitotoxic neurons (Figure 5A). This proteolytic processing event is expected to generate long truncated N-terminal fragments of ∼ 57 kDa in size and short truncated C-terminal fragments of ∼6 kDa. We also observed significant decrease in abundance of phosphopeptides with pT^537^, pS^540^ and pS^542^ in the glutamate-treated neurons (Figure 5A), suggesting that the short C-terminal truncated fragments were either poorly phosphorylated or possibly degraded in neurons during excitotoxicity.

**Figure 5.**
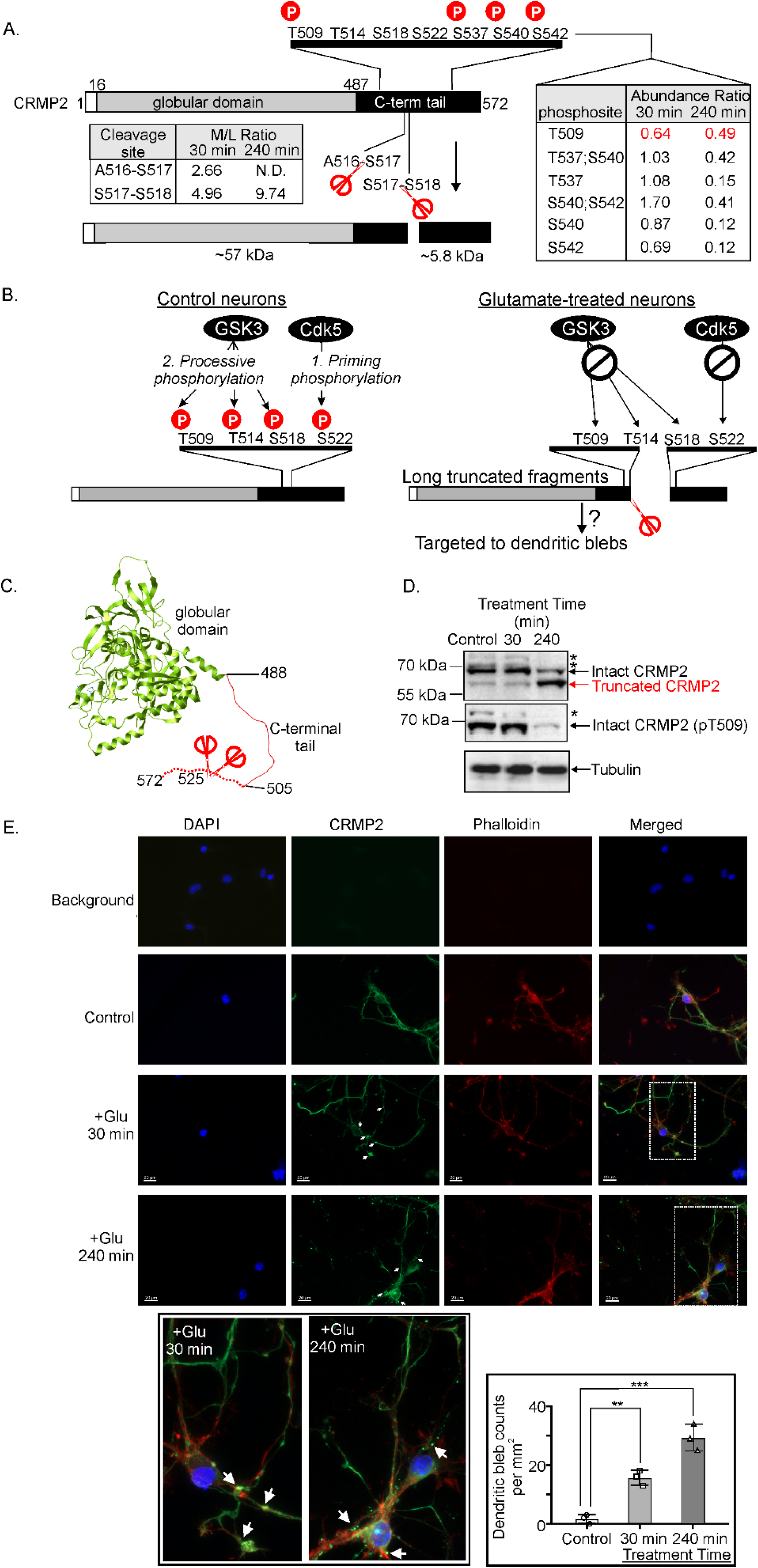
A newly discovered mechanism of dysregulation of neuronal CRMP2 during excitotoxicity. **(A)** CRMP2 is cleaved at sites near T509 in its C-terminal tail. Left inset: the abundance (M/L) ratios of the neo-N-terminal peptides at 30 and 240 min after glutamate treatment. Right inset: the abundance ratios of the identified phosphosites in the C-terminal tails of CRMP2 at 30 and 240 min after glutamate treatment. N.D.: not detected. Red scissors: cleavage sites. P in red sphere: phosphorylation. **(B)** A model depicting the new mechanism of dysregulation of neuronal CRMP2 during excitotoxicity uncovered by our proteomic findings. In control neurons, CRMP2 undergoes hierarchical phosphorylation by Cdk5 and GSK3 at sites in the C-terminal tail. Cdk5 phosphorylates the priming site S522. Upon phosphorylation, pS522 binds GSK3, which catalyses processive phosphorylation of CRMP2 at three other sites in the order of S518, T514 and T509. In excitotoxic neurons, cleavage of CRMP2 generates a long truncated CRMP2 fragment that lacks the priming site S522, abolishing S522 phosphorylation by Cdk5 and in turn suppressing processive phosphorylation of S518, T514 and T509 by GSK3. The truncation and lack of phosphorylation at T509, T514 and S518 may contribute to the accumulation of the immunoreactive CRMP2 signals at the dendritic blebs shown in panel E. **(C)** Structure of a phosphomimetic mutant of CRMP2 (PDB accession: 5yz5). Dotted line shows the disordered C-terminal tail region. **(D)** Western blots of lysates from control and glutamate-treated neurons probed with anti-CRMP2, anti-pT509 CRMP2 and tubulin antibodies. Asterisks: potential hyper-phosphorylated forms of intact CRMP2 detected by the anti-CRMP2 and anti-pT509 CRMP2 antibodies. **(E)** Fluorescence microscopy images showing actin (phalloidin), CRMP2 and nuclei (DAPI) in control and glutamate-treated neurons. White arrows indicate dendritic blebs. The close-up views of the images in the rectangles marked by white dotted lines are shown. Inset: The number of dendritic blebs per mm^2^ in control and the glutamate treated neurons in three biological replicates. **: p < 0,01; ***: p <0.001.

Besides T537, S540 and S542, the C-terminal tail of CRMP2 consists of several sites undergoing hierarchical phosphorylation by Cdk5 and GSK3β in neurons (Figure 5B, left panel). In this series of phosphorylation events, CRMP2 is first phosphorylated at S522 by Cdk5 (Uchida et al., 2005). The phospho-S522 then acts as a priming site recognized by GSK3β, which phosphorylates T509, T514 and S518 in a processive manner (Figure 5B, left panel) (Uchida et al., 2005). As cleavage at A^516^↓S^517^ and S^517^↓S^518^ removed the priming phosphorylation site S522, we predict that GSK3β could not recognize and efficiently phosphorylate the resultant long truncated CRMP2 fragments at T509, T514 and S518 (Figure 5B, right panel). Figure 5C shows that the long-truncated fragments retain the globular domain critical to promoting microtubule polymerization (Niwa et al., 2017). The microtubule polymerization-promoting activity of CRMP2 is regulated by GSK3β-mediated phosphorylation of these C-terminal tail sites (Yoshimura et al., 2005). Owing to the lack of phosphorylation at these sites, the microtubule polymerization-promoting activity of the fragments cannot be regulated by Cdk5 and GSK3β even though both kinases are known to be aberrantly activated in neurons during excitotoxicity. Indeed, results of our phosphoproteomic analysis revealed that CRMP2 phosphorylation at T509 was significantly reduced (Figure 5A). Consistent with our phosphoproteomic results, the truncated fragment CRMP2 fragments could not cross-react with the anti-pT509 CRMP2 antibody (Figure 5D). Collectively, our Western blot data validated our TAILS and phosphoproteomic findings that CRMP2 was cleaved at sites in the C-terminal tail to generate the long truncated N-terminal fragments that were notn phosphorylated by GSK3β at T509. Our data therefore support the model shown in Figure 5B that GSK3β could not efficiently phosphorylate T509 in the truncated fragments because they lacked the Cdk5-targeted priming phosphorylation site S522.

CRMP2 performs physiological functions such as promoting axonal elongation (Inagaki et al., 2001), modulating microtubule dynamics by interacting with other proteins such as tubulins and microtubules (Niwa et al., 2017; Yuasa-Kawada et al., 2003). Hence, it plays an important role in neuronal development and polarity (Yoshimura et al., 2005). Phosphorylation of S522, T509, T514 and S518 in the C-terminal tail are known to govern CRMP2 tetramer formation and impact CRMP2 interactions with GTP-tubulins (Niwa et al., 2017; Sumi et al., 2018; Wilson et al., 2014). Our findings of significantly enhanced truncation and reduced phosphorylation at T509 during excitotoxicity predict that these post-translational modifications could alter its subcellular localization in excitotoxic neurons. Consistent with this prediction, we demonstrated for the first time that glutamate treatment induced a significant accumulation of neuronal CRMP2 at specific bead-like structures on dendrites called dendritic blebs, previously known to form in neurons during excitotoxicity (Figure 5E) (Greenwood et al., 2007; Hasbani et al., 2001; Hosie et al., 2012). Moreover, the number of CRMP2-containing dendritic blebs in neurons at 240 min of glutamate treatment was significantly higher than that in neurons at 30 min of treatment (inset of Figure 5E). These findings are in agreement with the previous report that CRMP2 was cleaved by calpains in excitotoxic neurons and in rat brains after traumatic brain injury (Zhang et al., 2007). Figure 5C shows that the identified cleavage sites in CRMP2 reside at the flexible C-terminal tail, supporting the previous findings that calpains prefer to cleave disordered and flexible motifs in protein substrates (Moldoveanu et al., 2004; Sorimachi et al., 2011; Tompa et al., 2004).

In summary, our findings unveil a novel regulatory mechanism of CRMP2 during excitotoxicity as depicted in Figure 5B. In this mechanism, over-stimulation of neurons with glutamate leads to activation of a protease that cleaves CRMP2 at sites in the C-terminal tail to form long truncated N-terminal fragments that retain the microtubule-binding globular domain and short C-terminal fragments. As the long truncated fragments lack S522, which is phosphorylated by Cdk5, GSK3 is unable to bind to them and phosphorylate their T509 and T514 residues. Since CRMP2 is a crucial regulator of axonal guidance signalling (Inagaki et al., 2001), its enhanced truncation, reduced phosphorylation at T509 and T514, and accumulation in dendritic blebs can potentially contribute to injury to dendrites and synapses associated with excitotoxic neuronal death (Hasbani et al., 2001; Hosie et al., 2012).

### Processing of neuronal doublecortin-like kinase 1 (DCLK1) during excitotoxicity generates truncated fragments with intact functional domains

DCLK1, a member of the doublecortin (DCX) family (Gleeson et al., 1998) is a microtubule-associated protein (MAP), which plays a critical role in regulating microtubule assembly. DCLK1 is a bifunctional protein consisting of two DCX domains and a C-terminal serine/threonine kinase domain expressed in both mature and immature neurons (Gleeson et al., 1999; Schaar et al., 2004) (Reiner et al., 2006; Shu et al., 2006). The tandem doublecortin domains (DCX1 and DCX2), which drives microtubule assembly function, are connected to the C-terminal kinase domain by a large PEST sequence (linker rich in proline, glutamic acid, serine and threonine) susceptible to proteolytic cleavage (Figure S8A). Calpains are known to cleave DCLK1 at sites mapped to the PEST motif to generate an N-terminal fragment consisting of the two DCX domains and a C-terminal fragment consisting of the intact kinase domain and a C-terminal tail. The cleavage was predicted to regulate the kinase function of DCLK1 (Burgess and Reiner, 2001) (Patel et al., 2016) (Nawabi et al., 2015). Consistent with these previous observations, our TAILS results revealed that treatment of neurons with glutamate resulted in enhanced proteolytic processing of DCLK1 at the following sites: T^311^↓S^312^, S^312^↓S^313^ and N^315^↓G^316^ within the PEST sequence (Figure S8A). Cleavage at these sites is expected to generate N-terminal fragments of ∼35 kDa and C-terminal fragments of ∼49 kDa. In agreement with this prediction, Western blot of the cell lysates derived from the control and glutamate-treated neurons revealed enhanced formation of truncated DCLK1 fragments of ∼50-56 kDa (Figure S8B). As the epitope of the anti-DCLK1 antibody is mapped to the C-terminal tail (Star method), these truncated fragments are predicted to retain the intact protein kinase domain and the C-terminal tail.

### Calpeptin mitigates enhanced proteolysis of neuronal proteins in glutamate-treated neurons

Previous studies by us and others demonstrated that calpeptin, an inhibitor of calpains and cathepsins could protect cultured neurons against excitotoxic cell death (Hossain et al., 2013; Scholzke et al., 2003; Wang et al., 2015). Furthermore, there is ample evidence supporting calpains as the neurotoxic mediators of excitotoxic cell death induced by over-stimulation of iGluRs (Bevers and Neumar, 2008; D’Orsi et al., 2012; Liu et al., 2008). For the 234 and 365 significantly proteolyzed proteins identified in glutamate-treated neurons at these two treatment time points (Figure 1, Tables S5 & S6), we aimed to ascertain which of these proteins were the substrates of calpains and/or cathepsins. We previously demonstrated that calpeptin at 20 μM was able to protect cultured neurons against excitotoxic cell death induced by treatment with 100 μM of glutamate (Hossain et al., 2013). We therefore performed additional TAILS analysis to define the changes in N-terminome of the neurons induced by co-treatment with glutamate and calpeptin (+Glu and Calpeptin). Comparison of the changes in N-terminome of the co-treated neurons with those of the glutamate-treated neurons would unveil which of the significantly proteolyzed proteins in the glutamate-treated neurons were substrates of calpains and/or cathepsins.

By following the rationale depicted in Figure S5, we performed statistical analysis of the TAILS data of neurons co-treated with glutamate and calpeptin to determine the abundance ratio cut-off values for identification of the neo-N-terminal peptides undergoing significant proteolysis in the co-treated neurons (Figures S9 and S10). With the determined threshold values (Figure S10), we found 15 and 52 neo-N-terminal peptides to be derived from neuronal proteins undergoing significant proteolysis at 30 and 240 min of the co-treatment, respectively (Figure 6A). Among them, none and 2 neo-N-terminal peptides, respectively were also found to be derived from proteins undergoing significant proteolysis in neurons at 30 and 240 min of glutamate treatment (Figure 6A, Tables S7, S8 and S9). Hence, for the 234 and 363 cleavage sites in proteins found to be significantly proteolyzed during excitotoxicity, their proteolysis was abolished by calpeptin treatment (Figure 6A). These findings suggest that calpains and/or cathepsins were activated during excitotoxicity and catalyzed their proteolysis. This notion was validated by Western blotting, which showed that the calpeptin mitigated proteolytic processing of neuronal CRMP2 (Figure S11) and DCLK1 (Figure S8B) during excitotoxicity, indicating that calpains and/or cathepsins were their upstream proteases.

**Figure 6.**
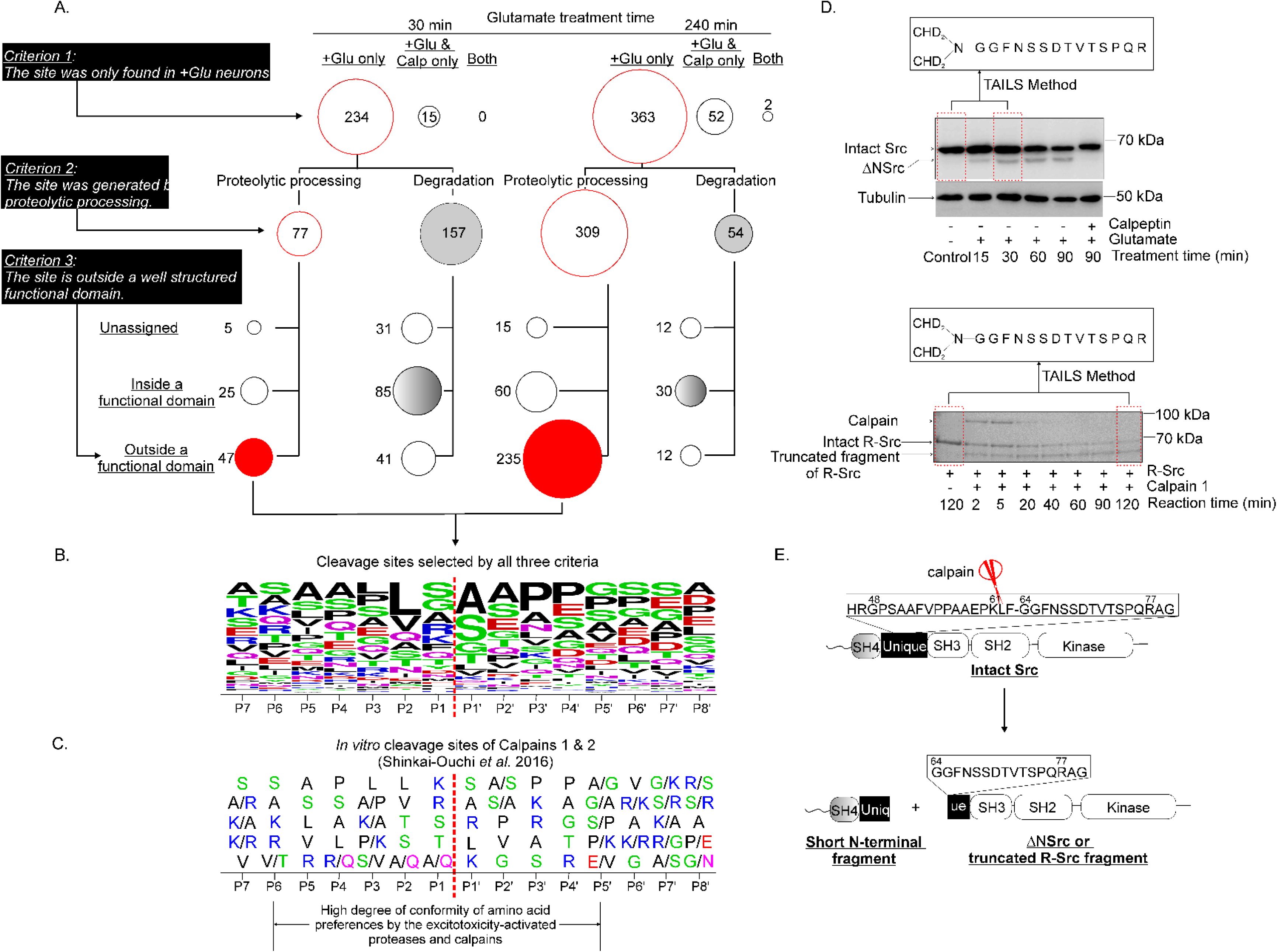
Identification of potential substrates of calpains in excitotoxic neurons. **(A)** Application of the three filtering criteria to define the neo-N-terminal peptides derived from neuronal proteins potentially cleaved by calpains during excitotoxicity. The criteria were applied to the neo-N- terminal peptides derived from neuronal proteins undergoing significantly enhanced proteolysis induced by glutamate treatment or glutamate/calpeptin co-treatment (Figures 3B and S11). The numbers in all groups of significantly changed neo-N-terminal peptides classified by sequentially applying each of the three filtering criteria are shown and depicted in bubble plots. The size of a bubble reflects the number of neo- N-terminal peptides in the group. Bubbles with red perimeter: the groups of peptides selected by Criteria 1 and 2. Red bubbles: the groups of peptides fulfilled all three criteria and are predicted to be derived from neuronal proteins cleaved by calpains during excitotoxicity. +Glu only: peptides identified only in glutamate-treated neurons; +Glu & Calp only: peptides identified only in the glutamate/calpeptin co-treated neurons; both: peptides identified in both the glutamate-treated and the glutamate/calpeptin co-treated neurons. **(B)** The frequencies of amino acids at positions P7 to P8’ proximal to the cleavage sites in the potential calpain substrates (red bubbles). Alignment of the P7-P8’ cleavage site sequences of the potential calpain substrates. The amino acid frequencies at each position are represented in WebLogo format (https://weblogo.berkeley.edu/logo.cgi) (Crooks et al., 2004). The sizes of representation of the listed amino acids in each position reflects the frequencies of its appearance in the identified sequences. **(C)** The top five most frequent amino acids at each position in the P7-P8’ cleavage site sequences in the synthetic peptides proteolyzed by calpain 1 and calpain 2 *in vitro* (Shinkai-Ouchi et al., 2016). The amino acids are presented from top to bottom in order of preference with the top-ranked most frequently encountered amino acid residue listed at the top and the fifth ranked frequently encountered residues listed at the bottom. The positions (P6-P5’) in the cleavage site sequences where both the excitotoxicity-activated proteases and calpains exhibit high conformity of amino acid preferences are shown at the bottom of panel B. Red: D and E; blue: K, R and H; black: ALVIPM; purple: Q and N; Green: S, T and G. **(D)** upper panel: Neuronal Src was cleaved by calpains during excitotoxicity to form a truncated protein fragment ΔNSrc. The cleavage was mitigated by calpeptin treatment. Lower panel: Coomassie blue-stained SDS-PAGE gel of reaction mixtures containing recombinant neuronal Src (R-Src) after incubation with Calpain 1 for 2 min to 120 min *in vitro*. Boxes with dotted red lines: samples analyzed with the TAILS method. Insets: the dimethyl labelled neo-N-terminal peptides identified by TAILS in neurons at 30 min after glutamate treatment and in reaction mixture of R-Src and calpain 1 at 120 min of incubation. The fragment ion chromatograms of these peptides are shown in Figure S16. **(E)** Schematic diagram depicting the functional domains of intact Src and the calpain cleavage site (red scissors) and formation of a short N-terminal fragment and ΔNSrc by limited proteolysis catalyzed by calpains.

### Most proteolytically processed proteins in excitotoxic neurons are potential substrates of Calpains

The three-dimensional structure of the calpain-1 protease core fragment revealed that the active site of calpains is deeper and narrower than that of papain, a cysteine protease with a broad substrate specificity (Moldoveanu et al., 2004). Owing to these structural constraints, the cleavage site motif in a protein substrate are expected to adopt a fully extended conformation before it is accessible to the active site of calpains (Moldoveanu et al., 2004). These findings suggest that the cleavage sites of calpains reside in unstructured regions or motifs with extended conformations outside a functional domain. Based upon these findings, we applied three filtering criteria to identify the potential substrates of calpains from the neo-N-terminal peptides derived from neuronal proteins undergoing significantly enhanced proteolysis induced by glutamate treatment and those induced by glutamate/calpeptin co-treatment (Figures 3B and 6A). These criteria are (i) the neo-N-terminal peptides were found in glutamate-treated neurons but not in the co-treated neurons, (ii) the neo-N-terminal peptides were derived from significantly enhanced proteolytic processing of neuronal proteins with the rationale and thresholds depicted in Figures 3A and S5 and, (iii) the neo-N-terminal peptides were derived from cleavage at sites located outside a functional domain. With these filtering criteria, 47 and 235 neo-N-terminal peptides, respectively were identified to be derived from proteins potentially cleaved by calpains in neurons at 30 and 240 min of glutamate treatment (Figure 6A). Figure S12A shows the locations of the cleavage sites in some of the neuronal proteins identified as calpain substrates by applying these three criteria. CaM kinase 2b, PAK1,3 and 6, creatine kinase β and the regulatory subunit of PKA (prkara) are neuronal proteins with known three-dimensional structures found to undergo significant proteolytic processing in neurons during excitotoxicity. In agreement with the suggestion that calpains cleave intact proteins at sites in unstructured motifs located outside a functional domain (Moldoveanu et al., 2004), the identified cleavage sites in these four proteins are mapped to linker regions, which either adopt a flexible loop structure or a disordered structure (Figure S12A).

Figure 6B shows the weblogo presentation of the frequencies of amino acids proximal to the cleavage sites of neuronal proteins selected by these criteria as potential calpain substrates during excitotoxicity. It is clear that the pattern of amino acid preferences from positions P6-P5’of the cleavage site sequences is very similar to that of the cleavage site sequences in synthetic peptides cleaved by calpain 1 and calpain 2 *in vitro* (Shinkai-Ouchi et al., 2016) (Figure 6C). For example, bulky hydrophobic residues at the P2 position and proline residue at P3’ position were the most preferred amino acid residues in the cleavage site sequences of *in vitro* peptide substrates of calpain 1 and calpain 2 and the cleavage sites of the potential calpain substrates selected by these three criteria. Such a high degree of similarity lends further support to the reliability of the three criteria we used to identify the potential substrates of calpains in neurons during excitotoxicity. As such, we postulate that calpains catalyze limited proteolysis of these neuronal proteins to generate stable truncated fragments during excitotoxicity.

When a protein undergoes degradation for clearance, the multiple degradation steps generate numerous unstructured or partially folded intermediate peptide fragments originating from both functional domains and the disordered structural motifs. As such, the cleavage sites of the degraded proteins are not expected to preferentially reside in unstructured motifs outside a functional domain. Consistent with this prediction, our bioinformatic analysis revealed that the identified cleavage sites in most of the significantly degraded neuronal proteins during excitotoxicity are mapped within functional domains with well-defined three-dimensional structures (Figures 6A). Vdac1, Creatine kinase and Pyruvate kinase are neuronal proteins with known three-dimensional structures (Figure S12B), which was found by our TAILS study to undergo significantly enhanced degradation during excitotoxicity. The identified cleavage sites in these proteins are located within functional domains with well-defined structures. The three-dimensional structures clearly show that in the intact proteins, the identified cleavage sites are not accessible to proteases (Figure S12B). Hence, the neo-N-terminal peptides identified by our TAILS analysis were likely derived from the intermediate peptide fragments generated during proteolytic degradation of the neuronal proteins for clearance during excitotoxicity (Figure S12B). As cathepsins catalyze proteolysis of cellular proteins destined for clearance, proteins undergoing significantly enhanced degradation during excitotoxicity identified in our study were likely direct substrates of cathepsins in neurons.

Neuronal Src was identified by the three criteria of our bioinformatic analysis to be cleaved by calpains to form a stable truncated protein fragment during excitotoxicity (Figures 6A and Table S6). The abundance of the dimethyl labelled N-terminal peptide (dimethyl-G^64^-GFNSSDTVTSPQR^77^) derived from Src was 5.6-fold higher in the glutamate-treated neurons versus that in the untreated neurons (M/L ratio = 5.6) (Table S6), indicating significantly enhanced proteolytic processing of neuronal Src by an excitotoxicity-activated protease catalyzing cleavage of Src at the F^63^-G^64^ bond (the cleavage site is referred to as F^63^↓G^64^) (Figure S13). The cleavage would generate a truncated Src fragment of 54 kDa. These proteomic findings are validated by the Western blot results shown in upper panel of Figure 6D, and are in agreement of our previous findings that glutamate treatment led to the formation of a truncated Src fragment of ∼54 kDa in neurons (Hossain et al., 2013). The F^63^-G^64^ bond resides at the intrinsically disordered motif of the Unique domain of Src (Arbesu et al., 2017) and calpeptin abolished its cleavage when neurons were co-treated with glutamate and calpeptin (Figure 6D). As such, it is a potential calpain cleavage site. To validate this prediction, we incubated recombinant Src (R-Src) with calpain 1 *in vitro* and found that R-Src was cleaved by calpain 1 to form a 54 kDa truncated fragment as early as 2 min after incubation. To determine the cleavage site in R-Src targeted by calpain 1, the reaction mixture consisting of intact R-Src only and that consisting of R-Src and calpain-1 after 120 min of incubation were subjected to isotopic dimethyl labelling prior to tryptic digestion (Figure 6D, lower panel). A dimethyl labelled peptide identical to the dimethyl labelled neo-N-terminal peptide (dimethyl-G^64^-GFNSSDTVTSPQR^77^) detected in the lysate of glutamate-treated neurons, was present only in the R-Src/calpain-1 reaction mixture but not in the reaction mixture with R-Src only (Figure S13). These results validated our prediction and suggest that calpain 1 or another calpain directly cleaved neuronal Src at the F^63^↓G^64^ bond during excitotoxicity (Figure 6E). The cleavage is expected to generate two truncated fragments: (i) a truncated Src fragment (ΔNSrc) containing the SH2, SH3 and kinase domains and the C-terminal tail and (ii) a short N-terminal fragment consisting of the SH4 domain and a large portion of the Unique region, which were previously found to adopt highly disordered conformations (Figure 6E) (Arbesu et al., 2017; Perez et al., 2009). Since the SH4 domain and Unique region are critical to anchoring of Src to plasma membrane and intracellular vesicular membrane (Amata et al., 2014; Arbesu et al., 2017), most ΔNSrc resides in the cytosol of neurons (Hossain et al., 2013).

### The role of calpain-mediated cleavage site of Src in excitotoxic neuronal death

Src was predicted by phoshoPATH to be a hub of protein interactions in excitotoxic neurons (Figure 4A), suggesting that it is a major player in regulation of neuronal survival during excitotoxicity. Previous studies by us demonstrated that dysregulation of Src and its related kinase Fyn is a key contributor of excitotoxic cell death of cultured cortical neurons and neuronal loss in a rat model of traumatic brain injury (Hossain et al., 2013; Liu et al., 2017). As such, we selected Src as a target for the development of potential therapeutic agents capable of protecting against excitotoxic neuronal death. Our discovery that neuronal Src is cleaved by calpains during excitotoxicity prompted us to investigate if blockade of calpain cleavage of Src in neurons is a neuroprotective strategy.

Based upon the findings of calpain-mediated cleavage of neuronal Src during excitotoxicity (Hossain et al., 2013), we designed a cell-permeable peptide TAT-Src consisting of the cell-permeable TAT-sequence and the segment encompassing residues 49-79 in the unique domain of Src and a cell-permeable control peptide Tat-Scrambled (Figure S14A). We demonstrated that Tat-Src but not Tat-Scramble could block cleavage of Src in neurons in excitotoxicity (Figure S14B) (Hossain et al., 2013). Importantly, Tat-Src but not Tat-Scrambled could protect the cultured cortical neurons against excitotoxic cell death (Hossain et al., 2013), indicating that Tat-Src peptide is a neuroprotective agent of cultured neurons.

We next examined the potential therapeutic value of our findings by examining if TAT-Src can enter neurons and protect against excitotoxic neuronal death *in vivo* in an animal model of neurotoxicity. We first examined if TAT-Src peptide could enter neurons. To this end, we synthesized FITC-TAT-Src, a derivative of TAT-Src peptide with a fluorescent tag covalently attached at the N-terminus. It was stereotaxically infused into the cortical and striatal regions of rat brains (Figures 7A and 7B). At one-hour post infusion, FITC-TAT-Src was detected in neurons but not astrocytes and microglia in the infused regions of the rat brain (Figure 7A), suggesting that it could enter neurons and block calpain-mediated cleavage of neuronal Src.

**Figure 7.**
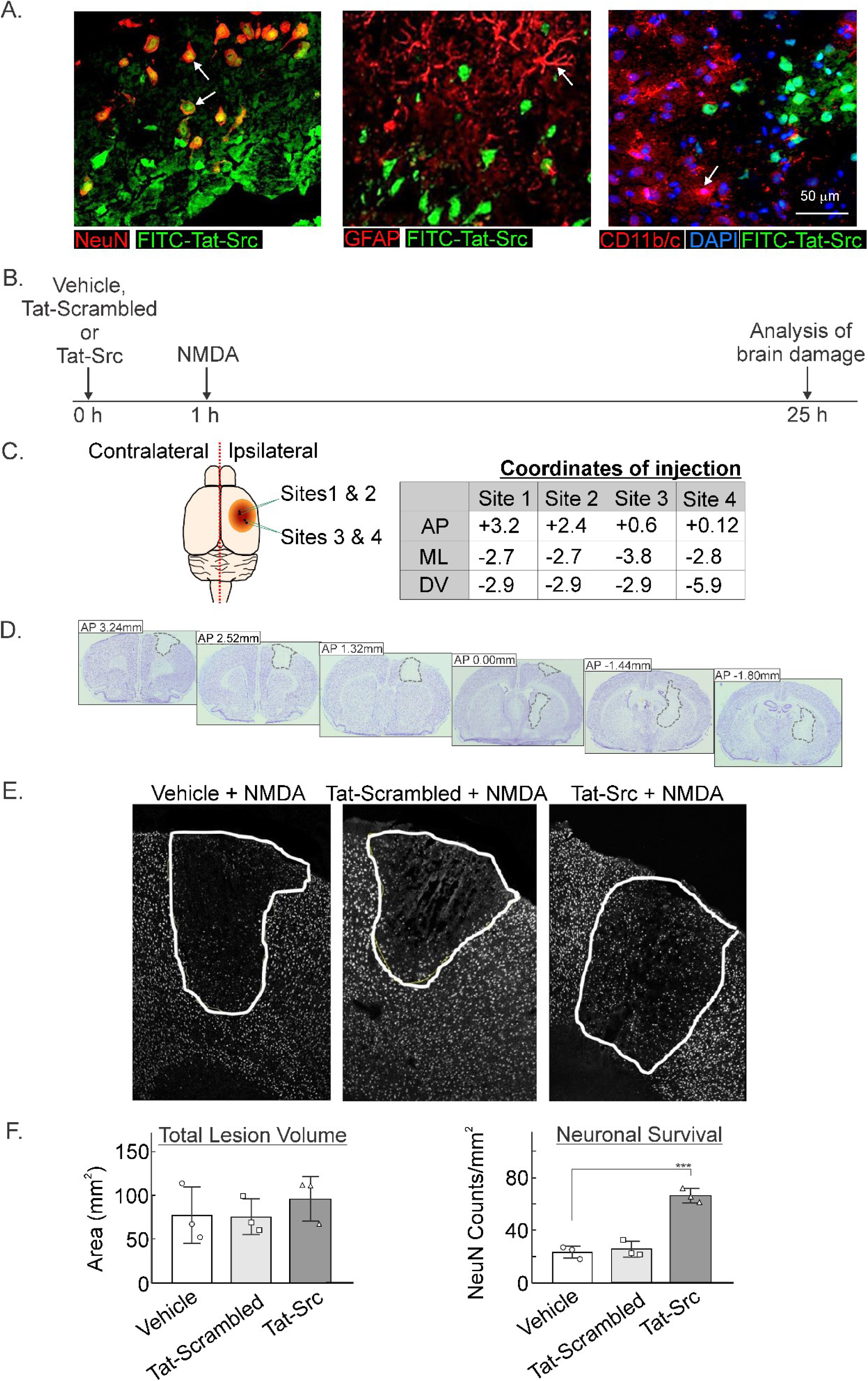
TAT-Src protects against neuronal loss *in vivo* in NMDA-mediated neurotoxicity. **(A)** Representative photomicrographs of FITC labelled TAT-Src infusion co-labelled against markers of neuronal cells (NeuN+ red); astrocytes (GFAP+; red) and microglia (CD11b/c OX-42+, red) and observed as orange. (B) Time line depicting treatment with Vehicle (Milli-Q H_2_O, 3 μl), Tat-Src (5 mM in Milli-Q H_2_O, 3 μl) or Tat-Scrambled (5 mM in Milli-Q H_2_O, 3 μl) at 1 hour prior to NMDA-induced excitotoxicity. (C) Stereotaxic coordinates of the four injection sites (sites 1 - 4) used to cerebrally inject NMDA to induce excitotoxicity (70 mM in PBS, 1 μl per site). (D) Representative thionin-stained coronal images of rat brains infused with NMDA to demonstrate damage to the motor cortex and dorsal striatum. (E) NeuN+ cells (transposed white using Image J software) in all three treatment groups. (F) Left panel: the total lesion volumes in the treatment groups. Right panel: the total number NeuN+ cells in each treatment group was point-counted using image J software with the number of surviving neurons within the lesion significantly increased in rats treated with TAT-Src (p<0.0001, n=3/group, one-way ANOVA) followed by the Bonferroni post-hoc test. Data were analyzed using GraphPad Prism, version 8 and presented as mean ± SD. Statistical significance was defined as p <0.05 for infarct volume and *p* < 0.0001 for NeuN cell count.

To examine the ability of TAT-Src to protect against excitotoxic neuronal death *in vivo*, we chose to use an *in vivo* rat model of neurotoxicity. In this model, TAT-Src peptide, vehicle (water) or TAT-scrambled peptide was stereotaxically injected at four sites in the cortical and striatal regions of rat brains (Figures 7B and 7C). One hour after the injection, neurotoxic dose of NMDA was infused to the same sites to induce excitotoxic brain damage. At 24 h after the infusion of NMDA, the rats were sacrificed, and brain sections were prepared to measure the infarct volume and the number of surviving neurons (Figure 7D). The absence, or reduction in NeuN immunoreactivity, revealed NMDA-induced lesions within the motor and parietal cortex (Figures 7D and 7E). The plot in left panel of Figure 7F show that total lesion volume was consistent across all treatment groups with no significant difference in the volume of damage detected between groups. Stereological point counting of NeuN positive cells within the lesion revealed treatment-specific effects where the number of neurons in rats treated with TAT-Src was significantly higher than in rats receiving vehicle or scrambled Tat-Src control (Figure 7F, right panel). Thus, injection of TAT-Src prior to NMDA infusion could protect against excitotoxic neuronal loss caused by the injected NMDA. Since FITC-TAT-Src, a derivative of TAT-Src could enter neurons but not astrocytes and microglia (Figure 7A), the ability of TAT-Src to protect against neuronal loss in NMDA-induced brain damage is attributed to its effects in neurons. Presumably, the effect is the result of the ability of TAT-Src to prevent cleavage of neuronal Src to form the neurotoxic truncated Src fragment (ΔNSrc) (Hossain et al., 2013). In conclusion, our results illustrate the therapeutic potential of blockade of cellular events in excitotoxic neurons uncovered by our proteomic analysis.

## DISCUSSION

Our multi-proteomic analysis of excitotoxic neurons demonstrate that over-stimulation of iGluRs perturbs intracellular signalling pathways governing synaptogenesis, cell adhesion, cell survival and mRNA processing. One of our major findings was the enhanced proteolytic cleavage and/or aberrant phosphorylation of hundreds of proteins in excitotoxic neurons. From the identities and sequences of phosphosites in some neuronal proteins exhibiting significant increase in phosphorylation during excitotoxicity, over twenty protein kinases are predicted to be activated and directly phosphorylate these phosphosites during excitotoxicity. Based on our findings, we now have a shortlist of specific pharmacological tools such as inhibitors of the predicted activated kinases shown in Figure 4 to further chart the signalling pathways directing excitotoxic neuronal death. Experimental approaches to chart the cytotoxic signalling pathways in excitotoxic neurons are discussed in the sections below.

Besides contributing to the construction of an atlas documenting thousands of cellular events occurring in the neurons during excitotoxicity, the present study also advances the field of calpain research. As modulator proteases playing key roles in many pathophysiological processes, calpains perform their functions by catalyzing limited proteolysis of specific cellular proteins to generate stable truncated protein fragments with modified regulatory properties and cellular functions. Hence, determination of the cleavage sites in cellular proteins targeted by calpains in cells and *in vivo* is an avenue to define how the calpain-catalyzed cleavage modulates the cellular functions of the proteins.

### The cleavage sites in neuronal proteins undergoing significantly enhanced proteolytic processing during excitotoxicity are potential calpain sites

Mechanisms of substrate recognition by calpains still remain poorly understood because of the relative scarcity of confirmed calpain cleavage site data. However, duVerle and Mamitsuka unveiled around one hundred experimentally confirmed calpain cleavage sequences in mammalian proteins (duVerle and Mamitsuka, 2019). Among them, the exact sites of calpain cleavage sites were determined by *in vitro* study. Based upon the limited data of confirmed calpain cleavage sites in protein substrates and results of peptide library studies to define the substrate specificity determinants near the cleavage sites in synthetic peptides (Cuerrier et al., 2005; Shinkai-Ouchi et al., 2016), a number of algorithms for prediction of calpain substrates and the cleavage sites were designed. The most notable algorithms include calCleaveMKL (duVerle and Mamitsuka, 2019), iProt-Sub (Song et al., 2019), DeepCalpain (Liu et al., 2019) and GPS-CCD (Liu et al., 2011). Here, we demonstrated for the first time a high degree of conformity of amino acid preferences at the P6-P5’ positions in both the potential calpain substrates in neurons and the *in vitro* peptide substrates of calpains 1 and 2 (Figures 6B and 6C). Calpains 1 and 2 are the major calpain isoforms expressed in neurons. Our findings suggest that determinants governing substrate specificity of calpains 1 and 2 are present in the P6-P5’ positions proximal to the cleavage sites in the properly folded calpain protein substrates. Among the 200-300 cleavage sites in neuronal proteins potentially cleaved by calpains during excitotoxicity identified in our TAILS study (Figure 6), over 90% of them were identified for the first time as potential cleavage sites in properly folded proteins in live cells. We anticipate that incorporating information of the cleavage site sequences and/or three-dimensional structures of the corresponding calpain substrates will improve the predictive accuracy of these algorithms.

Besides conformity of the primary structure proximal to the cleavage site with the optimal calpain cleavage sequence and the three-dimensional structural features of a potential calpain substrate, its co-localization with a specific isoform of calpains also governs whether it is cleaved by calpains *in vivo*. For example, the C-terminal tail of calpains 1 and 2 contain different PDZ-binding motifs which target them to different subcellular compartments where they proteolyze specific subsets of protein substrates (Baudry and Bi, 2016; Wang et al., 2013). The N-terminal part of calpain 1 but not calpain 2 contains a amphipathic α-helical domain, which specifically targets it to mitochondria (Badugu et al., 2008). Future investigations to decipher where and when the potential calpain substrates identified in our TAILS analysis form protein complexes with calpains 1 and 2 in neurons during excitotoxicity will further bridge the knowledge gap concerning how the two isoforms of calpains recognize their substrates in neurons.

### Experimental approaches for future investigation to decipher the signalling pathways directing excitotoxic neuronal death

One approach to decipher the signalling pathways directing excitotoxic neuronal death involves the use specific inhibitors of neurotoxic cellular events that occur at the early phase of cell death. The major over-stimulated iGluRs governing excitotoxic neuronal death are NMDA receptors. In neurons, NMDA receptors are located both at the synapses and in extrasynaptic regions. It is well established that the extrasynaptic NMDA receptors enriched in the GluN2B subunit, upon stimulation in pathological conditions such as excitotoxicity, emanate cytotoxic signals to direct neuronal death (Hardingham et al., 2002; Kaufman et al., 2012). Further studies established that the GluN2B C-terminal tail of stimulated extrasynaptic NMDA receptors form complexes with the scaffold protein PSD95, which couples the stimulated receptors with NADPH oxidase 2 (NOX2) and neuronal nitric synthase (nNOS) (Brennan et al., 2009; Girouard et al., 2009; Sattler et al., 1999). The complex formation facilitates over-activation of NOX2 and nNOS to form excess reactive oxygen species and nitric oxide, respectively. Nerinetide was modelled after the PSD95-binding motif in GluN2B C-terminal tail (Aarts et al., 2002; Hill et al., 2020). By blocking the interactions between the aberrantly stimulated extrasynaptic Glu2B-enriched NMDA receptors with PSD95, Nerinetide inhibits over-activation of nNOS (Aarts et al., 2002) and NOX2 (Chen et al., 2015) and in turn protects against excitotoxic neuronal death *in vitro* and *in vivo* in rodent and primate models of neurotoxicity (Cook et al., 2012; Sun et al., 2008). More importantly, Nerinetide has been shown to reduce brain damage caused by micro-strokes as well as strokes caused by large vessel occlusion (Hill et al., 2020; Hill et al., 2012). Hence, Nerinetide is a useful tool to be used in conjunction with proteomic approaches to decipher the signalling events operating downstream of nNOS and NOX2 over-activated by the extrasynaptic NMDA receptors in excitotoxic neurons.

Besides activation of nNOS and NOX2, the over-stimulated NMDA receptors also aberrantly activate calpains. Studies of the knock-out mice deficient of the endogenous calpain inhibitor calpapstatin, transgenic mice overexpressing calpapstatin, and mice and neurons treated with chemical inhibitors of calpains indicate that calpains are the major mediators of excitotoxic neuronal death (Schoch et al., 2012; Takano et al., 2005; Wang et al., 1996). Studies by Baudry and co-workers provided evidence that calpain 1 is selectively activated by synaptic NMDA receptors while calpain 2 is activated by the GluN2B-enriched extrasynaptic NMDA receptor (Wang et al., 2013). However, previous studies from different groups of researchers provided conflicting evidence of the roles of calpain 1 and calpain 2 in mediating the neuroprotective signals of the synaptic NMDA receptors and the neurotoxic signals of extrasynaptic NMDA receptors(Wang et al., 2013; Xu et al., 2009; Yamada et al., 2012). Future investigation to define the proteomic changes induced by glutamate treatment in calpain 1-deficient neurons and calpain 2-deficient neurons will decipher the roles of these two calpain isoforms in mediating the neurotoxic signals of the over-stimulated NMDA receptors during excitotoxicity.

Results from the present study and our previous studies show that the neuroprotective Tat-Src peptide, specifically blocks calpain cleavage of neuronal Src to form truncated Src fragment ΔNSrc (Hossain et al., 2015; Hossain et al., 2013). We previously provided convincing evidence that ΔNSrc is a mediator of the neurotoxic signals of the over-stimulated NMDA receptors. It causes neuronal death at least in part by inhibiting the pro-survival Akt kinase (Hossain et al., 2013). Additionally, the kinase-dead mutation abolishes the neurotoxic action of ΔNSrc (Hossain et al., 2013). These findings suggest that ΔNSrc exerts its neurotoxic action by phosphorylating specific neuronal proteins, and one of more of the neuronal proteins, upon phosphorylation by ΔNSrc, contributes to neuronal death by suppressing activity of Akt. Besides being a critical regulator of neuronal survival, Akt plays critical roles in regulation of synaptogenesis of adult neurons (Cuesto et al., 2011), suggesting that inhibition of Akt by ΔNSrc also perturbs synaptogenesis of neurons during excitotoxicity. Thus, these findings lend further credence to the prediction of significant perturbation of the canonical signalling pathways governing neuronal survival, synaptogenesis, cell adhesion and cell-cell communications during excitotoxicity (Figure S7).

In light of these findings, future investigation to define the changes in phosphoproteome and N-terminome of neurons in response to glutamate treatment and those of neurons co-treated with glutamate and Tat-Src peptide will define the neurotoxic signalling events downstream of calpain cleavage of Src. Since ΔNSrc is a protein tyrosine kinase, identifying the proteins directly phosphorylated by ΔNSrc in neurons is an appropriate approach to define its neurotoxic mechanism. The proteomic method developed by Bian, *et al*. to define the phosphotyrosine-proteome of cultured cells using the superbinder mutants of SH2 domains (Bian et al., 2016) is therefore a method of choice to identify the substrates of ΔNSrc in neurons during excitotoxicity.

### Neuronal proteins proteolytically processed during excitotoxicity are potential targets for the development of neuroprotective therapeutics

Currently, there is no FDA-approved neuroprotective drug for treating patients suffering from acute neurological disorders or neurodegenerative disease. With so many likely candidates failing to impact clinical disease management, new selection and screening criteria call for the development of novel therapies that reduce neuronal loss whilst not inferring with normal physiological processes. Potential new therapies should interact only with their targets during states of pathological activation but not interfere with the targets under physiological conditions (Lipton, 2007). Of relevance in the present study, 363 neuronal proteins were found to undergo significantly enhanced proteolysis as a result of aberrant activation of calpains and cathepsins during excitotoxicity (Figure 6A). Among them, some underwent limited proteolysis catalyzed by the aberrantly activated calpains and/or cathepsins to form stable truncated fragments, which may mediate the neurotoxic signals originating from the over-stimulated iGluRs. Since calpains and cathepsins are aberrantly activated in pathological conditions, these neuronal proteins are potential targets for the development of neuroprotective drugs. Small-molecule inhibitors selectively blocking their proteolysis by calpains and/or cathepsins during excitotoxicity are potential neuroprotective drugs. In this and our past studies, we demonstrated that blockade of calpain cleavage of Src to form the neurotoxic truncated fragment ΔNSrc during excitotoxicity could protect against excitotoxic cell death of cultured neurons (Hossain et al., 2013; Iqbal Hossain et al., 2015) and neuronal loss *in vivo* (Figure 7). These findings suggest that small-molecule compounds that selectively inhibit cleavage of Src by calpains are potential neuroprotective drug candidates to reduce excitotoxic neuronal loss in neurological disorders. In addition to Src, many other neuronal proteins were found by our TAILS study to be cleaved by the aberrantly activated calpains and/or cathepsins to form stable truncated fragments in excitotoxicity. For example, we discovered for the first time, cleavage of neuronal Fyn at the unique domain during excitotoxicity (Table S9). The cleavage is expected to generate a truncated Fyn fragment (ΔNFyn) with a high degree of structural similarities to the neurotoxic ΔNSrc. We therefore postulate that ΔNFyn is neurotoxic, and blockade of calpain and/or cathepsin-mediated cleavage of Fyn to form ΔNFyn is a potential neuroprotective strategy. Likewise, for the >200 intact neuronal proteins such as CRMP2, DCLK1 and CAMK2b cleaved by the aberrantly activated calpains and/or cathepsins to form stable truncated fragments (Figures 5, S8 and S12A), if future investigation found that their cleavage is critical to neuronal death, small molecules blocking their cleavage by calpains and/or cathepsins are potential neuroprotective drug candidates.

## STAR METHODS

### KEY REOURCES TABLES

KEY RESOURCES TABLES
| Antibody | Catalogue number | Manufacturer | RRID |
| --- | --- | --- | --- |
| Rabbit monoclonal anti-CRMP2 antibody | 35672 | Cell Signalling Technology | RRID:AB_2799082 |
| Anti-CRMP2 (phospho T509) antibody (ab192799) | ab192799 | Abcam |  |
| Anti-DCLK1 (Anti-DCMKL1) antibody against amino acids 679-729 near the C-terminus of DCLK1. | Ab106635 | Abcam |  |
| Anti-Src Mab327 (equivalent to anti-Src antibody (Clone 327) supplied by Abcam Cat# ab16885) |  |  | RRID:AB_443522 |
| Anti-CD11b (Clone OX-42) mouse monoclonal (formerly serotec) | MCA275G | BioRad |  |
| NeuN (rabbit polyclonal) | ABN78 | MERK Millipore | RRID:AB_10917954 |
| Anti-GFAP (mouse monoclonal) (Clone GA5) | MAB360 | MERK Millipore (Sigma Aldrich) | RRID:AB_185664 |

| Chemicals | Catalogue Number | Manufacturer |
| --- | --- | --- |
| Pierce™ High pH Reversed-Phase Peptide Fractionation Kit | 84868 | ThermoFisher |
| High-Select™ Fe-NTA Phosphopeptide Enrichment Kit | A32992 | ThermoFisher |
| Titansphere Phos-TiO<sub>2</sub> | 5010-21315 | GL Sciences |
| TEAB buffer | T7408 | Sigma |
| TCEP | 20490 | ThermoFisher |
| Hydroxy Polyglycerol Aldehyde (HPG-ALD) Polymer |  | http://clip.ubc.ca/<br>Kleifeld et al., 2010 |
| Formaldehyde | DLM-805-PK (CD<sub>2</sub>O)<br>ULM-9498-PK (CH<sub>2</sub>O) | Cambridge Isotope |
| Fmoc-PAL-PEG-PS™ (Rink resin; loading capacity 0.21 mmol/g) | GEN913383 | ThermoFisher |
| Fmoc-protected amino acids |  | GL Biochem (Shanghai) |

| Type of animals | Strain | Usage | Sources & RRID |
| --- | --- | --- | --- |
| Mice | C57BL/6 mice (pregnant and at gestational day 14-15) | Cultured primary cortical neurons | Animal Resources Centre Canning Vale WA 6970 Australia |
| Mice | C57BL/6 mice | Mouse brains for spectral libraries | Animal Resources Centre Canning Vale WA 6970 Australia |
| Rats | Hooded Wistar (male) | *In vivo* model of neurotoxicity to test the neuroprotective action of TAT-SRC peptide | Animal Resources Centre Canning Vale WA 6970 Australia |

| Software and algorithm | Source and RRID |
| --- | --- |
| Proteome Discoverer | Thermo Scientific; RRID:SCR_014477 |
| MaxQuant | RRID:SCR_014485 |
| Spectronaut (Spectronaut 11) | Biognosys |
| PhosphoSitePlus: Protein Modification Site | Cell Signalling Technology RRID:SCR_001837 |
| Image J | RRID:SCR_003070 |

| Deposited Data | Source | Identifier |
| --- | --- | --- |
| Proteomics raw and analysed data | This paper |  |

### EXPERIMENTAL MODELS

#### Preparation of cultured mouse primary cortical neurons

The procedures were approved by the University of Melbourne Animal Ethics Committee (Licence number: 161394.1) and were performed in accordance with the Prevention of Cruelty to Animals Act 1986 under the guidelines of the National Health and Medical Research Council Code of Practice for the Care and Use of Animals for Experimental Purposes in Australia. Briefly, embryos were collected from pregnant C57BL/6 mice (gestational day 14-15) after they were euthanized by CO_2_ asphyxiation. The cortical region was aseptically micro-dissected out of the brains of the embryos, free of meninges and digested with 0.025 % (w/v) trypsin in Krebs Buffer (0.126 M NaCl, 2.5 mM KCl, 25 mM NaHCO3, 1.2 mM NaH2PO4, 1.2 mM MgCl2, 2.5 mM CaCl2, pH 7.2) and incubated at 37°C with shaking to dissociate the connective tissues. After 20 min of incubation, 0.008% (w/v) DNase I (Roche Applied Science) and 0.026 % (w/v) soybean trypsin inhibitor (Sigma) in 10 ml Krebs solution (DNase I/soybean trypsin inhibitor solution) were added to the suspension to stop the trypsin action and initiate digestion of DNA. Gentle mixing by inversion of the suspension was performed. Cells in the suspension were collected by centrifugation at 1000 × g for 3 min at room temperature. They were resuspended in 1 ml of DNase I/soybean trypsin inhibitor solution. After aspiration of the supernatant, the cells were resuspended in plating medium (minimum essential medium (MEM) supplemented with 2 mM L-glutamine, 0.22% v/v NaHCO3, 1% penicillin-streptomycin, 5% v/v horse serum and 10% v/v foetal calf serum). Approximately 800,000 cells per well were seeded to a 12-well plate pre-treated with 0.1 mg/ml Poly-D-lysine. After incubation at 37°C in 5% CO_2_ for 2 h, the medium was replaced with neurobasal medium supplemented with 0.2 mM L-glutamine, 0.01 mg/ml penicillin-streptomycin and B27 supplement (NB/B27). Cultures were maintained for seven days (days *in vitro* 7 (DIV7)). The DIV7 cultures, highly enriched with neurons, were used for the experiments.

To induce excitotoxicity, the DIV 7 neuronal cultures were treated with 100 μM glutamate in NB/B27 for 30 and 240 min. For co-treatment with glutamate and calpeptin, the DIV 7 neuronal cultures were treated with 100 μM glutamate and 20 μM calpeptin in NB/B27 for 30 and 240 min. For control, untreated cells from the same batch of DIV 7 neuronal cultures were used for proteomic and biochemical analyses.

#### Mouse models of ischemic stroke and traumatic brain injury, related to Figures 1 and S2

##### Animals

All experiments were performed in strict accordance with the guidelines of the National Health & Medical Research Council of Australia Code of Practice for the Care and use of Animals for Experimental Purposes in Australia. For the dual carotid artery ligation stroke (ischemic stroke) mouse model and the controlled cortical impact mouse model of traumatic brain injury (TBI), C57Bl/6 mice (20-30g), male, were used. Animals were held in the Australian Medical Research and Education Precinct (AMREP) animal house facility and kept on a 12 h light/dark cycle with access to food and water *ad libitum*. These studies were approved by the AMREP Animal Ethics Committee (E/1683/2016/M). For the animals used in the rat model of neurotoxicity, male Hooded Wistar rats weighing 300-340g sourced from Laboratory Animal Services, University of Adelaide, Australia were used. The protocol was approved by the St Vincent’s Hospital animal ethics committee (AEC016/12). All surgery was performed under general anaesthesia, and paracetamol (2 mg/kg in drinking water) was provided for 24 h prior to and after surgery in order to minimize suffering and included monitoring each rat throughout the length of the study to ensure their wellbeing.

##### Dual Carotid Artery Ligation (DCAL) Stroke Model

Mice were placed into a plastic box and initially anaesthetised with 5% Isoflurane (in 1.0 ml/min O_2_), and maintained on 1.5% Isoflurane for the duration of the experiment. A temperature probe connected to a thermoblanket (Harvard Apparatus Ltd., Kent, UK) was inserted into the rectum to monitor body temperature, and body temperature was maintained at 37°C throughout the experiment using a heat lamp.

All surgical procedures were performed using aseptic technique. Surgical instruments were sterilised using a bead steriliser (Steri350, Sigma Aldrich) prior to use. The mouse was affixed in a supine position on the thermoblanket using tape, and the neck shaved and swabbed with alcohol. Through a ventral midline incision, the carotid arteries were exposed via blunt dissection and carefully dissected clear of the vagus nerve and surrounding tissue. A stabilisation period of 10 min is allowed between the isolation of each artery. The right jugular vein was also exposed via a cut to the skin and blunt dissection for the purpose of drug administration. Following the surgical procedures, the animals were allowed to stabilise for 10 min before the experiment proceeded. The left carotid artery is permanently ligated using 6-0 silk suture (6-0 black braided silk suture, SDR Scientific, Sydney, Australia). Following a stabilisation period of 10 min, the right carotid artery is then transiently occluded for 30 min using a small haemostatic clamp. After the transient occlusion, the neck incision was closed using tissue glue (Leukosan Adhesive, BSN medical Inc.), and the mouse allowed to recover under 1% O_2_. Animals were housed separately following surgery with access to food and water *ad libitum*. At 24 h post-surgery, the mice were euthanized, and their brains removed and stored at −80 °C.

##### Controlled cortical impact model of traumatic brain injury

For the induction of traumatic brain injury (TBI) the controlled cortical impact (CCI) model was performed. This procedure is well established and induces a reproducible brain trauma with a mortality of less than 5% (Sashindranath et al., 2011). After anaesthesia with 0.5 g/kg avertin (1.875% solution dissolved in 0.9% sodium chloride pre-warmed at 37°C, 2,2,2-tribromoethanol; Sigma Aldrich #T48402; 1mg/ml in tert-amyl alcohol), injected intraperitoneally (i.p.), mice were placed in a stereotaxic frame (Kopf, Tujunga, CA). A sagittal incision in the midline of the head was performed and the skull cleaned with a 6% hydrogen peroxide solution using a sterile cotton swab. This was followed by a 5 mm diameter craniotomy performed with a drill over the left parietal cortex. The impactor was positioned in a 20° angle with the tip (cylindrical rod) touching the brain surface, and a single blunt force trauma was inflicted to the exposed brain area with an impact depth of 2mm, a velocity of 5 m/s and dwell time of 400 ms inducing a moderate to severe brain trauma. The exposed brain was then sealed using bone wax (Ethicon, Johnson and Johnson #W810T) and the skin incision was sutured with a non-absorbable braided treated silk (Dynek, Dysilk) and treated with a local anaesthetic (xylocaine) and an antiseptic (betadine). For the sham procedure, only the scalp incision, without craniotomy and CCI was performed since even the craniotomy without CCI results in a brain lesion (Sashindranath et al., 2015). Regardless of the experimental design, however, mice were placed on a 37 °C heat pad within 30 min after induction of anaesthesia for post-surgery recovery, and they usually recovered within 40-60 min. Animals were housed separately following surgery with access to food and water *ad libitum*. At 24 h post-surgery, the mice were euthanized, and their brains removed and stored at −80 °C.

#### *In vivo* model of NMDA neurotoxicity, related to Figure 8

##### Induction of NMDA excitotoxicity in vivo

Male Hooded Wistar rats (n = 12) weighing 300-340g (Laboratory Animal Services, University of Adelaide, Australia) were used in this study. Rats were anesthetized with intraperitoneal administration of ketamine and xylazine (75mg/kg and 10 mg/kg, respectively), and maintained by inhalation of isoflurane (95% oxygen and 5% isoflurane). Rats were positioned in a stereotaxic frame (Kopf), and 4 burr holes were drilled into the right hemisphere corresponding to the predetermined sites for NMDA infusion (Site 1: AP +3.2, ML −2.7, DV −2.9; Site 2: AP +2.4, ML −2.7, DV −2.9; Site 3: AP +0.6, ML −3.8, DV −2.9; Site 4: AP +0.12, ML −2.8, DV −5.9). Rats were randomly assigned into 3 cohorts and received either vehicle (sterile Milli-Q H_2_O, 3 μl per site), TAT-SRC peptide (5 mM in sterile MilliQ H_2_O, 3 μl per site), or TAT-Scrambled peptide (5 mM in sterile MilliQ H_2_O, 3 μl per site) via direct infusion at 0.2 μl/min into each site 1 h prior to infusion of NMDA (70 mM, 1μl PBS per site). Following infusion each burr hole was filled with bone wax and wounds sutured. In separate studies, rats were infused with FITC-TAT-SRC peptide without NMDA to assess neuronal uptake and cell specificity at one hour after infusion.

Rats that received treatment with NMDA +/- Tat-Src, Tat-Scrambled or Vehicle were allowed to recover for 24 h prior to lethal overdose (lethobarb) and decapitation. Forebrains were collected and rapidly frozen over liquid nitrogen and stored at −80°C prior to processing. Coronal sections (16 µm thick) were prepared using a Leica cryostat (Leica Microsystems, Wetzlar, Germany) across the 4 coronal planes corresponding to the NMDA infusion sites.

##### Immunohistochemistry

Immunofluorescence staining was performed in forebrain tissue sections to identify Tat-Src cell specificity and treatment effects following NMDA excitotoxicity. Sections were first fixed in 4% PFA for 15 minutes at RT prior to wash (3 × 5 min washes with 0.1 M PBS) and a blocking step in 5 % NGS and 0.3 % Triton X-100 and 0.1 M PBS for 30 min. Sections were again washed (3 × 5 min washes with 0.1 M PBS) and adjacent sections incubated with primary antibodies to detect neurons, astrocytes and microglia using the following primary antibodies: mouse anti-NeuN (1:500, Chemicon); mouse anti-GFAP (1:400, Millipore); and mouse anti-OX-42 (1:100, Serotec) in 2% NGS, 0.3% Triton X-100 and 0.1M PBS overnight at 4°C. The following day sections were again washed (3 × 5 min washes with 0.1 M PBS) and incubated with secondary fluorophore-conjugated antibody Alexa 555 goat anti-mouse (1:500, Thermo Fisher Scientific) for visualisation of each primary antibody. For all experiments, DNA counterstain DAPI (Molecular Probes, Thermo Fisher Scientific) was applied before coverslipping with ProLong gold anti-fade reagent (Invitrogen). Control studies included omission of each primary antibody.

##### Lesion assessment and stereology

Triplicate sections from each NMDA infusion site were visualized using an Olympus microscope (Albertslund, Denmark) equipped with a 578–603 nm filter set for the detection of red fluorescence, at ×20 magnification. NMDA-induced lesions were identified by a distinct reduction or absence of NeuN fluorescence, which was analysed manually by tracing the site of injury using ImageJ software (NIH, Bethesda, MD, USA). Lesion volume was then determined as described by Osborne, *et al*. by integrating the cross-sectional area of damage between each stereotaxic infusion site and the distance between sites (Osborne et al., 1987). The number of NeuN positive cells within each lesion were also point counted using ImageJ software using a grid overlay to estimate the total number of NeuN positive cells within each region, and expressed as number of cells/mm^2^. Data obtained for infarct volume and the effects of treatment on neuronal counts were analysed by one-way ANOVA followed by the Bonferroni post-hoc test. Data were analysed using GraphPad Prism, version 6 (GraphPad Software Inc., San Diego) and presented as mean ± standard error of the mean (SEM). Statistical significance was defined as *p* <0.05.

##### Investigation of TAT-Src cell specificity in rat brains

Immunofluorescence within adjacent tissue sections was visualised using a fluorescent microscope equipped with a 578–603 nm filter set for detection of red fluorescence (NeuN, GFAP, OX-42), a 495–519 nm filter set for the detection of green fluorescence (FITC-Tat-Src), and a 478-495nm filter set for detection of blue fluorescence (DAPI) (ZeissAxioskop2, North Ryde, Australia). Immunohistochemical co-localisation of the stereotaxically infused FITC-Tat-Src was clearly identified in the cortex and striatum and was co-localised with the neuronal marker NeuN but not with the astrocytic marker GFAP or the microglial marker OX-42 one-hour post-infusion.

##### Infarct assessment

Absence, or reduction in NeuN immunoreactivity, revealed NMDA-induced lesions within the motor and parietal cortex as well as the striatum. Total lesion volume was consistent across treatment groups with no significant difference in volume of damage detected between groups (P >0.05, n=3/group, one-way ANOVA). Stereological point counting of NeuN positive cells within the lesion revealed treatment specific effects where the number of neurones in rats treated with TAT-Src was significantly greater than in rats receiving vehicle or scrambled Tat-Src control (P<0.0001, n=3/group, one-way ANOVA).

### METHODS DETAILS

#### Cell viability assays, related to Figure S1

##### MTT cell viability assay

Cell viability was determined from primary cortical neurons (seeded in 24-well plates) using the 3- (4,5-dimethylthiazole-2-yl)-2,5-diphenyltetrazolium bromide (MTT) assay. MTT stock solution (5 mg/ml (w/v) in sterile PBS) was diluted 1/100 in culture medium. At the end of treatment of cultured neurons, the culture medium was aspirated and replaced by the diluted MTT solution. After incubation for 2 h, the diluted MTT solution was removed and 200 µl DMSO was added per well to dissolve the formazan crystals formed. Absorbance at 570 nm was measured using the Clariostar Monochromator Microplate Reader (BMG Lab Technologies, Durham, NC). Cell viability was expressed as a percentage of the control cells.

##### LDH release assay of neuronal death

The activity of LDH released from the damaged neurons to the culture medium was measured. Briefly, 50 μl of culture medium from each well of the culture plate was transferred to 96 well-microtitre plates (Falcon). 100 μl of LDH assay mixture containing equal volume of LDH assay substrate solution, LDH Assay dye solution and LDH assay cofactor was then added to each well. The reaction was allowed to proceed at room temperature for 30 min in the dark and was stopped by adding 50 μl of 1 mM acetic acid. The absorbance at 490 nm of whole mixture was measured in triplicate the Clariostar Monochromator Microplate Reader (BMG Lab Technologies, Durham, NC). The release of LDH was calculated as a percentage of the untreated control.

#### Construction of spectral libraries of cultured cortical neurons and mouse brain tissues, related to Figures 1 and S2

Prior to quantitative analysis on the mass spectrometer, we built a spectral library, which contains all distinctive information of each identified peptides such as its retention time, charge state and the fragment ion information. The success of the quantitation largely depends on the peptides to be quantified being present in this spectral library (Ludwig et al., 2018). As such, we pooled lysates from neurons with and without glutamate (100 µM) treatment for 30 min and 240 min together with brain lysates of mouse stroke and traumatic brain injury models. Briefly, proteins in the lysates were precipitated with cold acetone (−20°C) and then resuspended in 8 M urea in 50 mM triethylammonium bicarbonate (TEAB). Proteins are then reduced with 10 mM tris(2-carboxyethyl)phosphine Hydrochloride (TCEP), alkylated with 55 mM iodoacetamide, and digested with trypsin (trypsin to protein ratio of 1:50 (w/w)) overnight at 37°C. The resultant tryptic peptides were purified by solid phase extraction (SPE) (Oasis HBL cartridge, Waters). For global proteome analysis, 100µg of these peptides were fractionated into 8 fractions using the high pH reversed-phase fractionation kit (Pierce) according to the manufacturer’s protocol before analysis on the Q-Exactive Orbitrap. For phosphoproteome analysis, phosphopeptides in the tryptic digest were enriched with a serial enrichment strategy (Thingholm et al., 2008). Briefly, the tryptic peptides after the SPE clean-up step were enriched for phosphopeptides in a serial fashion using both Fe-NTA and TiO_2_ based selective media. To increase the depth of coverage, the enriched phosphopeptides were fractionated into 8 fractions using the high pH reversed-phase fractionation kit before analysis on the Q-Exactive Orbitrap. Indexed Retention Time (iRT) peptides (Biognosys) were spiked into all samples prior to analysis.

The LC system coupled to the Q-Eaxctive Orbitrap mass spectrometer was equipped with an Acclaim Pepmap nano-trap column (Dinoex-C18, 100 Å, 75 µm x 2 cm) and an Acclaim Pepmap RSLC analytical column (Dinoex-C18, 100 Å, 75 µm × 50 cm). After pre-fractionation with the high pH reversed-phase fractionation kit, tryptic peptides in each of the 8 fractions were injected to the enrichment column at an isocratic flow of 5 µl/min of 2 % (v/v) CH_3_CN containing 0.1 % (v/v) formic acid for 6 min before the enrichment column was switched in-line with the analytical column. The eluents were 0.1 % (v/v) formic acid (Solvent A) and 100 % (v/v) CH_3_CN in 0.1 % (v/v) formic acid (Solvent B). The flow gradient was (i) 0 - 6 min at 3 % Solvent B, (ii) 6 - 95 min, 3 – 20 % Solvent B (iii) 95 - 105 min, 20 – 40 % Solvent B (iv) 105 - 110 min, 40 – 80 % Solvent B (v) 110 - 115 min, 80 – 80 % Solvent B (vi) 115 - 117 min 85 – 3 % Solvent B and equilibrated at 3% Solvent B for 10 minutes before the next sample injection. In the Data-Dependent Acquisition (DDA) mode, full MS1 spectra were acquired in positive mode, 70 000 resolution from 300-1650 *m/z*, AGC target of 3e^6^ and maximum IT time of 50 ms. Fifteen of the most intense peptide ions with charge states ≥ 2 and intensity threshold of 1.7e^4^ were isolated for MSMS. The isolation window was set at 1.2 m/z and precursors fragmented using normalized collision energy of 30, 17 500 resolution, AGC target of 1e^5^ and maximum IT time of 100 ms. Dynamic exclusion was set to be 30 sec. In the Data Independent Acquisition (DIA) mode, the separation gradient was identical to that for DDA analysis. The QExactive plus mass spectrometer was operated in the hyper reaction monitoring/data independent (HRM/DIA) mode, whereby full MS1 spectra were acquired in positive mode from 400 – 1000 *m/z*, 70 000 resolution, AGC target of 3e^6^ and maximum IT time of 50 ms. The isolation window was set at 21 *m/z* with a loop count of 30 and all precursors fragmented using normalized collision energy of 30, 17 500 resolution, AGC target of 1e^6^.

#### Analysis of global proteome, phosphoproteome and N-terminome of neurons

##### Quantitative Global and phospho-proteomic analysis, related to Figures 1, 2, S2, S3 and S4

Neuronal lysates (500 µg) were mixed with cold acetone (−20°C) (1:5, v/v) in microfuge tubes and incubated at −20°C overnight to precipitate proteins. Acetone precipitated proteins (in control and treated lysates) were resuspended in 8 M urea in 50 mM TEAB (pH 8.0), and protein estimation was carried out using BCA assay (Pierce-Thermo Scientific) according to manufacturer’s instruction. Equal amounts of protein were reduced with 10 mM tris-(2-carboxyethyl)-phosphine (TCEP) for 45 min at 37°C in a bench top vortex shaker. Reduced samples were alkylated with 55 mM iodoacetamide shaking 45 min at 37°C. Samples were diluted to 1 M urea (diluted with 25 mM TEAB) and digested with sequencing grade modified trypsin (1:50) overnight at 37°C. Digested samples were acidified to 1% (v/v) with pure formic acid, and solid phase extraction (SPE) was carried out with 60 mg Oasis HBL cartridge (Waters) to clean up the digested peptides. Briefly, the cartridge was washed with 80% acetonitrile (ACN) containing 0.1% trifluoroacetic acid (TFA) first and then with only 0.1% TFA before sample loading. Samples were washed again with 0.1% TFA and eluted with 800 µl 80% ACN containing 0.1% TFA. An aliquot (20 µg) of eluted peptides were freeze-dried overnight prior to analysis of changes in global proteome. For quantitative global proteomic analysis, 1 µg peptide in the presence of spiked-in iRT peptide was injected into the mass spectrometer and analysed using the HRM/DIA mode followed by analysis with the Spectronaut DIA-MS methodology and making use of the global proteome-specific spectral library built with the SEQUEST search engine incorporated in Proteome Discover (PD) (Wang et al., 2020) (Figure S2).

For quantitative phosphoproteome analysis, phosphopeptides in the remaining tryptic digest after the SPE clean-up step (470 µg) were enriched in a serial fashion using both Fe-NTA and TiO_2_ based selective media as described by Thingholm, *et al*. (Thingholm et al., 2008). The enriched phosphopeptides using this method was injected into the Q-Exactive Orbitrap in the presence of spike-in iRT peptides and using the HRM/DIA mode followed by analysis using the Spectronaut DIA-MS methodology. To monitor the changes of phosphoproteome in neurons induced by glutamate treatment, we first used two different search engines including Andromeda incorporated in MaxQuant (MQ) and SEQUEST to construct phosphopeptide-specific spectral libraries (Figure S2). We then conducted Spectronaut DIA-MS analysis using these libraries to identify and quantify changes in the phosphorylation of neuronal proteins induced by the treatment.

##### Analysis of the changes in neuronal N-terminome during excitotoxicity by the <u>T</u>erminal <u>A</u>mine Isotopic labelling of <u>S</u>ubstrates (TAILS) method, related to Figures 3, S5, S9 and S11

The workflow of TAILS analysis we adapted for our study is shown in Figure S1. In brief, neurons were treated with the neurotoxic concentration of glutamate (100 µM) for 30 min and 240 min. This toxic treatment strategy has been known to induce enhanced limited proteolysis (referred to as proteolytic processing) as well as degradation of specific neuronal proteins by specific proteases activated in response to glutamate over-stimulation (referred to as excitotoxicity-activated proteases) (El-Gebali et al., 2019; Hossain et al., 2013). Proteins in the cell lysates of control (untreated) neurons and glutamate-treated neurons were precipitated by ice-cold acetone. After resuspension and denaturation in 8 M guanidinium hydrochloride in 100 mM HEPES, the neuronal proteins were reduced and alkylated. This is followed by isotopic dimethyl labelling of the free amino groups including the N^α^-amino groups and ε-amino groups of the lysine side chains in the neuronal proteins. Proteins of the control neurons were labelled with formaldehyde (CH_2_O) (referred to as light dimethyl labelled), while those of the glutamate-treated neurons were labelled with deuterated formaldehyde (CD_2_O) (referred to as medium dimethyl labelled). Thus, the neo-N-termini of truncated protein fragments generated by proteolysis catalyzed by the excitotoxicity-activated proteases were medium dimethyl labelled (i.e., covalently linked with deuterated dimethyl groups). The light dimethyl labelled proteins from control neurons and the medium dimethyl labelled proteins from treated neurons were mixed at a ratio of 1:1 (w/w). The protein mixture was subjected to tryptic digestion. Tryptic peptides derived from the N-terminal end of neuronal proteins were devoid of free amino groups because the amino groups were either naturally blocked by modifications such as acetylation and myristoylation *in vivo* or by dimethylation *in vitro*. The other tryptic peptides derived from other parts of neuronal proteins contained the newly formed free N^α^-amino group resulting from trypsinization. These peptides were selectively captured by the Hydroxy Polyglycerol Aldehyde (HPG-ALD) Polymer by reaction of their N^α^-amino groups with the aldehyde (-CHO) groups of the polymer. After ultrafiltration to remove the complex of tryptic peptide-bound HPG-ALD polymers from the naturally modified and dimethyl labelled N-terminal peptides. The modified N-terminal peptides from each sample were pre-fractionated into 4 fractions SDB-RPS (styrene-divinylbenzene reverse phase sulfonate) based fractionation prior to LC-MS/MS analysis on the Orbitrap Elite mass spectrometer.

LC-MS/MS was carried out on LTQ Orbitrap Elite (Thermo Scientific) with a nanoESI interface in conjunction with an Ultimate 3000 RSLC nano HPLC (Dionex Ultimate 3000). The LC system was equipped with an Acclaim Pepmap nano-trap column (Dionex-C18, 100 Å, 75 µm x 2 cm) and an Acclaim Pepmap RSLC analytical column (Dionex-C18, 100 Å, 75 µm x 50 cm). The tryptic peptides were injected to the enrichment column at an isocratic flow of 5 µL/min of 3% v/v CH_3_CN containing 0.1% v/v formic acid for 5 min before the enrichment column was switched in-line with the analytical column. The eluents were 0.1% v/v formic acid (solvent A) and 100% v/v CH_3_CN in 0.1% v/v formic acid (solvent B). The flow gradient was (i) 0-6 min at 3% B, (ii) 6-95 min, 3-20% B (iii) 95-105 min, 20-40% B (iv) 105-110 min, 40-80% B (v) 110-115 min, 80-80% B (vi) 115-117 min 85-3% and equilibrated at 3% B for 10 minutes before the next sample injection. The LTQ Orbitrap Elite spectrometer was operated in the data-dependent mode with nanoESI spray voltage of 1.8kV, capillary temperature of 250°C and S-lens RF value of 55%. All spectra were acquired in positive mode with full scan MS spectra from *m/z* 300-1650 in the FT mode at 240,000 resolution. Automated gain control was set to a target value of 1.0e6. Lock mass of 445.120025 was used. The top 20 most intense precursors were subjected to rapid collision induced dissociation (rCID) with the normalized collision energy of 30 and activation q of 0.25. Dynamic exclusion with of 30 seconds was applied for repeated precursors.

#### Quantification and statistical analysis of proteomics data

##### Quantitation of global and phosphoproteome changes in mouse primary cortical neurons induced by glutamate treatment, related to Figures 1, 2, S3 and S4

Protein/peptide identification from the DDA-based analysis and subsequent spectral library generation were conducted using Proteome Discoverer (v.2.1, Thermo Fischer Scientific) with the Sequest HT search engine in combination with the Percolator semi-supervised learning algorithm (Kall et al., 2007) and PhosphoRS (Taus et al., 2011) or with MaxQuant software utilizing the Andromeda search engine (Tyanova et al., 2016) on M*us musculus* protein database (SwissProt (TaxID = 10090)(version 2017-10-25)) (Figure S2). The search parameters were: a precursor tolerance of 20 ppm, MSMS tolerance of 0.05 Da, fixed modifications of carbamidomethylation of <u>cysteine</u> (+57 Da), <u>methionine</u> oxidation (+16 Da) or phosphorylation of serine, threonine and tyrosine (+80Da) . Peptides were accepted based on a false discovery rate (FDR) of <0.01 at both the <u>peptide and protein</u> levels.

DIA based quantitative analysis was carried out using the Spectronaut software (Spectronaut 11, Biognosys). To build the spectral library in Spectronaut based on MaxQuant and Proteome Discoverer results, all the default parameters were used except that best N-terminal fragments per peptide was set to minimum 5 and maximum 20. Two spectral libraries were built for phosphopeptides whereas one library from the search results of Proteome Discoverer was built for the global library. The spectral libraries for both global and phosphopeptides are generated identically (Figure S2). The DIA files for each sample were converted to htrms format sing HTRMS Converter (Biognosy) and loaded on Spectronaut 11 for generation of protein and phosphopeptide intensities. The default parameters were used for data extraction, XIC extraction, calibration, identification, and protein inference. The iRT profiling strategy was used with unify peptide peak options enabled. For quantitation of both global and phosphopeptides the Q value percentile of 0.75 was set. For global analysis, quantitation is based on stripped sequence, and global normalization was performed. For phosphopeptide quantitation, the modified sequence with local normalization was used. The Spectronaut results output were further processed with Perseus software (version 1.6) (Tyanova and Cox, 2018). For phosphoproteome analysis, only the set of phosphopeptides identified with exact site and charge states matched in two spectral libraries (built from MaxQuant search and Proteome Discoverer), were used for analysis in Perseus. The reproducibility of the DIA phosphopeptide quantitation was evaluated by calculating Pearson’s correlation coefficients and visualised in the heatmap. To identify the phosphopeptides that were significantly changed two-sample student’s t-tests were performed and cut-offs were applied as follows: p-value ≤ 0.01, FDR 1% and fold change ± 2. These significantly regulated phosphopeptides were exported to Excel and processed for bioinformatic analysis.

##### Classification of the N-terminal peptides identified and quantified by TAILS method, related to Figures 3, S5 and S10

Using the TAILS method (Kleifeld et al., 2011), we identified and quantified (i) the “natural” free N-termini of mature proteins biosynthesized in physiological conditions, and (ii) the newly formed neo-N-termini of stable truncated protein fragments and intermediate peptide fragments generated by proteolysis of neuronal proteins during excitotoxicity. Of the quantifiable N-terminal peptides identified by the TAILS analysis of excitotoxic neurons, over 2,300 contained acetylated N-termini and over 2,600 contained dimethyl labelled N-termini (Figures 1C and S7, Tables S6A and S7A). Whereas the acetylated N-terminal peptides were derived from intact proteins undergoing N-terminal acetylation during biosynthesis and maturation in neurons, the dimethyl-labelled N-terminal peptides were derived from free N-termini of neuronal proteins *in vitro* via the TAILS procedure. The dimethyl labelled N-terminal peptides were further sub-divided into four groups: (i) those retaining the first methionine encoded by the start codon (+ Met); (ii) those with the start codon-encoded methionine removed (-Met); (iii) those with the N-terminal signal peptide segments removed during maturation of the parental neuronal proteins *in vivo* (-signal peptide); and (iv) those containing neo-N-termini generated by proteolysis of intact mature neuronal proteins (Neo) (Figure 1C). We focused our further analysis on the neo-N-terminal peptides as they revealed both the cleavage sites and identities of neuronal proteins undergoing proteolysis during excitotoxicity. Based on the rationale depicted in Figure S6, neo-N-terminal peptides exhibiting increased abundance were assigned as those derived from the stable truncated protein fragments generated from enhanced proteolytic processing of neuronal proteins during excitotoxicity. Neo-N-terminal peptides exhibiting decreased abundance were assigned as being derived from neuronal proteins undergoing enhanced degradation for clearance during excitotoxicity.

##### Determination of the abundance ratio cut-off values to identify neuronal proteins exhibiting significantly enhanced proteolysis induced by treatment with glutamate or co-treatment with glutamate and calpeptin, related to Figures 3 and S10

Over 2,000 neo-N-terminal peptides were found by the TAILS method to be generated by enhanced proteolysis during excitotoxicity (Figures 1C and 3). Figure 3A depicts results of our statistical analysis to calculate the thresholds for their assignment to be neo-N-terminal peptides generated by significantly enhanced degradation and those generated by significantly enhanced proteolytic processing of neuronal proteins during excitotoxicity. Based upon the normalized distribution of the log_2_ M/L ratios of the identified neo-N-terminal peptides, we determined the median and standard deviations (S.D.) of the distribution of the log_2_ M/L ratios. Statistically, values that are outside the 1.5 interquartile range (IQR) are considered as outliers (Tukey, 1977) (Figure 3A, upper panels). The 1.5×IQR values can be calculated as 5 × S.D. of the distributions of the abundance ratios (log_2_ M/L ratios) of all quantifiable N-terminal peptides (Figure 3A, lower panels). Using the Tukey’s 1.5 × interquartile range (IQR) rule (Tukey, 1977), we determined the abundance ratio cut-off values (M/L) for the neo-N-terminal peptides exhibiting significantly reduced abundance and those exhibiting significantly increased abundance during excitotoxicity (Figure 3A, lower panels).

To determine the abundance ratio cut-off values to identify the neo-N-terminal peptides generated by significant proteolysis by neuronal proteases activated in response to co-treatment with glutamate and calpeptin, the M/L ratio cut-off values equivalent to 1.5 interquartile range (IQR) were calculated. As depicted in Figure S10, for the N-terminal peptides co-treated for 30 min, the lower and upper cut-off values were 0.271 and 3.8 (i.e. M/L ≥ 3.8 for the neo-N-terminal peptides generated by significantly enhanced proteolytic processing and M/L ≤ 0.271 for those generated by significantly enhanced degradation). For those derived from neurons co-treated for 240 min, the lower and upper cut-off values were 0.386 and 2.876, respectively.

#### Synthesis of FITC-Tat-Src peptide, related to Figure 7

Peptides were constructed on a CEM Liberty 12-Channel Automated Microwave Peptide Synthesizer using Fmoc-PAL-PEG-PS^TM^ (Rink resin; loading capacity 0.21 mmol/g) (ThermoFisher, Cat.#: GEN913383) and Fmoc-protected amino acids (GL Biochem (Shanghai)). Fmoc deprotections were performed using 20% piperidine in DMF. Activation of Fmoc-amino acids was achieved using a 0.5 M solution of HCTU (2-(6-chloro-1H-benzotriazole-1-yl)-1,1,3,3-tetramethylaminium hexafluorophosphate) and DIPEA (diisopropylethylamine) in DMF in a ratio of 2 ml:2 ml:0.34 ml per 1 mmol of Fmoc amino acid used. Peptide coupling and deprotection efficiency was monitored using a 2,4,6-trinitrobenzene-sulfonic acid (TNBSA) assay (Hancock and Battersby, 1976). The FITC-Tat-ahx-Src 49-79 peptide (FITC-Tat-Src) was synthesized sequentially as two main parts with aminohexanoic acid linker used between the two main sequences. First, the sequence ahx-Src 49-79 was synthesized. The synthesized peptide was validated via ESI-MS and RP-HPLC. Then, the Tat peptide with sequence ahx-GRKKRRQRRRPQ was continued on the ahx-Src 49-79 sequence while still on resin to generate ahx-Tat-ahx-Src 49-79 peptide-resin. Fluorescein isothiocyanate (FITC) was coupled to ahx-Tat-ahx-Src 49-79 peptide-resin using HCTU and DIPEA. The peptide was cleaved from the resin using TFA/triisopropylsilane/H_2_O mixture (volume ratio of 95:2.5:2.5) for 90 min. Excess TFA was removed via evaporation with stream of N_2_ gas. The peptide was precipitated by addition of diethyl ether. The mixture was then centrifuged and the ether decanted. The pellet containing the peptide was redissolved in 30% acetonitrile/H_2_O and filtered through a 0.22 μm filter. The crude peptide solution was lyophilized prior to purification by semi-preparative RP-HPLC. Fractions containing the pure peptide were pooled and lyophilized. The dried peptide was stored at 4 °C until further use. The purified peptide was analyzed and validated via ESI-MS with mass [M + H^+^] of 5385.4 Da and RP-HPLC shown as a single peak in HPLC chromatogram.

## SUPPLEMENTAL INFORMATION

Supplemental information includes fourteen figures and ten tables can be found with this article online. The mass spectrometry proteomics data have been deposited to the ProteomeXchange Consortium via the PRIDE (Perez-Riverol et al., 2019) partner repository with the dataset identifiers PXD019211 for the project entitled “N-terminomics Analysis of Mouse cultured cortical neuron treated with glutamate”, and PXD019527 for the project entitled “Phosphoproteomics of mouse cortical neuron”.

## AUTHOR CONTRIBUTIONS

S.S.A. and C.-S.A. conceptualized, designed, performed, and analyzed most of the experiments and co-wrote the paper. H.-C.C., N.A.W., A.D., O.K., G.D.C. and C.R. conceptualized, designed and analyzed the proteomic data, neuronal cell biology data or data of in vivo model of neurotoxicity. M.I.H., M.A.K., H.-J.Z., G.D.C., A.H., L.B. and C.R. conceptualized, designed and performed the experiments of excitotoxic cell death of cultured neurons and effects of Tat-Src and Tat-Scrambled peptides on neuronal loss in vivo. S.S.A., H.-C.C., C.-S.A., N.A.W., S.S., H.N., R.M. and D.D. performed experiments to generate the spectral libraries for DIA proteomic analysis of the changes in global and phospho-proteomes of neurons. M.L., H.-C.C., A.D., O.K. N.A.W. and C.-S.A. analyzed the N-terminomic data to define the potential substrates of calpains in neurons. D.L., H.-J.Z., A.D., I.S.L. and J.P.L. conceptualized, interpreted experimental data on excitotoxic neuronal death and experimental validation of the proteomic results and co-wrote the paper.

## ACKNOWLEDGEMENTS

This work was funded by grants to H.-C.C., H.N., R.M., I.S.L., A. Dhillon and A. Dufour. from the National Health and Medical Research Council (NHMRC) of Australia and NSERC of Canada (NHMRC project grant #1050486 to H.-C.C.; NHMRC project grant #1141906 to A. Dhillon; NSERC discovery grant #DGECR-2019-0012). O.K is supported by grants from the Israel Science Foundation (1623/17 and 2167/17). We thank Robert Qi, Prasad Paradkar, Swati Varshney and Anderly Chueh for comments and suggestions for drafting this manuscript.

## SUPPLEMENTARY FIGURES

**Figure S1.**
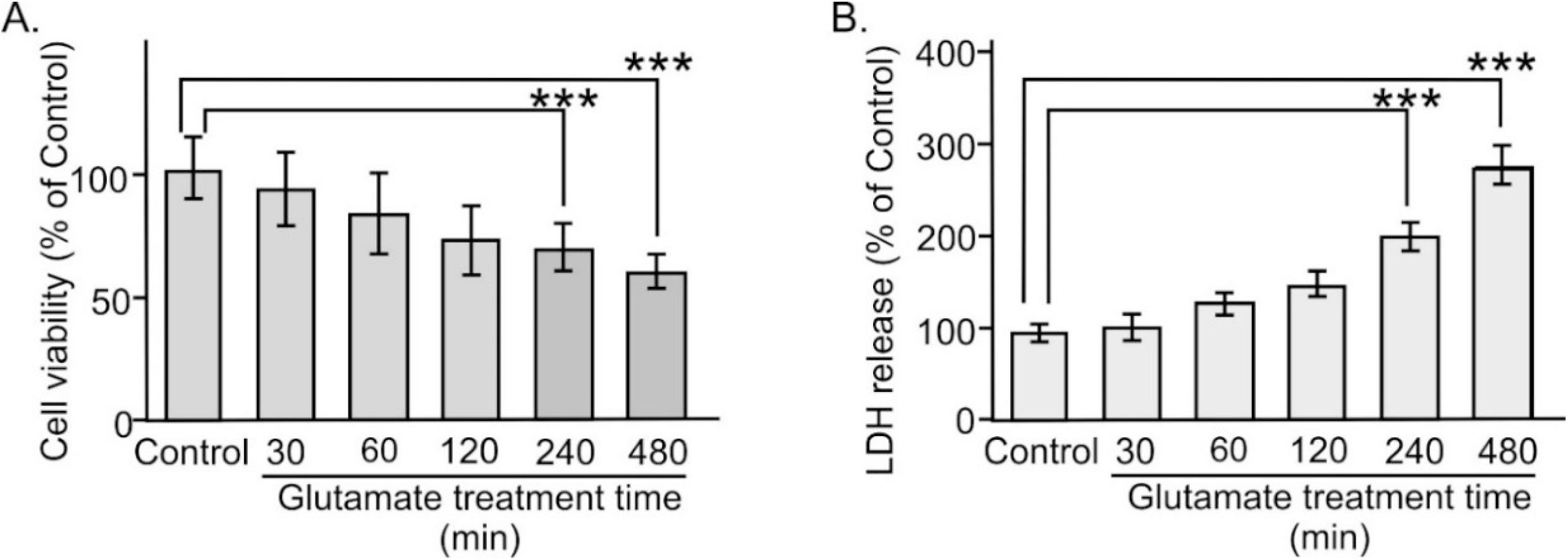
Viability of cultured cortical neurons after different duration of glutamate treatment, related to Figure 1. **(A)** MTT assay results showing neuronal viability at different time points after glutamate treatment. The amount of formazon formed by the viable neurons at each time point of treatment is presented as the percentage of that formed by the viable untreated neurons (Control). Data represent as mean ± S.D. (error bars), n = 6; *** represents *p <* 0.005, Student’s *t-*test. **(B)** The amount of LDH released from the damaged neurons at each designated treatment time point is presented as the percentage of that released by the untreated neurons (Control). Data represent as mean ± S.D. (error bar), n = 3; *** represents *p* < 0.005, Student’s *t*-test.

**Figure S2.**
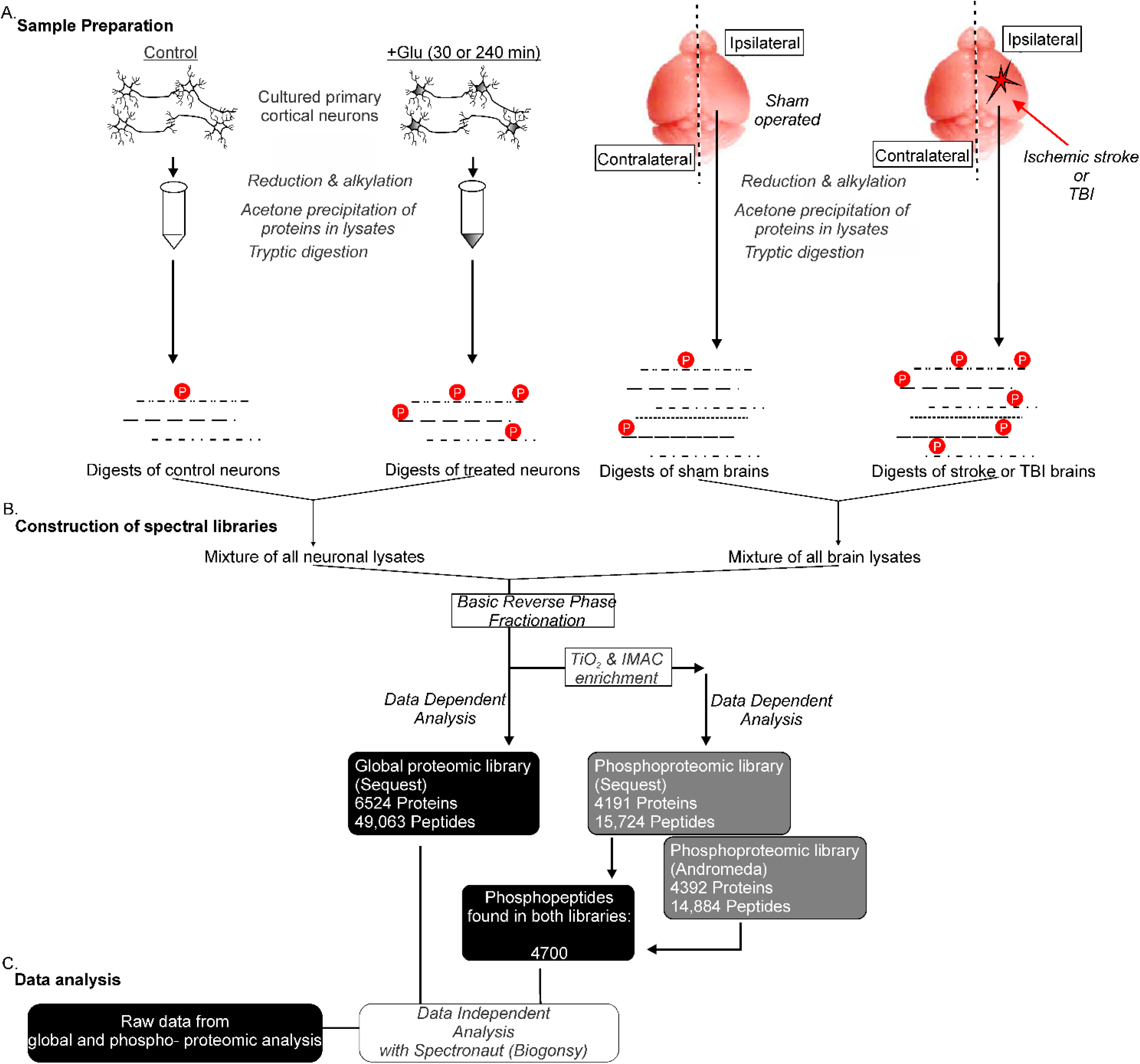
Workflow for the construction of spectral libraries from lysates of mouse neurons and brain tissues, related to Figure 1. **(A)** Lysates of untreated (Control) and glutamate-treated neurons were pooled and combined with lysates of brain tissues from sham operated mice and from mouse models of ischemic stroke and traumatic injury (TBI). Proteins in the combined cell and tissue lysates were precipitated, resuspended, reduced and alkylated prior to tryptic digestion. **(B)** The resulting tryptic peptides were purified by solid phase extraction (SPE) (Oasis HBL cartridge, Waters) followed by fractionation with basic reverse-phase column chromatography. An aliquot consisting of 20 μg of peptides in each fraction were subjected to LC-MS/MS for the construction of the spectral library for global proteomic analysis. Phosphopeptides were enriched from the remaining portion of peptides in each fraction by TiO_2_ and Fe^3+^-NTA affinity columns, prior to LC-MS/MS with the QE plus Orbitrap mass spectrometer. The identified peptides and phosphopeptides were used for construction of the spectral libraries. **(C)** Data independent analysis to profile changes in global proteome and phosphoproteome of cortical neurons induced by glutamate treatment.

**Figure S3.**
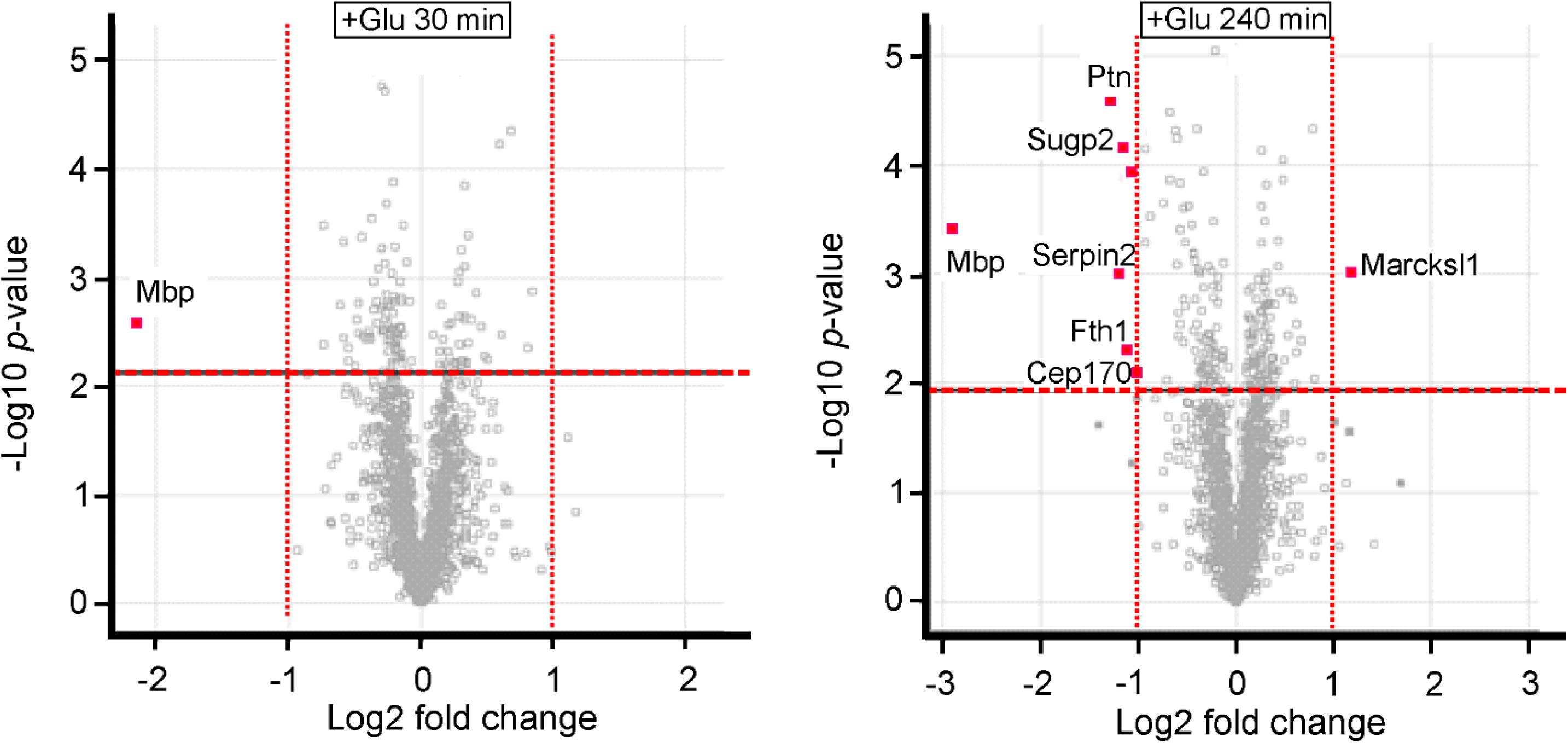
Neuronal proteins exhibiting significant changes in abundance at 30 min and 240 min after glutamate treatment revealed by global proteomic analysis, related to Figure 1. Volcano plots showing abundance ratios of the identified proteins in glutamate-treated versus Control neurons. The red dotted line in each plot indicates the threshold of false discovery rate (FDR) ≤ 5% (or 0.05) in the two-sample t-test. The fold change for each protein was calculated as log_2_ (Treatment/Control abundance) ratio. Red dots depict proteins exhibiting significantly changed abundance (presented as Uniprot accession numbers) with FDR ≤ 0.05 and Treated/Control abundance ratio ≥ 2 or ≤ 0.5. The identified proteins that did not exhibit significant abundance change or with FDR ≥ 0.05 are represented as grey coloured square boxes.

**Figure S4.**
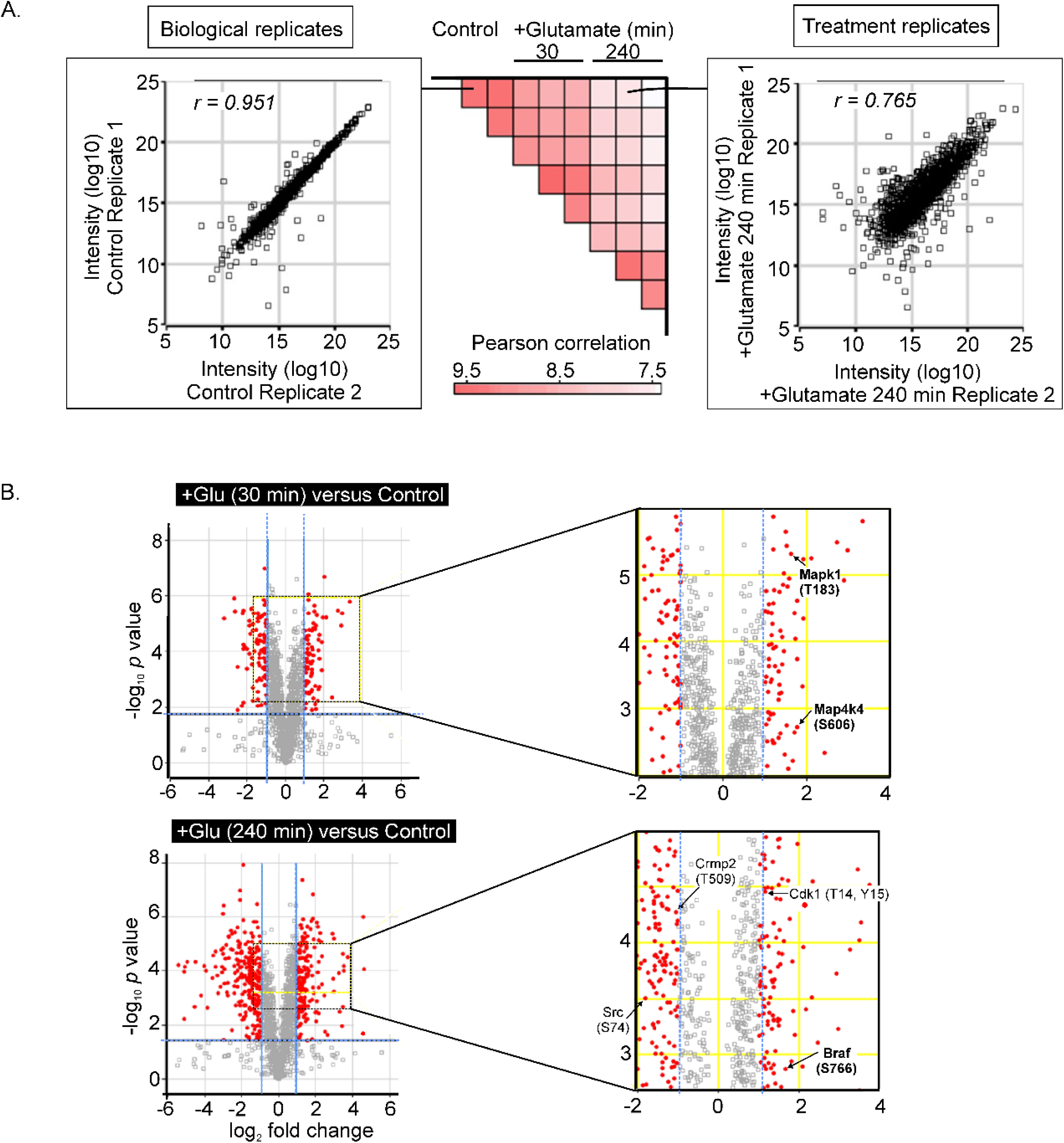
Heat maps, scatter plots and volcano plots of phosphoproteomic changes, related to Figure 2. **(A)** Heat maps and scatter plots showing the correlation between replicates of phosphoproteomic changes at different time points. *Middle Panel*: Heat map of Pearson correlation coefficients between replicates of the treatment conditions, untreated (control), glutamate treatment for 30 min and glutamate treatment for 240 min; n = 3. *Left panel*: Scatter plot showing the correlation between two selected biological replicates in Control. Right panel: Scatter plot showing the correlation between one control replicate and one replicate of the +Glu 240 neurons. **(B)** Volcano plots showing phosphopeptides undergoing significant changes in abundance upon glutamate treatment. Two sample t-test was performed in Perseus software (version 1.6) to calculate the fold change of each identified phosphopeptide induced by glutamate overstimulation at 30 min and 240 min. The abundance of the phosphopeptides in the untreated neurons (control) was considered as the reference point. The abundance of phosphopeptides derived from the control and neurons treated with glutamate at 30 min and 240 min are presented in volcano plots. Red dots: phosphopeptides with ± 2-fold changes (thresholds indicated by vertical blue lines) induced by glutamate treatment and identified with a false discovery rate (FDR or *p*-value) < 0.01 in the two-sample *t*-test. Grey squares: phosphopeptides not exhibiting significant changes in glutamate over-stimulation or identified with FDR > 1%. Symbols (red dots and grey squares) above the blue horizontal lines are phosphopeptides identified with FDR ≤ 1%. Log_2_ fold change: log_2_ abundance ratio of each phosphopeptide in treated versus control. *Right panels*: close-up views of the portions in the volcano plots with labels of the selected significantly changed phosphopeptides and the corresponding phosphosites.

**Figure S5.**
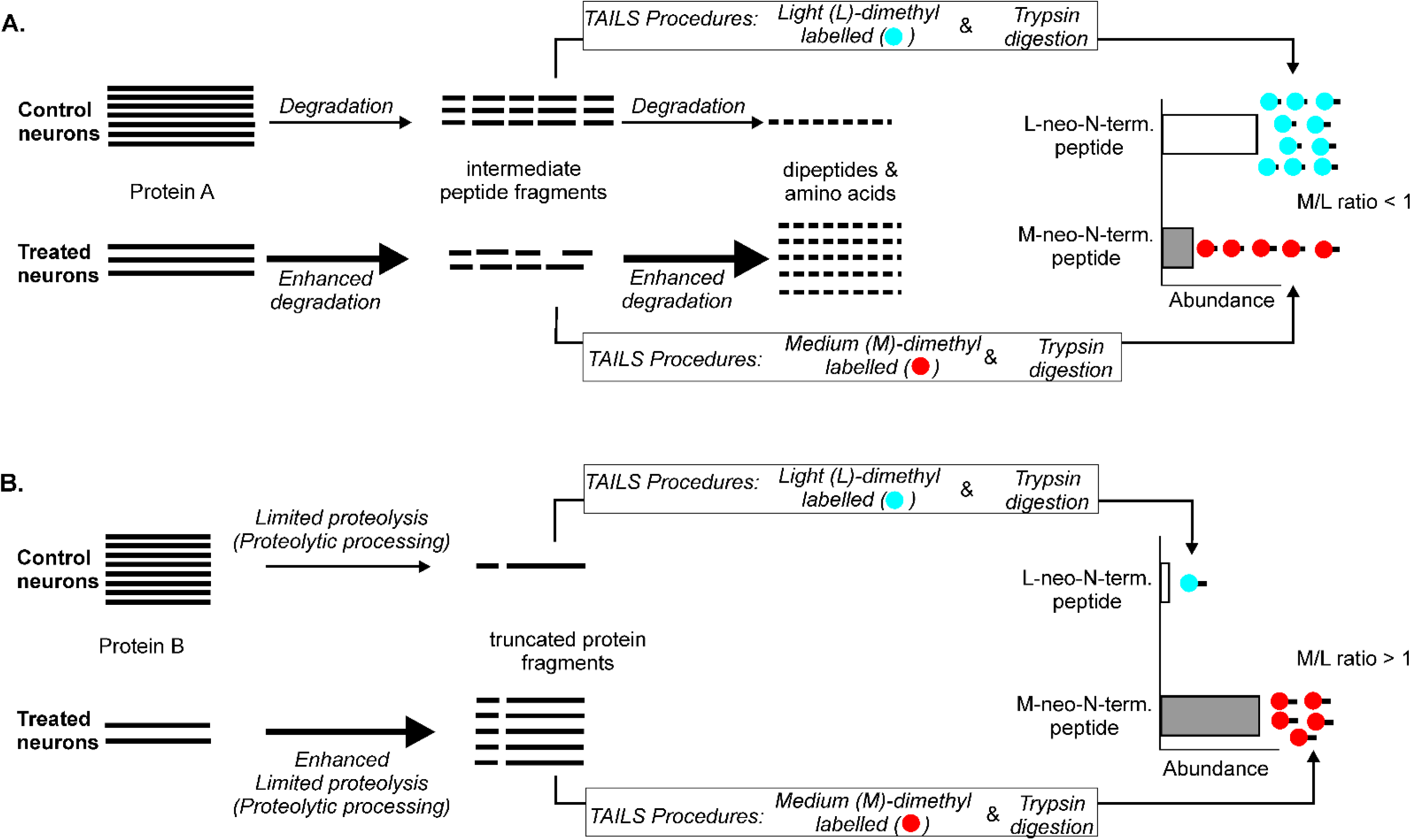
Rationale for assignment of neo-N-terminal peptides generated by enhanced degradation and those generated by enhanced proteolytic processing, related to Figure 3. **(A)** The treatment induces Protein A to undergo enhanced degradation for clearance. It is first proteolyzed to form intermediate peptide fragments, which are then further degraded to form dipeptides and amino acids. In the TAILS procedures, the intermediate peptide fragments are isotopically dimethyl labelled at their neo-N-termini, followed by tryptic digestion to generate the isotopically labelled neo-N-terminal peptides. Since the treatment induces enhanced degradation of the intermediate peptide fragments, the neo-N-terminal peptides derived from Protein A are less abundant in the treated neurons than those in the control neurons (i.e. M/L ratio <1). **(B)** The treatment induces Protein B to undergo enhanced limited proteolysis (proteolytic processing) to form stable truncated protein fragments. Upon isotopic dimethyl labelling of the neo-N-termini of the truncated protein fragments and tryptic digestion in the TAILS procedures, the resultant labelled neo-N-terminal peptides derived from Protein B are more abundant in the treated neurons than those in the control neurons (i.e. M/L ratio <1).

**Figure S6.**
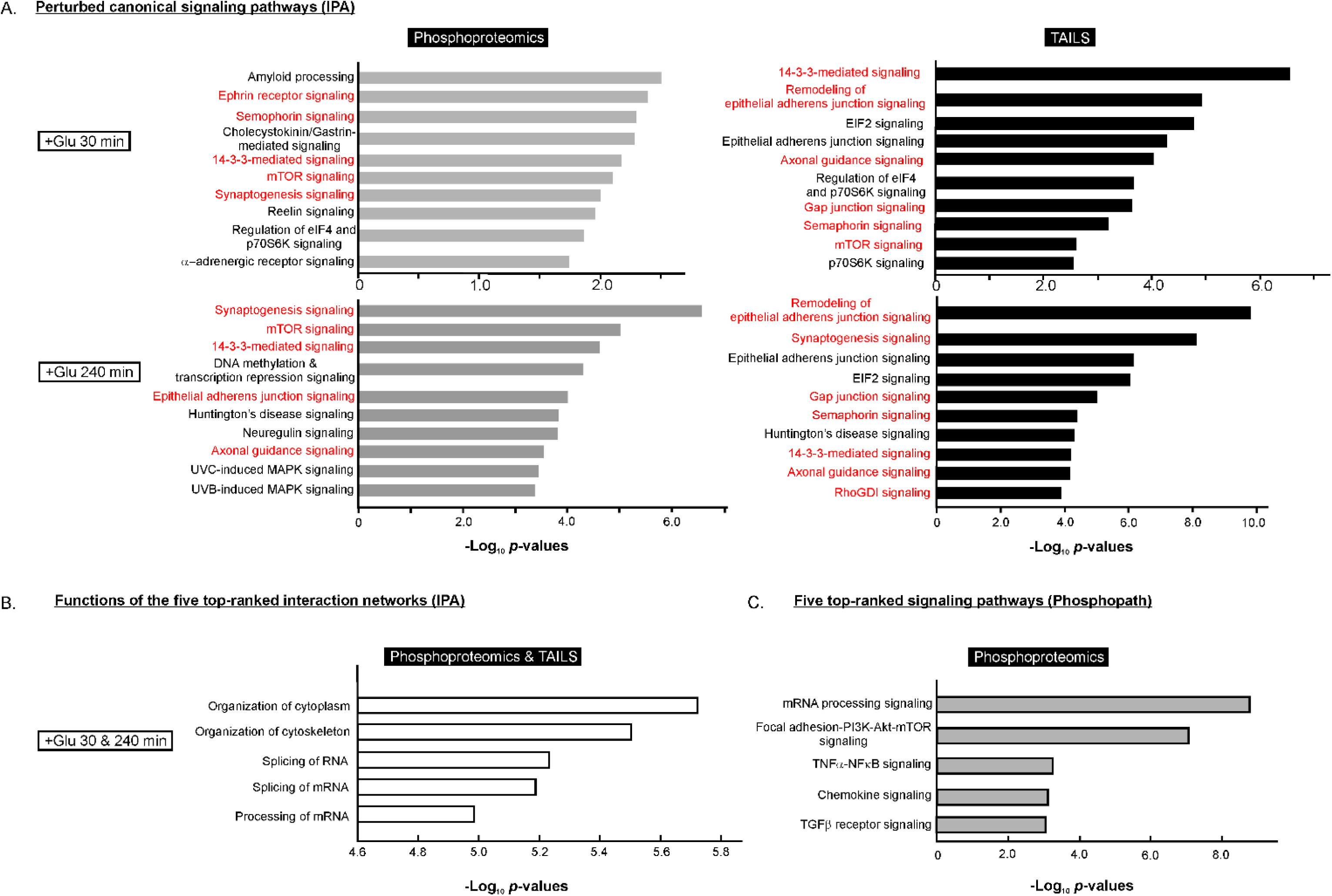
Canonical signalling pathways and cellular functions in neurons perturbed by glutamate treatment, related to Figures 4 and S7. **(A)** The top-ranked perturbed neuronal signalling pathways in response to glutamate treatment revealed by analysis of the changes in phosphoproteome and N-terminome determined by TAILS with the Ingenuity Pathway Analysis (IPA) software. The minimum threshold for *p-*value was set to < 0.01. The components of the pathways in red fonts are presented in Figure 4 of the main text. **(B)** Functions of the five top-ranked interaction networks formed by neuronal proteins with altered phosphorylation and those undergoing enhanced proteolysis during excitotoxicity as revealed by analysis with the IPA software. **(C)** Top-ranked phosphosite-enriched signalling pathways perturbed in neurons during excitotoxicity predicted by analysis of proteomics data with PhosphoPATH App. The interaction networks harbouring these pathways are shown in Figure 4A.

**Figure S7.**
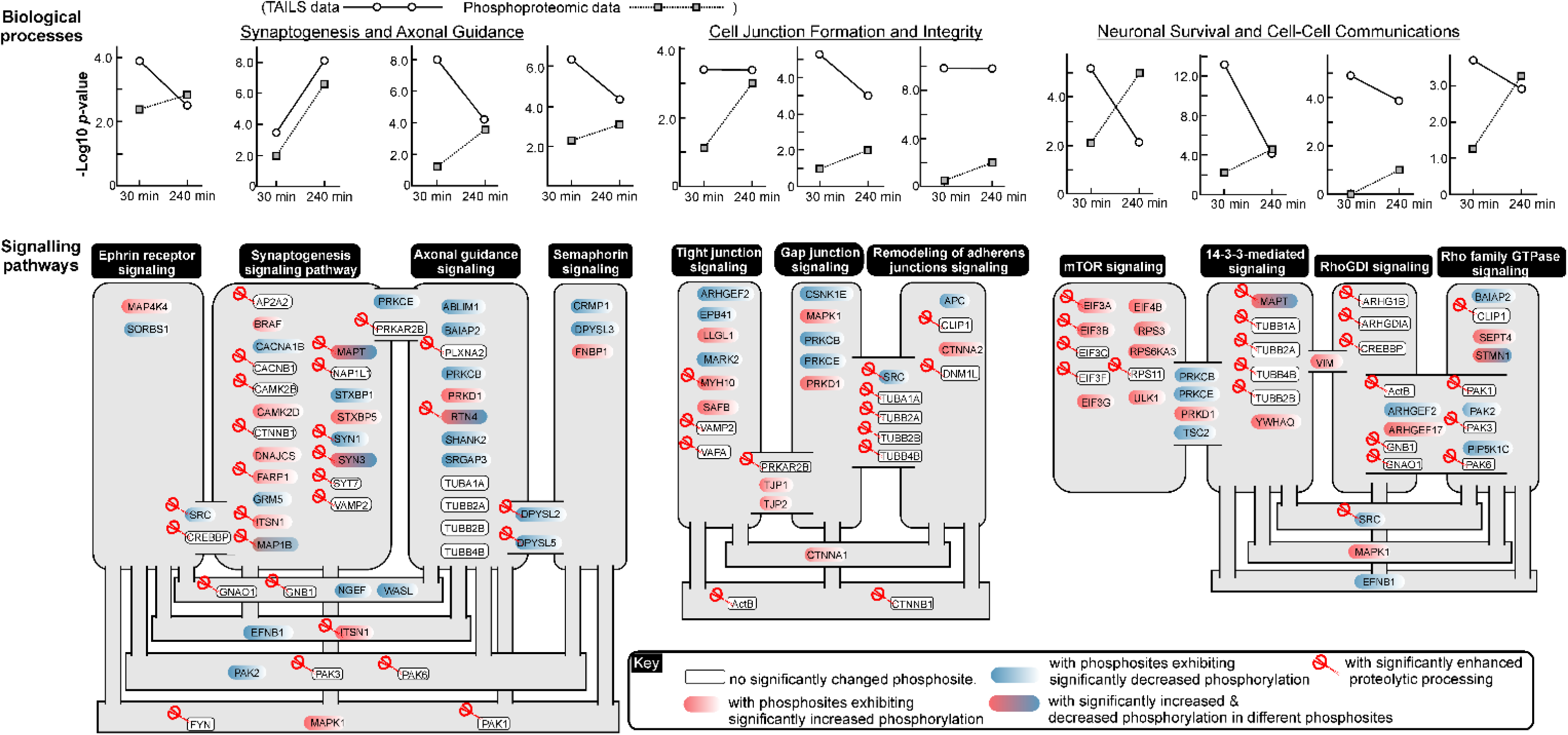
Major neuronal signalling pathways perturbed during excitotoxicity as revealed by TAILS and phosphoproteomic analyses, related to Figures S6A and S6B. Ingenuity pathway analysis (IPA) of neuronal proteins undergoing significant changes in phosphorylation and/or proteolytic processing in glutamate-treated neurons to identify the biological processes and signalling pathways in neurons perturbed by glutamate treatment. The top-ranked significantly perturbed biological processes are: (i) synaptogenesis and axonal guidance, (ii) cell junction formation and integrity relevant to synaptogenesis and axonal guidance, and (iii) neuronal survival and cell-cell communications. The plots indicate changes in degree of significance (–log10 p-values) of perturbation of the associated top-ranked signalling pathways at both treatment time points revealed by the TAILS and phosphoproteomic analysis. Neuronal proteins participating in the perturbed signalling pathways are shown in grey boxes. Those participating in multiple perturbed signalling pathways are shown in the grey connecting text boxes. Inset: key of the symbols. Red scissors: proteins undergoing significant proteolytic processing. White boxes: proteins did not show detectable changes in phosphorylation state. Red boxes: proteins with at least one phosphosite showing increased phosphorylation; Blue boxes: proteins with at least one phosphosite showing decreased phosphorylation. Red and blue boxes: proteins with at least one phosphosite showing decreased phosphorylation as well as one showing increased phosphorylation.

**Figure S8.**
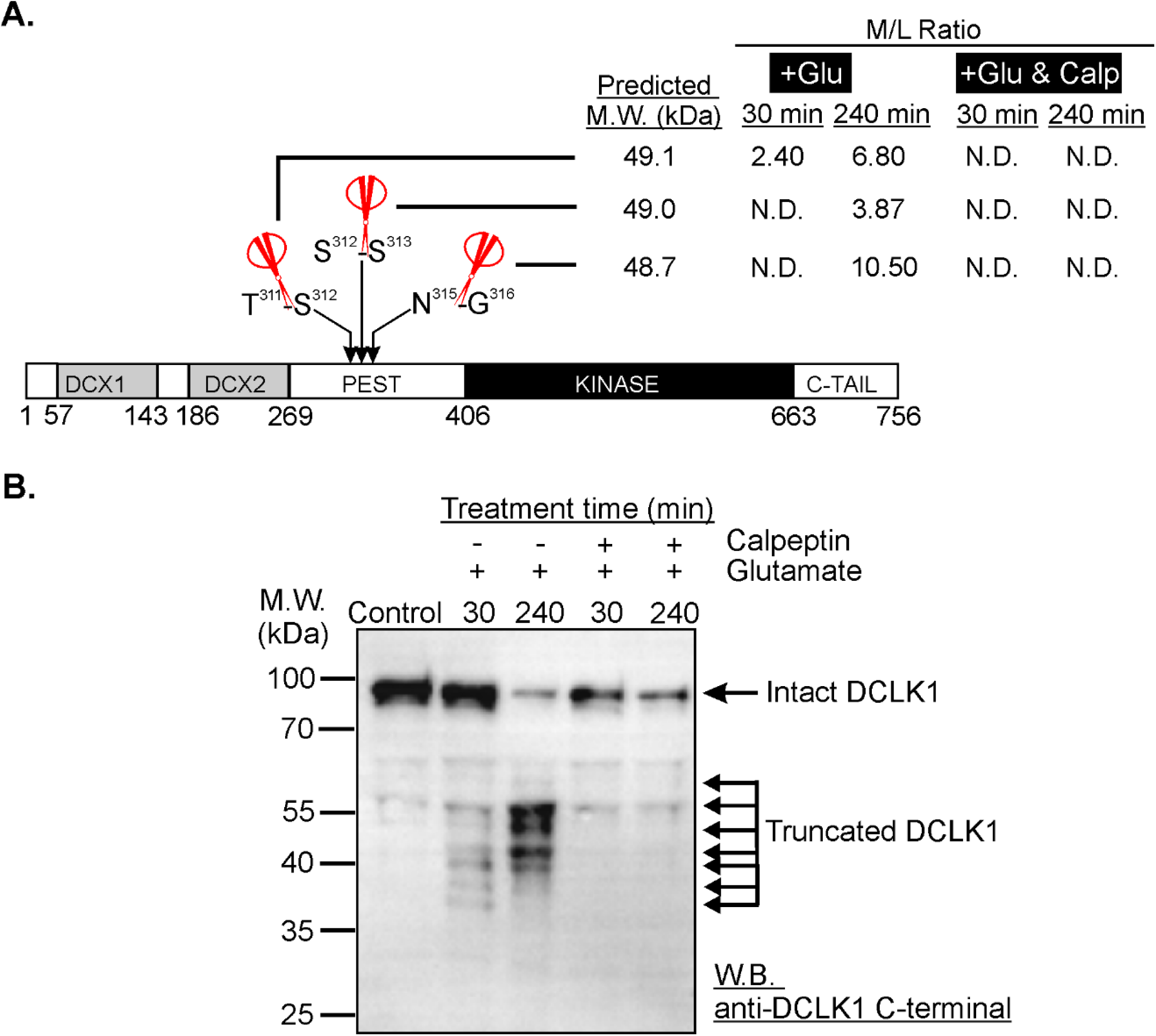
Calpeptin abolished significant internal processing of most neuronal proteins such as DCLK1 induced by glutamate treatment, related to Figure 6. **(A)** The cleavage sites in DCLK1 identified by TAILS. Red scissors: cleavage sites of significantly enhanced proteolytic processing during excitotoxicity. DCX1 and DCX2: doublecortin domains 1 and 2. PEST: sequence rich in proline, glutamate, serine and threonine. KINASE: protein kinase domain. C-TAIL: C- terminal terminal. **(B)** Western blot of lysates from untreated (Control), glutamate-treated, and glutamate/calpeptin co-treated neurons probed with the anti-C-terminal DCLK1 antibody. Some of the truncated DCLK1 fragments were likely generated by cleavage at sites of significantly enhanced proteolytic processing identified by TAILS shown in panel A.

**Figure S9.**
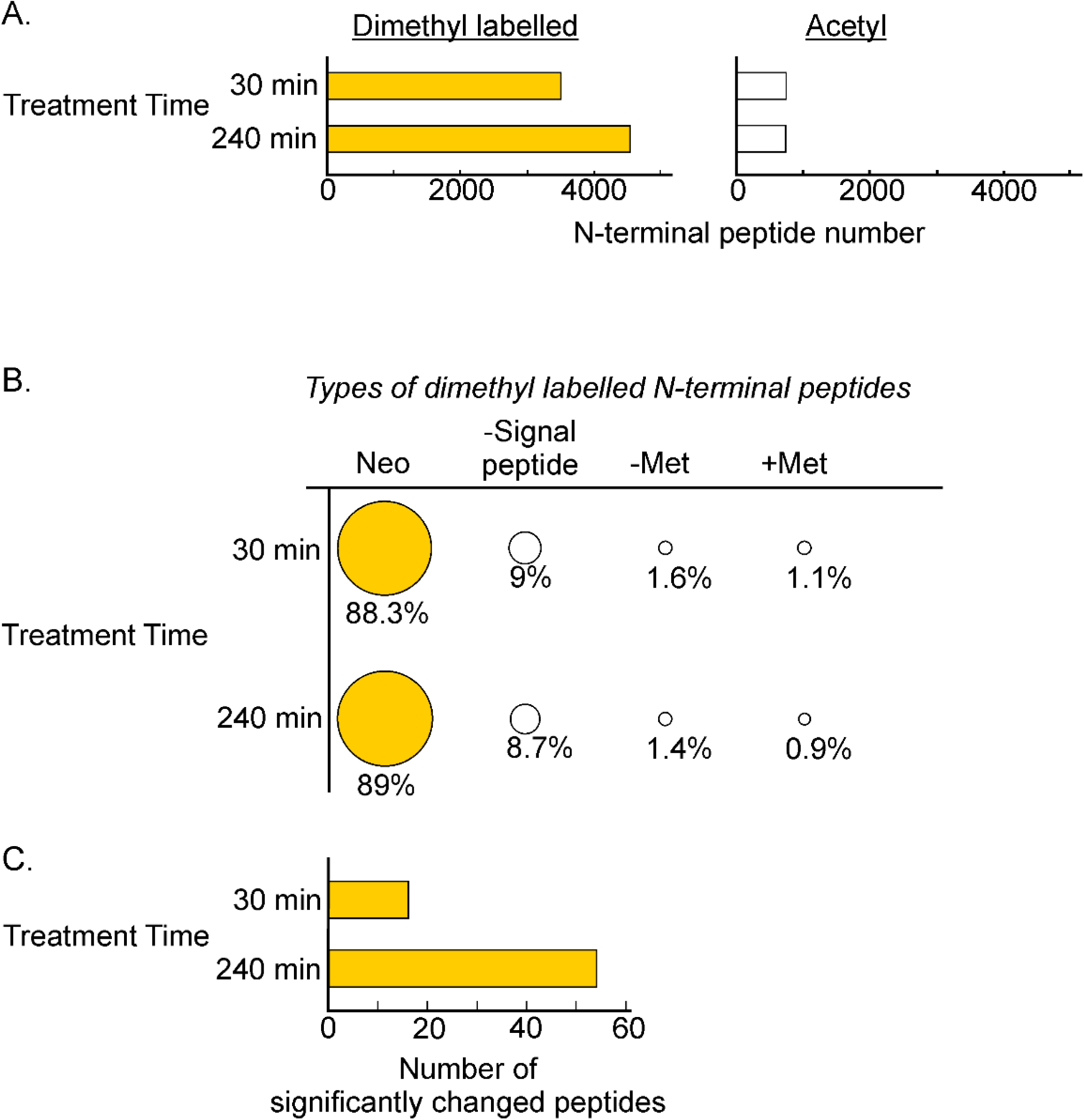
Proportion of the different types of N-terminal peptides identified from neurons co-treated with glutamate and calpeptin, related to Figure 6. **(A)** 4,260 and 5,303 neuronal N-terminal peptides, respectively were identified with high confidence (FDR ≤ 1.0%) from neurons co-treated with glutamate and calpeptin at 30 min and 240 min. Among them are the acetyl N-terminal peptides and those with free N-termini, which were isotopically dimethyl labelled in the TAILS procedures. **(B)** The different types of dimethyl labelled N-terminal peptides. Neo: neo-N-terminal peptides derived from truncated protein fragments of proteins undergoing limited proteolysis or intermediate peptide fragments of proteins undergoing degradation for clearance; -Signal peptide: N-terminal peptides derived from proteins with the signal peptide segment removed during biosynthesis; -Met: N-terminal peptides derived from proteins with the first methionine removed during biosynthesis; +Met: N-terminal peptides derived from proteins with the methionine encoded by the start codon intact. **(C)** Among the neo- N-terminal peptides, only 16 and 54 were derived from proteins undergoing significantly enhanced proteolysis at 30 min and 240 min of the co-treatment, respectively.

**Figure S10.**
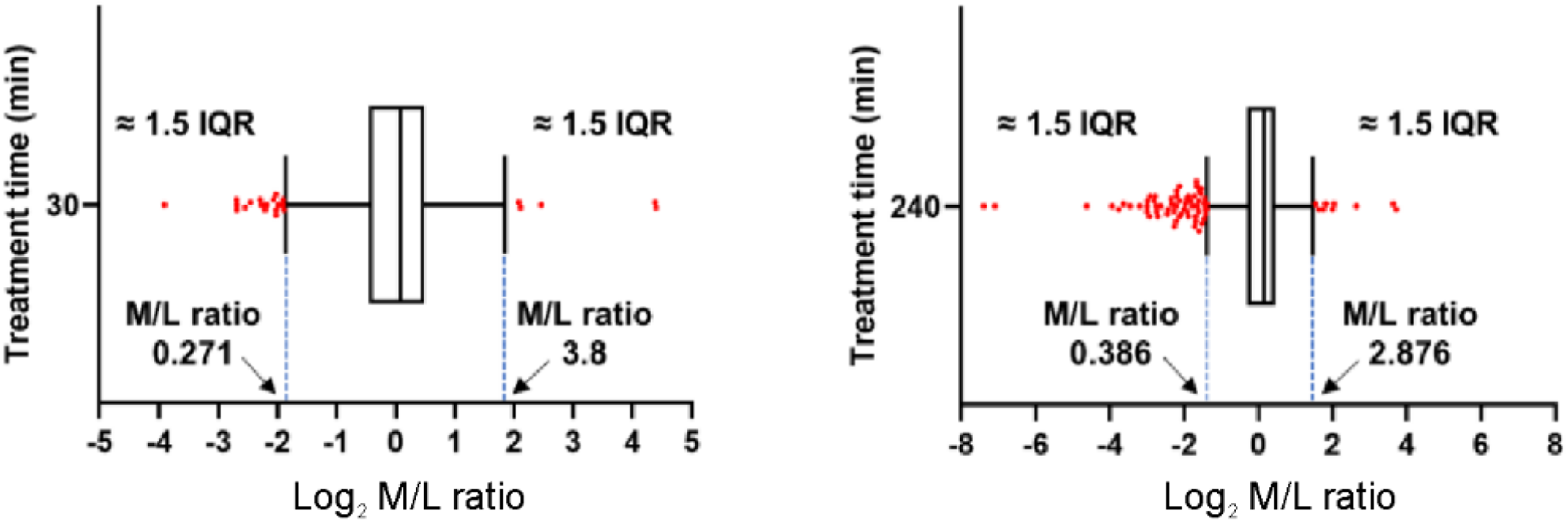
Determination of the abundance ratio cut-off values for classification of neo-N-terminal peptides into those generated by significantly enhanced degradation or significantly enhanced proteolytic processing in neurons co-treated with glutamate and calpeptin, related to Figure 6. Box-and-whisker plots of the Log_2_-nomalized abundance (M/L) ratios of the neo-N-terminal peptides exhibiting significant changes in abundance at 30 min (*left panel*) and 240 min (*right panel*) of the co-treatment. The M/L ratio cut-off values equivalent to 1.5 interquartile range (IQR) were calculated. For the N-terminal peptides co-treated for 30 min, the lower and upper cut-off values were 0.271 and 3.8 (i.e. M/L ≥ 3.8 for the neo-N- terminal peptides generated by significantly enhanced proteolytic processing and M/L ≤ 0.271 for those generated by significantly enhanced degradation). For those derived from neurons co-treated for 240 min, the lower and upper cut-off values were 0.386 and 2.876, respectively. Red dots represent the neo-N-terminal peptides with M/L ratios ≥ the upper cut-off values or ≤ the lower cut-off values.

**Figure S11.**
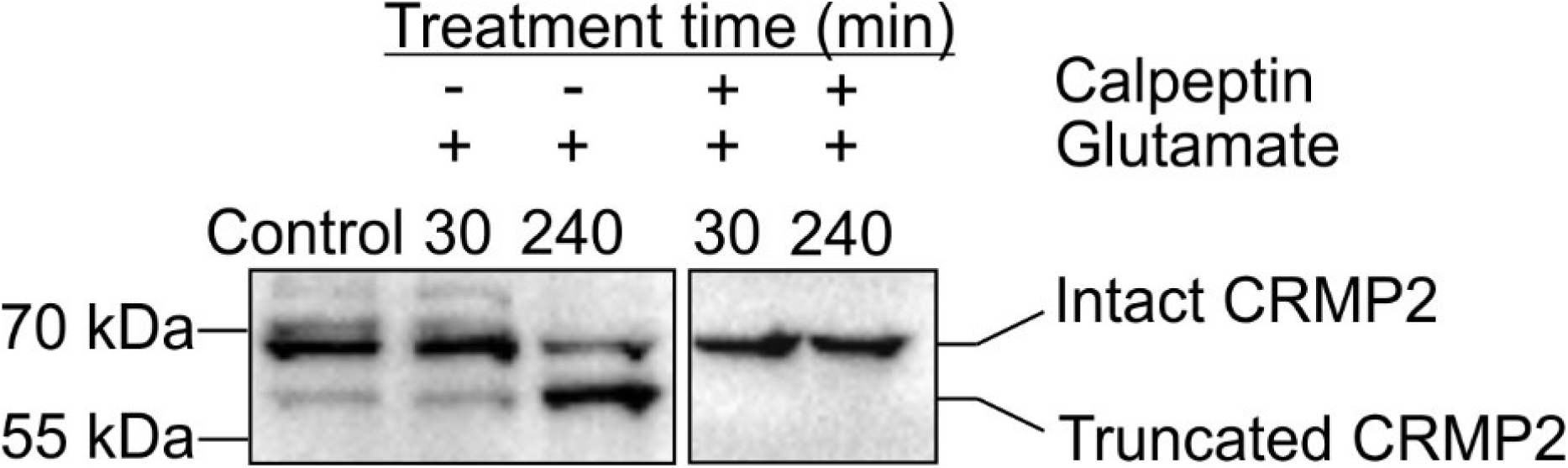
Calpeptin abolished proteolytic processing of CRMP2 to form truncated fragments induced by glutamate over-simulation, related to Figures 5 and 6. Western blots of lysates of neurons treated with glutamate and neurons co-treated with glutamate and calpeptin probed with the anti-CRMP2 antibody. The image of lysates of control neurons and neurons treated with glutamate is also shown in Figure 5D.

**Figure S12.**
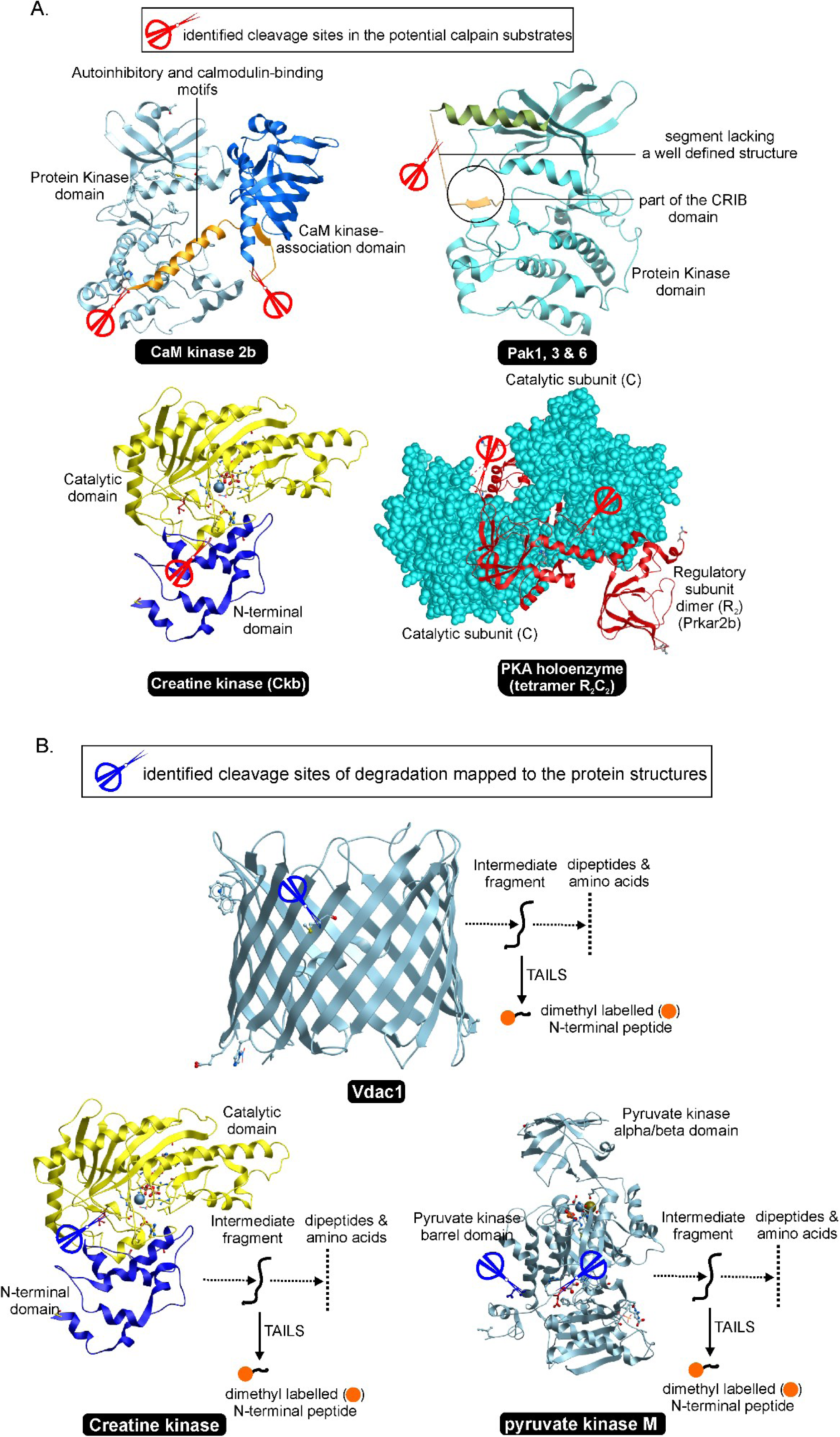
Cleavage sites identified in selected proteins in neurons during excitotoxicity, related to Figure 6. **(A)** Proteins undergoing enhanced proteolytic processing catalyzed by calpains. Calmodulin-dependent protein kinase 2b (CaM kinase 2b), Creatine kinase, p21-activated kinases 1, 3 and 6 (PAK1, 3, & 6) and the regulatory subunit (R_2_) of cAMP-dependent protein kinase (Prkar2b) were identified by the TAILS method to underwent significantly enhanced proteolytic processing during excitotoxicity. The ribbon model representations of their three dimensional structures are depicted. The red scissors indicate the locations of the identified cleavage sites. The PDB accession codes of these structures are: CaM kinase 2b, 3SOA; PAK1,3 & 6, 3tnp (this is the structure of PAK4, which shows a high degree of sequence homology with PAK1, 3 and 6); creatine kinase, 3b6r; holoenzyme of cAMP-dependent protein kinase (PKA) with Prkar2b as the regulatory subunits (R_2_) and two catalytic subunits (C subunit) with the protein kinase domain, 3tnp. All cleavage sites in these proteins are located outside a functional domain. **(B)** Proteins undergoing enhanced degradation. Voltage-dependent anion-selective channel protein 1 (Vdac1), Creatine kinase and pyruvate kinase M were identified by the TAILS method as neuronal proteins undergoing significantly enhanced degradation for clearance in excitotoxicity. The red scissors indicate the locations of the identified cleavage sites. As these sites are within the properly folded proteins or functional domains, the identified neo-N-terminal peptides were likely generated by enhanced cleavage of the intermediate fragments in neurons during excitotoxicity. The PDB accession codes of these structures are: Vdac1, 6g6u; creatine kinase, 3b6r; pyruvate kinase M, 3srf.

**Figure S13.**
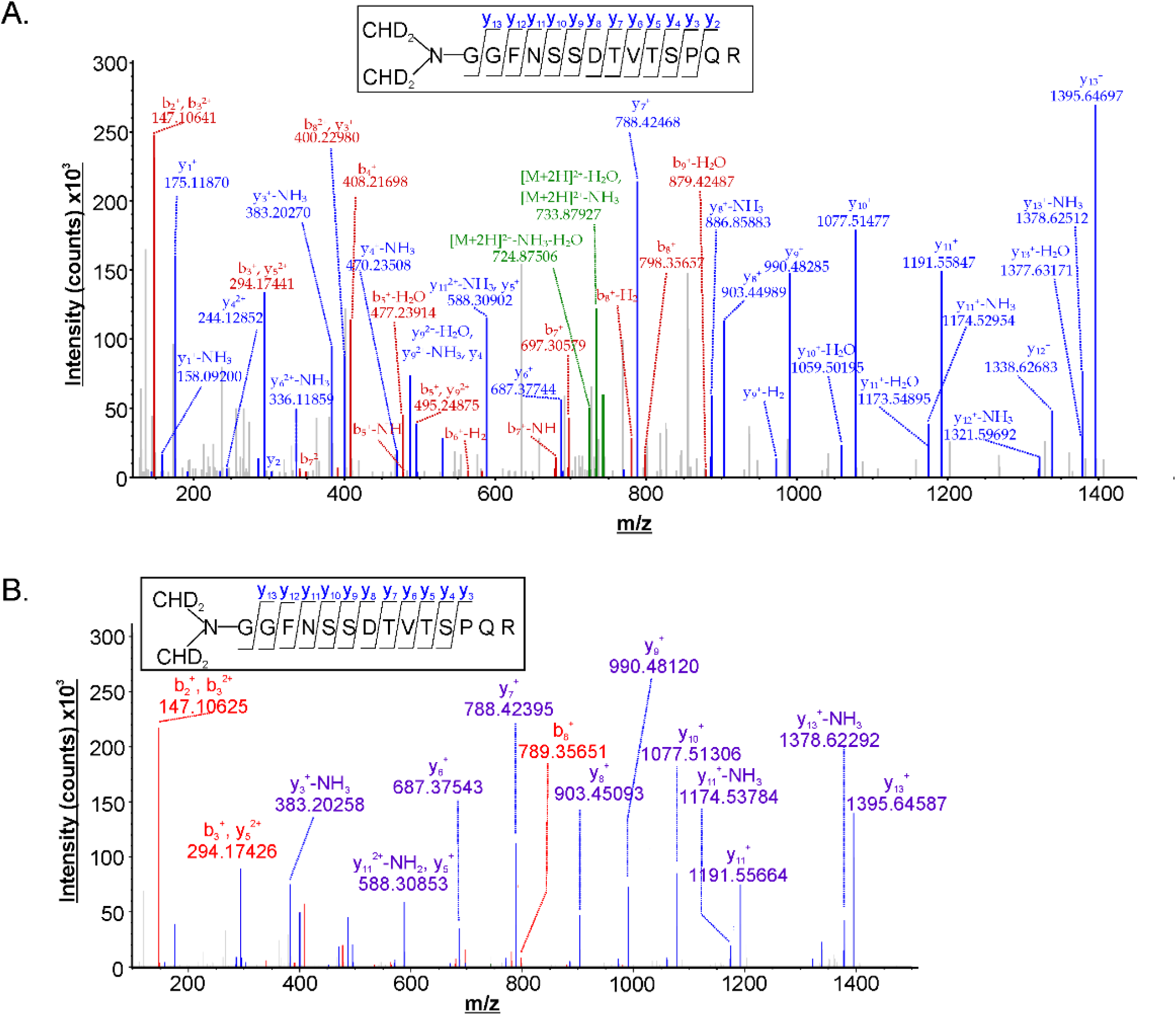
Identification of the neo-N-terminal peptide derived from the truncated Src fragments in excitotoxic neurons and in the reaction of *in vitro* cleavage of recombinant Src (R-Src) by calpain 1, related to Figures 6 and 7. **(A)** Fragment ion chromatogram identifying a neo-N-terminal peptide encompassing residues 54 to 77 of neuronal Src (inset) detectable exclusively in neurons treated with glutamate for 30 min. **(B)** The fragment ion chromatogram identifying the deuterated dimethyl-labelled Src (64-77) segment of R-Src (inset) as the neo-N-terminal peptide detected only in the reaction mixture containing R-Src and calpain 1 at 120 min of incubation. The insets show the confirmed amino acid sequence. Blue: y ions, Red: b ions.

**Figure S14.**
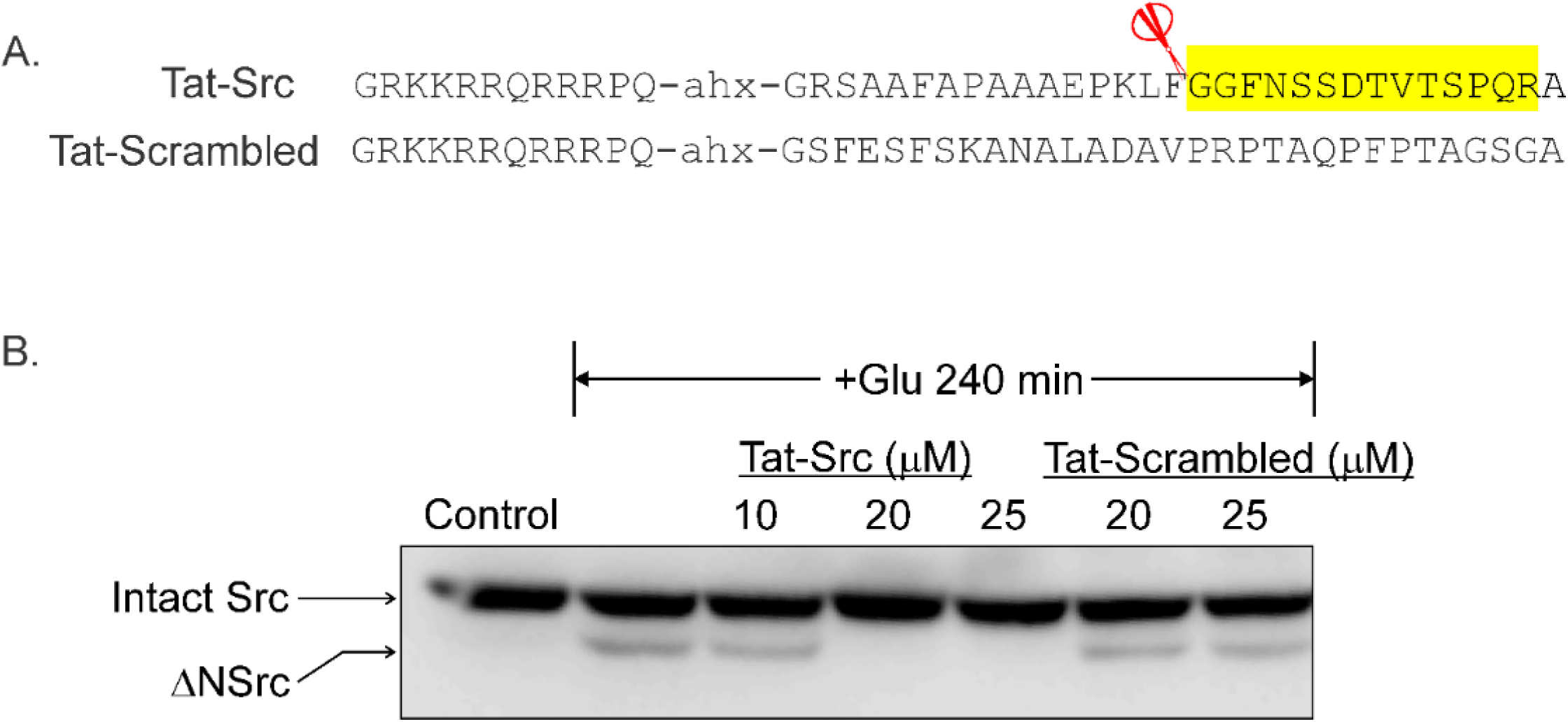
Tat-SRC but not Tat-Scrambled blocks cleavage of Src in neurons, related to Figures 6D, 6E and 7. **(A)** Sequences of Tat-Src and Tat-scrambled-Src. Segment highlighted in yellow: the neo-N-terminal peptide corresponding to Src(64-77) detected exclusively in glutamate-treated neurons (Figure 13A). Red scissor: the cleavage site in neuronal Src targeted by the excitotoxicity-activated proteases in neurons and by calpain 1 *in vitro*. **(B)** Tat-Src or Tat-Scrambled of the designated concentrations were added to the culture medium 1 h prior to treatment of cultured neurons with 100 μM glutamate. Cleavage of Src during excitotoxicity was monitored by anti-Src Western blot. ΔNSrc: truncated Src fragment.

## Notes

### Competing Interest Statement

The authors have declared no competing interest.

## REFERENCES

Aarts, M., Liu, Y., Liu, L., Besshoh, S., Arundine, M., Gurd, J.W., Wang, Y.T., Salter, M.W., and Tymianski, M. (2002). Treatment of ischemic brain damage by perturbing NMDA receptor-PSD-95 protein interactions. Science 298, 846–850.

Amata, I., Maffei, M., and Pons, M. (2014). Phosphorylation of unique domains of Src family kinases. Front Genet 5, 181.

Arbesu, M., Maffei, M., Cordeiro, T.N., Teixeira, J.M., Perez, Y., Bernado, P., Roche, S., and Pons, M. (2017). The Unique Domain Forms a Fuzzy Intramolecular Complex in Src Family Kinases. Structure 25, 630–640 e634.

Arthur, P.G., Matich, G.P., Pang, W.W., Yu, D.Y., and Bogoyevitch, M.A. (2007). Necrotic death of neurons following an excitotoxic insult is prevented by a peptide inhibitor of c-jun N-terminal kinase. J Neurochem 102, 65–76.

Badugu, R., Garcia, M., Bondada, V., Joshi, A., and Geddes, J.W. (2008). N terminus of calpain 1 is a mitochondrial targeting sequence. J Biol Chem 283, 3409–3417.

Baudry, M., and Bi, X. (2016). Calpain-1 and Calpain-2: The Yin and Yang of Synaptic Plasticity and Neurodegeneration. Trends Neurosci 39, 235–245.

Bevers, M.B., and Neumar, R.W. (2008). Mechanistic role of calpains in postischemic neurodegeneration. J Cereb Blood Flow Metab 28, 655–673.

Bian, Y., Li, L., Dong, M., Liu, X., Kaneko, T., Cheng, K., Liu, H., Voss, C., Cao, X., Wang, Y., et al. (2016). Ultra-deep tyrosine phosphoproteomics enabled by a phosphotyrosine superbinder. Nat Chem Biol 12, 959–966.

Boczek, E.E., Luo, Q., Dehling, M., Ropke, M., Mader, S.L., Seidl, A., Kaila, V.R.I., and Buchner, J. (2019). Autophosphorylation activates c-Src kinase through global structural rearrangements. J Biol Chem 294, 13186–13197.

Brennan-Minnella, A.M., Shen, Y., El-Benna, J., and Swanson, R.A. (2013). Phosphoinositide 3-kinase couples NMDA receptors to superoxide release in excitotoxic neuronal death. Cell Death Dis 4, e580.

Brennan, A.M., Suh, S.W., Won, S.J., Narasimhan, P., Kauppinen, T.M., Lee, H., Edling, Y., Chan, P.H., and Swanson, R.A. (2009). NADPH oxidase is the primary source of superoxide induced by NMDA receptor activation. Nat Neurosci 12, 857–863.

Brorson, J.R., Marcuccilli, C.J., and Miller, R.J. (1995). Delayed antagonism of calpain reduces excitotoxicity in cultured neurons. Stroke 26, 1259–1266; discussion 1267.

Burgess, H.A., and Reiner, O. (2001). Cleavage of doublecortin-like kinase by calpain releases an active kinase fragment from a microtubule anchorage domain. J Biol Chem 276, 36397–36403.

Chan, P.H. (2001). Reactive oxygen radicals in signaling and damage in the ischemic brain. J Cereb Blood Flow Metab 21, 2–14.

Chen, S., Murphy, J., Toth, R., Campbell, D.G., Morrice, N.A., and Mackintosh, C. (2008). Complementary regulation of TBC1D1 and AS160 by growth factors, insulin and AMPK activators. Biochem J 409, 449–459.

Chen, Y., Brennan-Minnella, A.M., Sheth, S., El-Benna, J., and Swanson, R.A. (2015). Tat-NR2B9c prevents excitotoxic neuronal superoxide production. J Cereb Blood Flow Metab 35, 739–742.

Chen, Z., Lei, C., Wang, C., Li, N., Srivastava, M., Tang, M., Zhang, H., Choi, J.M., Jung, S.Y., Qin, J., et al. (2019). Global phosphoproteomic analysis reveals ARMC10 as an AMPK substrate that regulates mitochondrial dynamics. Nat Commun 10, 104.

Choi, D.W. (1988). Glutamate neurotoxicity and diseases of the nervous system. Neuron 1, 623–634.

Choi, D.W., Maulucci-Gedde, M., and Kriegstein, A.R. (1987). Glutamate neurotoxicity in cortical cell culture. J Neurosci 7, 357–368.

Cook, D.J., Teves, L., and Tymianski, M. (2012). A translational paradigm for the preclinical evaluation of the stroke neuroprotectant Tat-NR2B9c in gyrencephalic nonhuman primates. Sci Transl Med 4, 154ra133.

Crooks, G.E., Hon, G., Chandonia, J.M., and Brenner, S.E. (2004). WebLogo: a sequence logo generator. Genome Res 14, 1188–1190.

Cuerrier, D., Moldoveanu, T., and Davies, P.L. (2005). Determination of peptide substrate specificity for mu-calpain by a peptide library-based approach: the importance of primed side interactions. J Biol Chem 280, 40632–40641.

Cuesto, G., Enriquez-Barreto, L., Carames, C., Cantarero, M., Gasull, X., Sandi, C., Ferrus, A., Acebes, A., and Morales, M. (2011). Phosphoinositide-3-kinase activation controls synaptogenesis and spinogenesis in hippocampal neurons. J Neurosci 31, 2721–2733.

D’Orsi, B., Bonner, H., Tuffy, L.P., Dussmann, H., Woods, I., Courtney, M.J., Ward, M.W., and Prehn, J.H. (2012). Calpains are downstream effectors of bax-dependent excitotoxic apoptosis. J Neurosci 32, 1847–1858.

Ding, X., Liu, S., Tian, M., Zhang, W., Zhu, T., Li, D., Wu, J., Deng, H., Jia, Y., Xie, W., et al. (2017). Activity-induced histone modifications govern Neurexin-1 mRNA splicing and memory preservation. Nat Neurosci 20, 690–699.

Dudek, H., Datta, S.R., Franke, T.F., Birnbaum, M.J., Yao, R., Cooper, G.M., Segal, R.A., Kaplan, D.R., and Greenberg, M.E. (1997). Regulation of neuronal survival by the serine-threonine protein kinase Akt. Science 275, 661–665.

duVerle, D.A., and Mamitsuka, H. (2019). CalCleaveMKL: a Tool for Calpain Cleavage Prediction. Methods Mol Biol 1915, 121–147.

El-Gebali, S., Mistry, J., Bateman, A., Eddy, S.R., Luciani, A., Potter, S.C., Qureshi, M., Richardson, L.J., Salazar, G.A., Smart, A., et al. (2019). The Pfam protein families database in 2019. Nucleic Acids Res 47, D427–D432.

Fraser, S.A., Davies, M., Katerelos, M., Gleich, K., Choy, S.W., Steel, R., Galic, S., Mount, P.F., Kemp, B.E., and Power, D.A. (2014). Activation of AMPK reduces the co-transporter activity of NKCC1. Mol Membr Biol 31, 95–102.

Fricker, M., Tolkovsky, A.M., Borutaite, V., Coleman, M., and Brown, G.C. (2018). Neuronal Cell Death. Physiol Rev 98, 813–880.

Ginet, V., Spiehlmann, A., Rummel, C., Rudinskiy, N., Grishchuk, Y., Luthi-Carter, R., Clarke, P.G., Truttmann, A.C., and Puyal, J. (2014). Involvement of autophagy in hypoxic-excitotoxic neuronal death. Autophagy 10, 846–860.

Girouard, H., Wang, G., Gallo, E.F., Anrather, J., Zhou, P., Pickel, V.M., and Iadecola, C. (2009). NMDA receptor activation increases free radical production through nitric oxide and NOX2. J Neurosci 29, 2545–2552.

Gleeson, J.G., Allen, K.M., Fox, J.W., Lamperti, E.D., Berkovic, S., Scheffer, I., Cooper, E.C., Dobyns, W.B., Minnerath, S.R., Ross, M.E., et al. (1998). Doublecortin, a brain-specific gene mutated in human X-linked lissencephaly and double cortex syndrome, encodes a putative signaling protein. Cell 92, 63–72.

Gleeson, J.G., Lin, P.T., Flanagan, L.A., and Walsh, C.A. (1999). Doublecortin is a microtubule-associated protein and is expressed widely by migrating neurons. Neuron 23, 257–271.

Greenwood, S.M., Mizielinska, S.M., Frenguelli, B.G., Harvey, J., and Connolly, C.N. (2007). Mitochondrial dysfunction and dendritic beading during neuronal toxicity. J Biol Chem 282, 26235–26244.

Haass, C., and Mandelkow, E. (2010). Fyn-tau-amyloid: a toxic triad. Cell 142, 356–358.

Hancock, W.S., and Battersby, J.E. (1976). A new micro-test for the detection of incomplete coupling reactions in solid-phase peptide synthesis using 2,4,6-trinitrobenzenesulphonic acid. Anal Biochem 71, 260–264.

Hardingham, G.E., Fukunaga, Y., and Bading, H. (2002). Extrasynaptic NMDARs oppose synaptic NMDARs by triggering CREB shut-off and cell death pathways. Nat Neurosci 5, 405–414.

Hasbani, M.J., Schlief, M.L., Fisher, D.A., and Goldberg, M.P. (2001). Dendritic spines lost during glutamate receptor activation reemerge at original sites of synaptic contact. J Neurosci 21, 2393–2403.

Herrero-Martin, G., Hoyer-Hansen, M., Garcia-Garcia, C., Fumarola, C., Farkas, T., Lopez-Rivas, A., and Jaattela, M. (2009). TAK1 activates AMPK-dependent cytoprotective autophagy in TRAIL-treated epithelial cells. EMBO J 28, 677–685.

Hetman, M., and Gozdz, A. (2004). Role of extracellular signal regulated kinases 1 and 2 in neuronal survival. Eur J Biochem 271, 2050–2055.

Hill, M.D., Goyal, M., Menon, B.K., Nogueira, R.G., McTaggart, R.A., Demchuk, A.M., Poppe, A.Y., Buck, B.H., Field, T.S., Dowlatshahi, D., et al. (2020). Efficacy and safety of nerinetide for the treatment of acute ischaemic stroke (ESCAPE-NA1): a multicentre, double-blind, randomised controlled trial. Lancet 395, 878–887.

Hill, M.D., Martin, R.H., Mikulis, D., Wong, J.H., Silver, F.L., Terbrugge, K.G., Milot, G., Clark, W.M., Macdonald, R.L., Kelly, M.E., et al. (2012). Safety and efficacy of NA-1 in patients with iatrogenic stroke after endovascular aneurysm repair (ENACT): a phase 2, randomised, double-blind, placebo-controlled trial. Lancet Neurol 11, 942–950.

Hofmann, K., and Falquet, L. (2001). A ubiquitin-interacting motif conserved in components of the proteasomal and lysosomal protein degradation systems. Trends in biochemical sciences 26, 347–350.

Hoque, A., Williamson, N.A., Ameen, S.S., Ciccotosto, G.D., Hossain, M.I., Oakhill, J.S., Ng, D.C.H., Ang, C.S., and Cheng, H.C. (2019). Quantitative proteomic analyses of dynamic signalling events in cortical neurons undergoing excitotoxic cell death. Cell Death Dis 10, 213.

Hornbeck, P.V., Zhang, B., Murray, B., Kornhauser, J.M., Latham, V., and Skrzypek, E. (2015). PhosphoSitePlus, 2014: mutations, PTMs and recalibrations. Nucleic Acids Res 43, D512–520.

Hosie, K.A., King, A.E., Blizzard, C.A., Vickers, J.C., and Dickson, T.C. (2012). Chronic excitotoxin-induced axon degeneration in a compartmented neuronal culture model. ASN Neuro 4.

Hossain, I.M., Hoque, A., Lessene, G., Aizuddin Kamaruddin, M., Chu, P.W., Ng, I.H., Irtegun, S., Ng, D.C., Bogoyevitch, M.A., Burgess, A.W., et al. (2015). Dual role of Src kinase in governing neuronal survival. Brain Res 1594, 1–14.

Hossain, M.I., Roulston, C.L., Kamaruddin, M.A., Chu, P.W., Ng, D.C., Dusting, G.J., Bjorge, J.D., Williamson, N.A., Fujita, D.J., Cheung, S.N., et al. (2013). A truncated fragment of Src protein kinase generated by calpain-mediated cleavage is a mediator of neuronal death in excitotoxicity. J Biol Chem 288, 9696–9709.

Inagaki, N., Chihara, K., Arimura, N., Menager, C., Kawano, Y., Matsuo, N., Nishimura, T., Amano, M., and Kaibuchi, K. (2001). CRMP-2 induces axons in cultured hippocampal neurons. Nature neuroscience 4, 781–782.

Iqbal Hossain, M., Hoque, A., Lessene, G., Aizuddin Kamaruddin, M., Chu, P.W., Ng, I.H., Irtegun, S., Ng, D.C., Bogoyevitch, M.A., Burgess, A.W., et al. (2015). Dual role of Src kinase in governing neuronal survival. Brain Res 1594, 1–14.

Kall, L., Canterbury, J.D., Weston, J., Noble, W.S., and MacCoss, M.J. (2007). Semi-supervised learning for peptide identification from shotgun proteomics datasets. Nat Methods 4, 923–925.

Kaufman, A.M., Milnerwood, A.J., Sepers, M.D., Coquinco, A., She, K., Wang, L., Lee, H., Craig, A.M., Cynader, M., and Raymond, L.A. (2012). Opposing Roles of Synaptic and Extrasynaptic NMDA Receptor Signaling in Cocultured Striatal and Cortical Neurons. J Neurosci 32, 3992–4003.

Kleifeld, O., Doucet, A., auf dem Keller, U., Prudova, A., Schilling, O., Kainthan, R.K., Starr, A.E., Foster, L.J., Kizhakkedathu, J.N., and Overall, C.M. (2010). Isotopic labeling of terminal amines in complex samples identifies protein N-termini and protease cleavage products. Nat Biotechnol 28, 281–288.

Kleifeld, O., Doucet, A., Prudova, A., auf dem Keller, U., Gioia, M., Kizhakkedathu, J.N., and Overall, C.M. (2011). Identifying and quantifying proteolytic events and the natural N terminome by terminal amine isotopic labeling of substrates. Nat Protoc 6, 1578–1611.

Kondo, S., Takahashi, K., Kinoshita, Y., Nagai, J., Wakatsuki, S., Araki, T., Goshima, Y., and Ohshima, T. (2019). Genetic inhibition of CRMP2 phosphorylation at serine 522 promotes axonal regeneration after optic nerve injury. Sci Rep 9, 7188.

Kramer, A., Green, J., Pollard, J., Jr., and Tugendreich, S. (2014). Causal analysis approaches in Ingenuity Pathway Analysis. Bioinformatics 30, 523–530.

Lankiewicz, S., Marc Luetjens, C., Truc Bui, N., Krohn, A.J., Poppe, M., Cole, G.M., Saido, T.C., and Prehn, J.H. (2000). Activation of calpain I converts excitotoxic neuron death into a caspase-independent cell death. J Biol Chem 275, 17064–17071.

Li, X., Wilmanns, M., Thornton, J., and Kohn, M. (2013). Elucidating human phosphatase-substrate networks. Sci Signal 6, rs10.

Liebl, M.P., and Hoppe, T. (2016). It’s all about talking: two-way communication between proteasomal and lysosomal degradation pathways via ubiquitin. American journal of physiology Cell physiology 311, C166–178.

Lipton, S.A. (2007). Pathologically activated therapeutics for neuroprotection. Nat Rev Neurosci 8, 803–808.

Liu, D.Z., Waldau, B., Ander, B.P., Zhan, X., Stamova, B., Jickling, G.C., Lyeth, B.G., and Sharp, F.R. (2017). Inhibition of Src family kinases improves cognitive function after intraventricular hemorrhage or intraventricular thrombin. J Cereb Blood Flow Metab 37, 2359–2367.

Liu, J., Liu, M.C., and Wang, K.K. (2008). Physiological and pathological actions of calpains in glutamatergic neurons. Sci Signal 1, tr3.

Liu, Z., Cao, J., Gao, X., Ma, Q., Ren, J., and Xue, Y. (2011). GPS-CCD: a novel computational program for the prediction of calpain cleavage sites. PLoS One 6, e19001.

Liu, Z.X., Yu, K., Dong, J., Zhao, L., Liu, Z., Zhang, Q., Li, S., Du, Y., and Cheng, H. (2019). Precise Prediction of Calpain Cleavage Sites and Their Aberrance Caused by Mutations in Cancer. Front Genet 10, 715.

Ludwig, C., Gillet, L., Rosenberger, G., Amon, S., Collins, B.C., and Aebersold, R. (2018). Data-independent acquisition-based SWATH-MS for quantitative proteomics: a tutorial. Mol Syst Biol 14, e8126.

MacDermott, A.B., Mayer, M.L., Westbrook, G.L., Smith, S.J., and Barker, J.L. (1986). NMDA-receptor activation increases cytoplasmic calcium concentration in cultured spinal cord neurones. Nature 321, 519–522.

Miller, C.J., and Turk, B.E. (2018). Homing in: Mechanisms of Substrate Targeting by Protein Kinases. Trends Biochem Sci 43, 380–394.

Moldoveanu, T., Campbell, R.L., Cuerrier, D., and Davies, P.L. (2004). Crystal structures of calpain-E64 and -leupeptin inhibitor complexes reveal mobile loops gating the active site. J Mol Biol 343, 1313–1326.

Morita, T., and Sobue, K. (2009). Specification of neuronal polarity regulated by local translation of CRMP2 and Tau via the mTOR-p70S6K pathway. J Biol Chem 284, 27734–27745.

Nakamura, T., Tu, S., Akhtar, M.W., Sunico, C.R., Okamoto, S., and Lipton, S.A. (2013). Aberrant protein s-nitrosylation in neurodegenerative diseases. Neuron 78, 596–614.

Nawabi, H., Belin, S., Cartoni, R., Williams, P.R., Wang, C., Latremoliere, A., Wang, X., Zhu, J., Taub, D.G., Fu, X., et al. (2015). Doublecortin-Like Kinases Promote Neuronal Survival and Induce Growth Cone Reformation via Distinct Mechanisms. Neuron 88, 704–719.

Niwa, S., Nakamura, F., Tomabechi, Y., Aoki, M., Shigematsu, H., Matsumoto, T., Yamagata, A., Fukai, S., Hirokawa, N., Goshima, Y., et al. (2017). Structural basis for CRMP2-induced axonal microtubule formation. Sci Rep 7, 10681.

Okada, M., Nada, S., Yamanashi, Y., Yamamoto, T., and Nakagawa, H. (1991). CSK: a protein-tyrosine kinase involved in regulation of src family kinases. J Biol Chem 266, 24249–24252.

Olney, J.W. (1969). Brain lesions, obesity, and other disturbances in mice treated with monosodium glutamate. Science 164, 719–721.

Osborne, K.A., Shigeno, T., Balarsky, A.M., Ford, I., McCulloch, J., Teasdale, G.M., and Graham, D.I. (1987). Quantitative assessment of early brain damage in a rat model of focal cerebral ischaemia. J Neurol Neurosurg Psychiatry 50, 402–410.

Patel, O., Dai, W., Mentzel, M., Griffin, M.D., Serindoux, J., Gay, Y., Fischer, S., Sterle, S., Kropp, A., Burns, C.J., et al. (2016). Biochemical and Structural Insights into Doublecortin-like Kinase Domain 1. Structure 24, 1550–1561.

Perez-Riverol, Y., Csordas, A., Bai, J., Bernal-Llinares, M., Hewapathirana, S., Kundu, D.J., Inuganti, A., Griss, J., Mayer, G., Eisenacher, M., et al. (2019). The PRIDE database and related tools and resources in 2019: improving support for quantification data. Nucleic Acids Res 47, D442–D450.

Perez, Y., Gairi, M., Pons, M., and Bernado, P. (2009). Structural characterization of the natively unfolded N-terminal domain of human c-Src kinase: insights into the role of phosphorylation of the unique domain. J Mol Biol 391, 136–148.

Perfetto, L., Briganti, L., Calderone, A., Cerquone Perpetuini, A., Iannuccelli, M., Langone, F., Licata, L., Marinkovic, M., Mattioni, A., Pavlidou, T., et al. (2016). SIGNOR: a database of causal relationships between biological entities. Nucleic Acids Res 44, D548–554.

Raaijmakers, L.M., Giansanti, P., Possik, P.A., Mueller, J., Peeper, D.S., Heck, A.J., and Altelaar, A.F. (2015). PhosphoPath: Visualization of Phosphosite-centric Dynamics in Temporal Molecular Networks. J Proteome Res 14, 4332–4341.

Reiner, O., Coquelle, F.M., Peter, B., Levy, T., Kaplan, A., Sapir, T., Orr, I., Barkai, N., Eichele, G., and Bergmann, S. (2006). The evolving doublecortin (DCX) superfamily. BMC genomics 7, 188.

Sashindranath, M., Daglas, M., and Medcalf, R.L. (2015). Evaluation of gait impairment in mice subjected to craniotomy and traumatic brain injury. Behav Brain Res 286, 33–38.

Sashindranath, M., Samson, A.L., Downes, C.E., Crack, P.J., Lawrence, A.J., Li, Q.X., Ng, A.Q., Jones, N.C., Farrugia, J.J., Abdella, E., et al. (2011). Compartment- and context-specific changes in tissue-type plasminogen activator (tPA) activity following brain injury and pharmacological stimulation. Lab Invest 91, 1079–1091.

Sattler, R., Xiong, Z., Lu, W.Y., Hafner, M., MacDonald, J.F., and Tymianski, M. (1999). Specific coupling of NMDA receptor activation to nitric oxide neurotoxicity by PSD-95 protein. Science 284, 1845–1848.

Schaar, B.T., Kinoshita, K., and McConnell, S.K. (2004). Doublecortin microtubule affinity is regulated by a balance of kinase and phosphatase activity at the leading edge of migrating neurons. Neuron 41, 203–213.

Schaffer, B.E., Levin, R.S., Hertz, N.T., Maures, T.J., Schoof, M.L., Hollstein, P.E., Benayoun, B.A., Banko, M.R., Shaw, R.J., Shokat, K.M., et al. (2015). Identification of AMPK Phosphorylation Sites Reveals a Network of Proteins Involved in Cell Invasion and Facilitates Large-Scale Substrate Prediction. Cell Metab 22, 907–921.

Schoch, K.M., Evans, H.N., Brelsfoard, J.M., Madathil, S.K., Takano, J., Saido, T.C., and Saatman, K.E. (2012). Calpastatin overexpression limits calpain-mediated proteolysis and behavioral deficits following traumatic brain injury. Exp Neurol 236, 371–382.

Scholzke, M.N., Potrovita, I., Subramaniam, S., Prinz, S., and Schwaninger, M. (2003). Glutamate activates NF-kappaB through calpain in neurons. Eur J Neurosci 18, 3305–3310.

Shen, C.H., Yuan, P., Perez-Lorenzo, R., Zhang, Y., Lee, S.X., Ou, Y., Asara, J.M., Cantley, L.C., and Zheng, B. (2013). Phosphorylation of BRAF by AMPK impairs BRAF-KSR1 association and cell proliferation. Mol Cell 52, 161–172.

Shinkai-Ouchi, F., Koyama, S., Ono, Y., Hata, S., Ojima, K., Shindo, M., duVerle, D., Ueno, M., Kitamura, F., Doi, N., et al. (2016). Predictions of Cleavability of Calpain Proteolysis by Quantitative Structure-Activity Relationship Analysis Using Newly Determined Cleavage Sites and Catalytic Efficiencies of an Oligopeptide Array. Mol Cell Proteomics 15, 1262–1280.

Shu, T., Tseng, H.C., Sapir, T., Stern, P., Zhou, Y., Sanada, K., Fischer, A., Coquelle, F.M., Reiner, O., and Tsai, L.H. (2006). Doublecortin-like kinase controls neurogenesis by regulating mitotic spindles and M phase progression. Neuron 49, 25–39.

Simon, R.P., Swan, J.H., Griffiths, T., and Meldrum, B.S. (1984). Blockade of N-methyl-D-aspartate receptors may protect against ischemic damage in the brain. Science 226, 850–852.

Song, J., Wang, Y., Li, F., Akutsu, T., Rawlings, N.D., Webb, G.I., and Chou, K.C. (2019). iProt-Sub: a comprehensive package for accurately mapping and predicting protease-specific substrates and cleavage sites. Brief Bioinform 20, 638–658.

Sorimachi, H., Hata, S., and Ono, Y. (2011). Calpain chronicle--an enzyme family under multidisciplinary characterization. Proc Jpn Acad Ser B Phys Biol Sci 87, 287–327.

Sumi, T., Imasaki, T., Aoki, M., Sakai, N., Nitta, E., Shirouzu, M., and Nitta, R. (2018). Structural Insights into the Altering Function of CRMP2 by Phosphorylation. Cell Struct Funct 43, 15–23.

Sun, H.S., Doucette, T.A., Liu, Y., Fang, Y., Teves, L., Aarts, M., Ryan, C.L., Bernard, P.B., Lau, A., Forder, J.P., et al. (2008). Effectiveness of PSD95 inhibitors in permanent and transient focal ischemia in the rat. Stroke 39, 2544–2553.

Takano, J., Tomioka, M., Tsubuki, S., Higuchi, M., Iwata, N., Itohara, S., Maki, M., and Saido, T.C. (2005). Calpain mediates excitotoxic DNA fragmentation via mitochondrial pathways in adult brains: evidence from calpastatin mutant mice. J Biol Chem 280, 16175–16184.

Taus, T., Kocher, T., Pichler, P., Paschke, C., Schmidt, A., Henrich, C., and Mechtler, K. (2011). Universal and confident phosphorylation site localization using phosphoRS. J Proteome Res 10, 5354–5362.

Thingholm, T.E., Jensen, O.N., Robinson, P.J., and Larsen, M.R. (2008). SIMAC (sequential elution from IMAC), a phosphoproteomics strategy for the rapid separation of monophosphorylated from multiply phosphorylated peptides. Mol Cell Proteomics 7, 661–671.

Tominaga, K., Nakanishi, H., Yasuda, Y., and Yamamoto, K. (1998). Excitotoxin-induced neuronal death is associated with response of a unique intracellular aspartic proteinase, cathepsin E. J Neurochem 71, 2574–2584.

Tompa, P., Buzder-Lantos, P., Tantos, A., Farkas, A., Szilagyi, A., Banoczi, Z., Hudecz, F., and Friedrich, P. (2004). On the sequential determinants of calpain cleavage. J Biol Chem 279, 20775–20785.

Tukey, J.W. (1977). Exploratory data analysis, Addison-Wesley.

Tyanova, S., and Cox, J. (2018). Perseus: A Bioinformatics Platform for Integrative Analysis of Proteomics Data in Cancer Research. Methods Mol Biol 1711, 133–148.

Tyanova, S., Temu, T., Sinitcyn, P., Carlson, A., Hein, M.Y., Geiger, T., Mann, M., and Cox, J. (2016). The Perseus computational platform for comprehensive analysis of (prote)omics data. Nat Methods 13, 731–740.

Uchida, Y., Ohshima, T., Sasaki, Y., Suzuki, H., Yanai, S., Yamashita, N., Nakamura, F., Takei, K., Ihara, Y., Mikoshiba, K., et al. (2005). Semaphorin3A signalling is mediated via sequential Cdk5 and GSK3beta phosphorylation of CRMP2: implication of common phosphorylating mechanism underlying axon guidance and Alzheimer’s disease. Genes Cells 10, 165–179.

Wakatsuki, S., Saitoh, F., and Araki, T. (2011). ZNRF1 promotes Wallerian degeneration by degrading AKT to induce GSK3B-dependent CRMP2 phosphorylation. Nat Cell Biol 13, 1415–1423.

Wang, K.K., Nath, R., Posner, A., Raser, K.J., Buroker-Kilgore, M., Hajimohammadreza, I., Probert, A.W., Jr., Marcoux, F.W., Ye, Q., Takano, E., et al. (1996). An alpha-mercaptoacrylic acid derivative is a selective nonpeptide cell-permeable calpain inhibitor and is neuroprotective. Proc Natl Acad Sci U S A 93, 6687–6692.

Wang, T., Ma, G., Ang, C.S., Korhonen, P.K., Stroehlein, A.J., Young, N.D., Hofmann, A., Chang, B.C.H., Williamson, N.A., and Gasser, R.B. (2020). The developmental phosphoproteome of Haemonchus contortus. J Proteomics 213, 103615.

Wang, W., Zhang, F., Li, L., Tang, F., Siedlak, S.L., Fujioka, H., Liu, Y., Su, B., Pi, Y., and Wang, X. (2015). MFN2 couples glutamate excitotoxicity and mitochondrial dysfunction in motor neurons. J Biol Chem 290, 168–182.

Wang, Y., Briz, V., Chishti, A., Bi, X., and Baudry, M. (2013). Distinct roles for mu-calpain and m-calpain in synaptic NMDAR-mediated neuroprotection and extrasynaptic NMDAR-mediated neurodegeneration. J Neurosci 33, 18880–18892.

Wilson, S.M., Ki Yeon, S., Yang, X.F., Park, K.D., and Khanna, R. (2014). Differential regulation of collapsin response mediator protein 2 (CRMP2) phosphorylation by GSK3ss and CDK5 following traumatic brain injury. Front Cell Neurosci 8, 135.

Woods, A., Dickerson, K., Heath, R., Hong, S.P., Momcilovic, M., Johnstone, S.R., Carlson, M., and Carling, D. (2005). Ca2+/calmodulin-dependent protein kinase kinase-beta acts upstream of AMP-activated protein kinase in mammalian cells. Cell Metab 2, 21–33.

Woods, A., Johnstone, S.R., Dickerson, K., Leiper, F.C., Fryer, L.G., Neumann, D., Schlattner, U., Wallimann, T., Carlson, M., and Carling, D. (2003). LKB1 is the upstream kinase in the AMP-activated protein kinase cascade. Curr Biol 13, 2004–2008.

Xu, J., Kurup, P., Zhang, Y., Goebel-Goody, S.M., Wu, P.H., Hawasli, A.H., Baum, M.L., Bibb, J.A., and Lombroso, P.J. (2009). Extrasynaptic NMDA receptors couple preferentially to excitotoxicity via calpain-mediated cleavage of STEP. J Neurosci 29, 9330–9343.

Xu, W., Wong, T.P., Chery, N., Gaertner, T., Wang, Y.T., and Baudry, M. (2007). Calpain-mediated mGluR1alpha truncation: a key step in excitotoxicity. Neuron 53, 399–412.

Xue, L., Wang, P., Cao, P., Zhu, J.K., and Tao, W.A. (2014). Identification of extracellular signal-regulated kinase 1 (ERK1) direct substrates using stable isotope labeled kinase assay-linked phosphoproteomics. Mol Cell Proteomics 13, 3199–3210.

Yamada, K.H., Kozlowski, D.A., Seidl, S.E., Lance, S., Wieschhaus, A.J., Sundivakkam, P., Tiruppathi, C., Chishti, I., Herman, I.M., Kuchay, S.M., et al. (2012). Targeted gene inactivation of calpain-1 suppresses cortical degeneration due to traumatic brain injury and neuronal apoptosis induced by oxidative stress. J Biol Chem 287, 13182–13193.

Yamashima, T., Kohda, Y., Tsuchiya, K., Ueno, T., Yamashita, J., Yoshioka, T., and Kominami, E. (1998). Inhibition of ischaemic hippocampal neuronal death in primates with cathepsin B inhibitor CA-074: a novel strategy for neuroprotection based on ’calpain-cathepsin hypothesis’. Eur J Neurosci 10, 1723–1733.

Yoshimura, T., Kawano, Y., Arimura, N., Kawabata, S., Kikuchi, A., and Kaibuchi, K. (2005). GSK-3beta regulates phosphorylation of CRMP-2 and neuronal polarity. Cell 120, 137–149.

Yu, W., Fantl, W.J., Harrowe, G., and Williams, L.T. (1998). Regulation of the MAP kinase pathway by mammalian Ksr through direct interaction with MEK and ERK. Curr Biol 8, 56–64.

Yuasa-Kawada, J., Suzuki, R., Kano, F., Ohkawara, T., Murata, M., and Noda, M. (2003). Axonal morphogenesis controlled by antagonistic roles of two CRMP subtypes in microtubule organization. The European journal of neuroscience 17, 2329–2343.

Zhang, Z., Ottens, A.K., Sadasivan, S., Kobeissy, F.H., Fang, T., Hayes, R.L., and Wang, K.K. (2007). Calpain-mediated collapsin response mediator protein-1, -2, and -4 proteolysis after neurotoxic and traumatic brain injury. J Neurotrauma 24, 460–472.

